# Indigenous plants promote insect biodiversity in urban greenspaces

**DOI:** 10.1101/2020.05.29.122572

**Authors:** Luis Mata, Alan N. Andersen, Alejandra Morán-Ordóñez, Amy K. Hahs, Anna Backstrom, Christopher D. Ives, Daniel Bickel, David Duncan, Estibaliz Palma, Freya Thomas, Kate Cranney, Ken Walker, Ian Shears, Linda Semeraro, Mallik Malipatil, Melinda L. Moir, Michaela Plein, Nick Porch, Peter A. Vesk, Tessa R. Smith, Yvonne Lynch

## Abstract

The contribution of urban greenspaces to support biodiversity and provide benefits for people is increasingly recognised. However, ongoing management practices still favour (1) vegetation oversimplification, often limiting greenspaces to lawns and tree canopy rather than multi-layered vegetation that includes under and midstorey; and (2) the use of nonnative plant species. These practices likely hinder the potential of greenspaces to sustain indigenous biodiversity, particularly for taxa like insects, that rely on plants for food and habitat. Yet, little is known about which plant species may maximise positive outcomes for taxonomically and functionally diverse insect communities in urban greenspaces. Additionally, while urban environments are expected to experience high rates of introductions, quantitative assessments of the relative occupancy of indigenous vs. introduced insect species in greenspace are rare – hindering understanding of how greenspace management may promote indigenous biodiversity while limiting the establishment of introduced insects. Using a hierarchically replicated study design across 15 public parks, we recorded occurrence data from 552 insect species on 133 plant species – differing in planting design element (lawn, midstorey and tree canopy), midstorey growth form (forbs, lilioids, graminoids and shrubs) and origin (nonnative, native and indigenous) – to assess: (1) the relative contributions of indigenous and introduced insect species and (2) which plant species sustained the highest number of indigenous insects. Our data indicates that the insect community was predominately composed of indigenous rather than introduced species. Our findings further highlight the core role of multi-layered vegetation in sustaining high insect biodiversity in urban areas, with indigenous midstorey and canopy representing key elements to maintain rich and functionally diverse indigenous insect communities. Intriguingly, graminoids supported the highest indigenous insect richness across all studied growth forms by plant origin groups. Taken together, our study emphasise the opportunity posed by indigenous understory and midstorey plants, particularly indigenous graminoids in our study area, to promote indigenous insect biodiversity in urban greenspaces. Our work provides a blueprint and stimulus for built-environment professionals to incorporate into their practice plant species palettes that foster a larger presence of indigenous over regionally native or nonnative plant species, whilst incorporating a broader mixture of midstorey growth forms.

## Introduction

Urban greenspaces provide well-documented benefits for biodiversity and people. Remnant bushland, parks, gardens, golf courses, greenroofs, pop-up parks and other types of greenspace support a great diversity of microbial, fungal, plant and animal species (Madre et al. 2013, Aronson et al. 2014, Baldock et al. 2015, Beninde et al. 2015, McGregor-Fors et al. 2016, Mata et al. 2017, Threlfall et al. 2017, Baldock et al. 2019, Mata et al. 2019), and provide a diverse array of health, mental, cognitive, social, cultural and spiritual benefits for people who interact with them (Keniger et al. 2013, Dadvand et al. 2015, Hartig and Kahn 2016, Flies et al. 2017, Maller et al. 2018, Lai et al. 2019, Mata et al. 2020). Hence, researchers, practitioners, built-environment professionals and policymakers are increasingly working together to promote the positive socio-ecological outcomes of greenspaces (Aronson et al. 2017, Lepczyk et al. 2017, Nilon et al. 2017, Parris et al. 2018, Soanes et al. 2019). Further, the importance of greenspaces has been recently highlighted at an international policy level with the United Nations’ New Urban Agenda committing to “promoting the creation and maintenance of well-connected and well-distributed networks of open, multipurpose, safe, inclusive, accessible, green and quality public spaces” (United Nations 2017).

An ubiquitous practice that can hinder the potential of greenspaces to support biodiversity is the oversimplification of vegetation structure (Le Roux et al. 2014, Threlfall et al. 2016), which has led many greenspaces to be vegetated by only two planting design elements: lawn and tree canopy (Ignatieva et al. 2015, Aronson et al. 2017). In contrast, greenspaces with a more complex, multi-layered vertical structure – those including understorey and midstorey vegetation (henceforth midstorey for brevity) – provide positive outcomes for a taxonomically and functionally diverse range of taxa (Beninde et al. 2015, Mata et al. 2017, Threlfall et al. 2017, Majewska and Altizer 2020). Unlike the lawn and tree canopy, the midstorey is a heterogenous mix of different plant growth forms, including forbs, graminoids, lilioids and shrubs amongst others. Yet, at present, the combination of plant species and growth forms that maximise positive outcomes for non-plant species in the midstorey remains poorly understood.

An additional issue that limits biodiversity in urban areas is that most non-remnant greenspaces, particularly intensively manicured ones such as residential gardens and public parks, are composed predominately of nonnative plant species (Threlfall et al. 2016). Nonnative plants are rarely well-suited to provide resources for indigenous primary consumers (e.g. herbivorous insects and frugivorous birds), nor to indigenous secondary and apex consumers (e.g. predatory and parasitoid insects and insectivorous birds and bats) that depend on primary consumers as food resources (Ballard et al. 2013, Burghardt and Tallamy 2013, Ikin et al. 2013, Salisbury et al. 2015, Threlfall et al. 2017). These studies highlight how management practices that promote the use of nonnative plants are likely to reduce the capacity of greenspaces to sustain diverse communities of indigenous biodiversity.

In most of the studies to date, plant origin has been treated as an aggregate plot-level explanatory variable (e.g. treatment plot as either nonnative or native; plot nativeness as percentage cover of native vegetation). A focus on plant species rather than plot as the unit of analysis allows for a more nuanced understanding of how plant origin may influence the capacity of plants to provide resources for associated consumer species. Moreover, focusing on the plant species level may advance understanding of how plant origin interacts with other plant-level attributes — such as planting design element and growth form — to produce positive outcomes for consumer species, particularly of taxa that rely extensively on plants for food and habitat resources, such as insects.

Insects are a particularly important component of urban biodiversity (Sattler et al. 2011, New 2015) and the ecological functions they perform provide numerous benefits to urban residents (Prather et al. 2013, Benett and Lovell 2014, Baldock et al. 2015), along with some disbenefits (Dunn 2010, Rust and Su 2012). Plants and insects have often coevolved in close association with each other, with many insect species showing high levels of specialisation (Forister et al. 2012). Plants also provide food, foraging, nesting, oviposition, shelter and overwintering resources to insect detritivores, predators and parasitoids; indeed, practices that promote these resources in agroecosystems by fostering plant diversity and structural complexity are key components of pest management strategies (Landis et al. 2000). The availability of suitable host plants within greenspaces is therefore a key determinant of insect diversity in urban environments (Aronson et al. 2016). However, as far as we are aware no study has sought to identify specific plant species of differing origins and growth forms that promote taxonomically and functionally diverse insect communities in urban greenspaces.

Urban environments are central nodes of human-mediated dispersal networks (Bullock et al. 2018) and hotspots of novel resource utilisation (Valentine et al. 2020), and introduced species are therefore often prominent (Cadotte et al. 2017, Paap et al. 2017). The number of introduced insect species can be relatively low (Madre et al. 2013, Mata et al. 2017), but they often occur in high abundance and this is especially the case for generalist bees and butterflies (Matteson & Langellotto 2010, Threlfall et al. 2015). However, as far as we are aware, no quantitative assessments of the relative contributions of indigenous and introduced species in greenspace insect communities have been reported in the literature. This paucity of data hinders understanding of how greenspaces may be managed to promote indigenous insect biodiversity while limiting the establishment of introduced species.

Here we use a plant-insect metanetwork dataset collected across 15 greenspaces within a densely urbanised inner-city municipality to assess: (1) the relative contributions of indigenous and introduced species in insect communities of urban environments, and (2) which plant species should be planted to support indigenous insect species. Th project followed the science-government partnerships model (Ives and Lynch 2014) – an approach that advocates for industry professionals and researchers to work in close association to guarantee that theoretically interesting and practically important questions are identified. This ensured that the implications of our research findings could be applied as practical on-ground actions that were embedded into a new business as usual at the City of Melbourne. We began by determining what proportion of insect species occurring in the greenspaces are indigenous, and whether indigenous insect species are more common than introduced species. We then examined how the richness of indigenous insect species varies with planting design element (lawn, midstorey and tree canopy), midstorey growth form (forbs, lilioids, graminoids and shrubs) and plant origin groups (nonnative, native and indigenous). Finally, we grouped plants species according to a combination of planting design element by plant origin (Fig. 1a), and midstorey growth form by plant origin (Fig. 1b) to assess how the richness of indigenous insect species vary amongst these groups. We do this assessment for the whole insect community and for five key functional groups: pollinators and other flower-visiting taxa, herbivores, predators, parasitoids and detritivores. We also examined variation amongst the plant groups in insect species composition and number of unique species.

**Figure 1.**
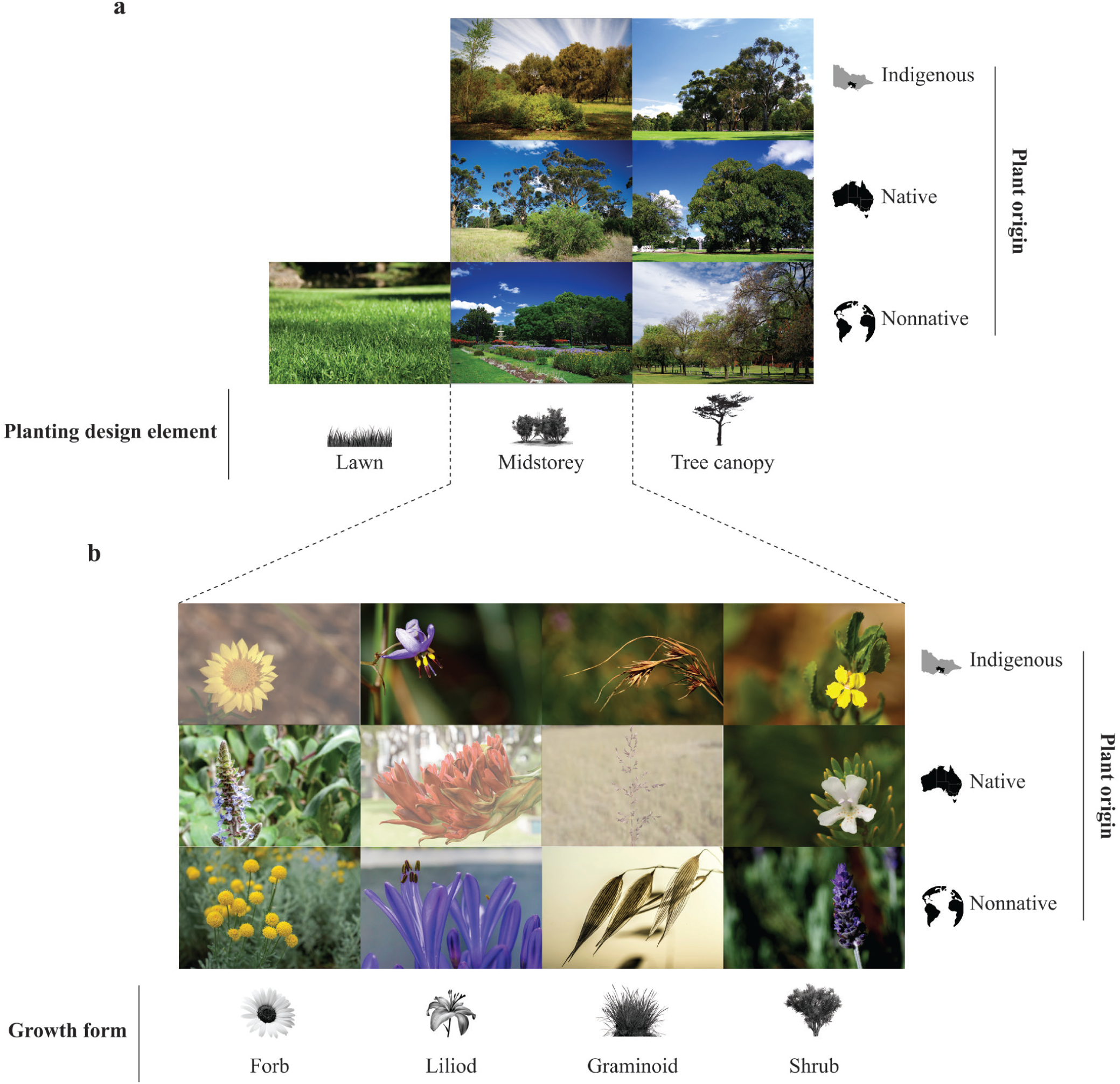
Visual representation of the planting design element by plant origin (a) and midstorey growth form by plant origin (b) groups that were part of the theoretical and empirical dimensions of this study. Dimmed boxes in (b) indicate theoretical combinations of midstorey growth form by plant origin groups that did/may not occur in the study area or that occur but not at surveyable densities within the study’s plots.

## Methods

### Study design

The study was conducted in the City of Melbourne, Australia. This is one of 31 municipalities within Greater Melbourne, a large metropolitan area spanning approximately 10,000 km^2^ within the Urban Growth Boundary and home to over five million people (Victorian State Revenue Office: https://www.sro.vic.gov.au/greater-melbourne-map-and-urban-zones). The City of Melbourne covers 37.7 km^2^ and incorporates a central business district, transport and distribution hubs. Approximately 13% of the land area is covered by vegetated open space, including grassy woodlands, wetlands, estuaries and a greenspace network of parks, gardens and streetscapes. The municipality is home to approximately 180,000 residents and receives approximately 900,000 daily visitors (City of Melbourne: https://www.melbourne.vic.gov.au/). Our study was conducted across 15 public parks, which varied in size across four orders of magnitude (1.1 × 10^3^ m^2^ – 1.3 × 10^6^ m^2^; Appendix 1: Table S1; Appendix 1: Fig. S1a). We established a total of 130 plots across the 15 parks, with the number of plots in each park (2-36), and their size (84-148 m^2^), varying according to the park’s area (Appendix S1: Eq. S1-S3) and planting design elements and midstorey growth forms present (Appendix 1: Table S1; Appendix 1: Fig. S1b). Within each plot we identified all plants, totalling 133 species, genera or species complexes across the study area (Appendix S2: Table S2).

We classified plant species by origin as indigenous (n=30), regionally native (n=9) and nonnative (n=94). We define indigenous plant species – also referred to in the literature as locally native – as those that are native to the local bioregions. For this study, indigenous plant species are those that occurred before European settlement in the Volcanic Victorian Plain and Gippsland Plain bioregions, (State of Victoria: https://www.environment.vic.gov.au). Regionally native (henceforth native) are species that are native to Australia but not to the local bioregions and have been anthropogenically introduced. Nonnative species are those that have been introduced to Australia.

We also classified plant species by planting design element (henceforth design element) as lawn complex (n=41), midstorey (n=67) and tree canopy (n=25) (Fig. 1a). Midstorey taxa were further stratified by growth form as forb (n=8), lilioid (n=13), graminoid (n=8) and shrub (n=38) (Fig. 1b). We define lawn complexes (henceforth lawns) as patches dominated by turf forming grasses (Poaceae) intermixed with one or more small, ruderal herbaceous species. Midstorey species included broad-leaved perennial and annual herbaceous plants (forbs); petaloid monocots in orders Liliales and Asparagales (lilioids); grasses, sedges and rushes of typically vertical habit with linear foliage and inconspicuous wind pollinated flowers (graminoids); and woody perennials with multiple stems and < 5 m in height (shrubs). Tree canopy species included single-stemmed woody plants > 5 m in height.

### Insect survey

We sampled plant species for 12 insect groups known to dominate insect communities on above-ground vegetation: ants, bees, beetles, cicadas, flies, heteropteran bugs, jumping plant lice, leaf- and treehoppers, parasitoid wasps, planthoppers, sawflies and stinging wasps. Samples were taken by direct observation and by sweeping with an entomological net. Observation time and sweeps per plant species were standardised as a proportion of the plant species’ volume within the plot (Appendix S1: Eq. S4-S5), with each plant species in each plot sampled three times from January (summer) to late March (autumn) 2015. Sampled insect specimens were processed in the laboratory and identified to species/morphospecies. We assigned these as (1) either indigenous to the studied bioregions and/or native to Australia (henceforth indigenous) or introduced to Australia, and (2) one or more of the following functional groups: pollinators and other flower-visiting taxa (henceforth pollinators), herbivores, predators, parasitoids and detritivores.

### Data analysis

#### Estimating insect occupancy and species richness per plant species

To assess the proportion of indigenous insect species occurring in the greenspaces, whether they were more common than introduced ones, and how their species richness varied amongst the studied single and combined plant groups, we analysed our data with a three-level hierarchical metacommunity occupancy model (Kéry and Royle 2016). Plant species was our unit of analysis for drawing inferences on insect species occupancy, and each repeated spatial (individuals of the same plant species sampled in different plots) and temporal (same individual of a given plant species sampled at different times) samplings constituted the unit of detection replication. We structured the model around three levels: one for species occupancy; a second for species detectability; and a third to treat the occupancy and detection parameters for each species as random effects (Kéry and Royle 2016). Specifically, we used a variation of the model described by Mata et al. (2017), in which we specified the occupancy level model as:

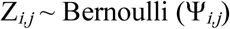

where Ψ_*i,j*_ is the probability that insect species *i* occurs at plant species *j*, and the detection level model as:

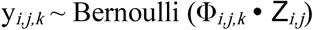

where Φ_*i,j,k*_ is the detection probability of insect species *i* at plant species *j* at spatiotemporal replicate *k*.

The occupancy and detection level linear predictors were specified on the logit-probability scale as:

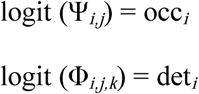

where occ_*i*_ and det_*i*_ are the species-specific random effects, which were specified as:

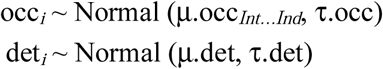

where the metacommunity mean occupancy hyperpriors for introduced and indigenous insect species, μ.occ_*Int*_ and μ.occ_*Ind*_, respectively, and the metacommunity mean detection hyperprior μ.det, were specified as Uniform (0, 1); and the metacommunity precision occupancy and detection hyperpriors, τ.occ and τ.det, respectively, were specified as Gamma (0.1, 0.1).

We then use the latent occurrence matrix Z_*ij*_ to estimate the insect species richness associated with each plant species SR_*j*_ through the summation:

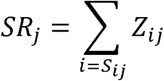

where *S*_*ij*_ is a ‘specificity’ vector indexing the insect species to be included in each plant species’ estimate. SR_*j*_ is then an estimate that accounts for plant-insect specificity, in which, for each plant species, the observed insect species are included with probability of occurrence = 1 and a limited random sub-sample of other insect species occurring in the study area are included with their 0 < *Z* < 1 estimated probabilities of occurrence. This allowed us to work within the reasonable ecological assumption that across the study area not every insect species will be associated with every co-occurring plant species. We conducted these estimations for the insect community as a whole but also independently for introduced and indigenous species. We estimated model parameters under Bayesian inference, using Markov Chain Monte Carlo (MCMC) simulations to draw samples from the parameters’ posterior distributions. As the species richness calculations were conducted within this modelling framework, we were able to derive the insect species per plant species estimates with their full associated uncertainties. This allowed us to average the species richness estimates of plant species belonging to the same group, and therefore obtain posterior distributions for each group that we could statistically compare. Our model was implemented in JAGS (Plummer 2003) and accessed through the R package *jagsUI* (Kellner 2016). We used three chains of 5,000 iterations, discarding the first 500 in each chain as burn-in. We visually inspected the MCMC chains and the values of the Gelman-Rubin statistic to verify acceptable convergence levels of R-hat < 1.1 (Gelman & Hill 2007).

#### Community dissimilarity

To determine whether the composition of insect species varied amongst the design elements/growth form by plant origin groups, we reorganised the data into insect-by-plant species matrices – cell values summarising the number of times a given insect species was sampled on a given plant species across its spatiotemporal replicates – and used these to calculate amongst-group community dissimilarity. Specifically, we used 1 - Jaccard similarity index as implemented in the R package *vegan* (Oksanen et al. 2016). We further used the data to create insect species lists for each group, which we partitioned into their corresponding Venn sets with the R package *VennDiagram* (Chen 2016). This allowed us to calculate the number of unique insect species – species found exclusively at a given plant group and not shared with any of the other groups – belonging to each group.

## Results

Our survey recorded 552 insect species, with the richest taxa being beetles (125 species), parasitoid wasps (121), flies (101), heteropteran bugs (61), leaf- and treehoppers (40) and jumping plant lice (31) (Appendix S2: Table S1). These represented 154 pollinator, 299 herbivore, 231 predator, 150 parasitoid and 231 detritivore species. The most commonly occurring species was the minute brown scavenger beetle *Cortinicara* sp. 1 (Latridiidae), an indigenous detritivore species that accounted for 12% of all records. The Argentine ant *Linepithema humile* was the most frequently occurring introduced species, accounting for approximately 3% of all records. Four new species were also discovered: one ant, one heteropteran bug and two jumping plant lice (Mata et al. 2015, 2016).

### Indigenous vs introduced insect species

There were approximately 30 times more indigenous (534) than introduced (18) insect species across the study area, with our model estimates indicating that any particular plant species was associated with 19 times more indigenous than introduced insect species (Fig. 2a; Appendix S2: Table S2). The mean number of introduced insect species found on individual plant species varied from zero to four; whereas the mean number of indigenous insect species varied from one to 109, with most plant species being associated with more than ten indigenous insect species (Fig. 2b; Appendix 2: Table S3; Appendix 2: Figure S1). The probability of occurrence of any particular insect species at a given plant across the study area was similarly low for introduced and indigenous species (Fig. 2c; Appendix 2: Table S4). The introduced insect fauna was represented by species showing moderate to very low (0.6 > Pocc > 0) species-specific probabilities of occurrence (Fig. 2d; Appendix 2: Table S1); whereas the species-specific probabilities of occurrence of indigenous insect species varied widely, with a few species showing very high occupancy levels (Pocc > 0.8) and most species showing low occupancy levels (Pocc < 0.4) (Fig. 2d; Appendix 2: Table S1). The probability of detecting any particular insect species at a given plant across the study area was similarly very low for introduced and indigenous species (Pdet < 0.03; Appendix S2: Table S4), and the species-specific probabilities of detection were consistently low for the large majority of insect species (Appendix S2: Table S1).

**Figure 2.**
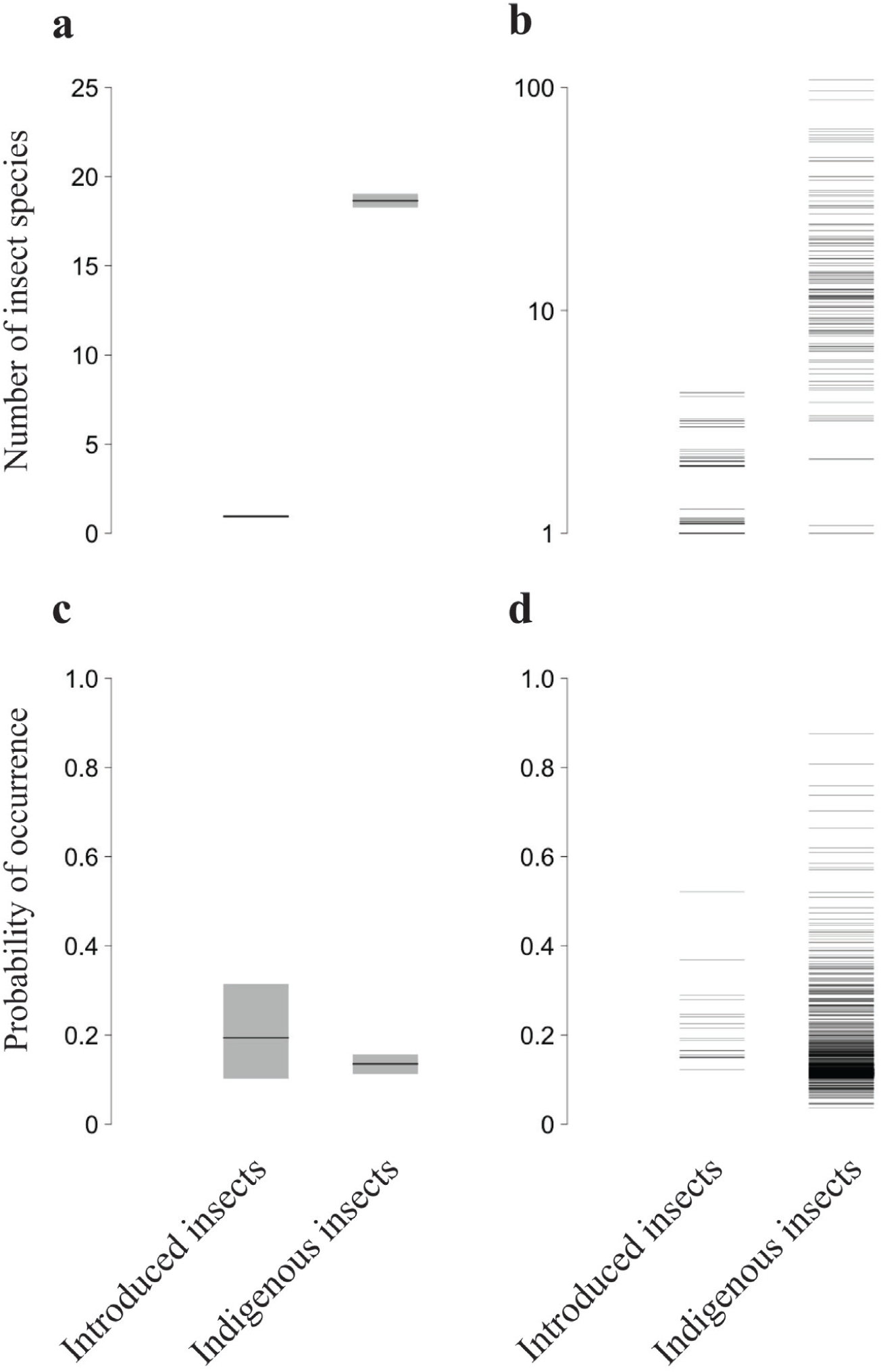
Estimated species richness of introduced and indigenous insect species at the average plant species across the study (a) and at each surveyed plant species (b). Estimated probabilities of occurrence of introduced and indigenous insect species at the average plant species across the study (c) and at each surveyed plant species (d). In all figures the black lines represent mean responses; the grey boxes in (a) and (c) represent the associated statistical uncertainty (95% Credible Intervals). Y-axis in (b) drawn in the log10 scale.

### Effect of design element, growth form and plant origin

Our model estimates indicate that all three design elements had different levels of indigenous insect species richness, with the average midstorey or tree canopy species showing approximately 2.5 times more insect species than the average lawn complex (Fig. 3a; Appendix S2: Table S5). Likewise, we found that the species richness of indigenous insects varied amongst midstorey growth forms, with the average graminoid species showing 2.9 times more insect species than the average lilioid, 2.4 times more than the average forb and 1.8 times more than the average shrub (Fig. 3b; Appendix S2: Table S5). Our estimates further indicate marked statistical differences with plant origin in the number of indigenous insect species (Appendix 2: Figure S1), with the average indigenous plant species showing 2.9 and 1.6 times more insect species than the average nonnative and native plant, respectively; and the average native plant showing 1.9 times more insect species than the average nonnative plant (Fig. 3c; Appendix S2: Table S5).

**Figure 3.**
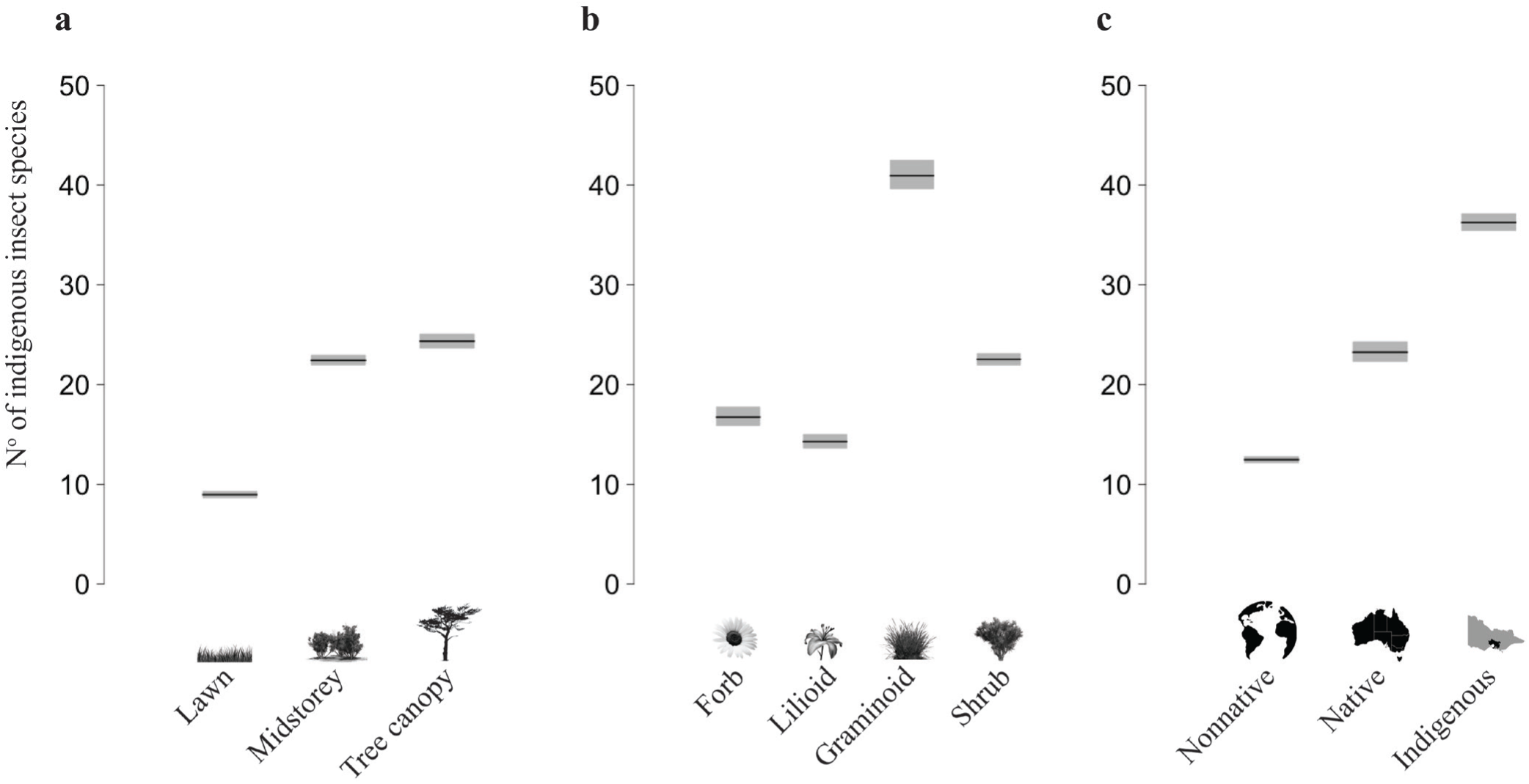
Estimated species richness of indigenous insects by planting design element (a), midstorey growth form (b) and plant origin (c). Black lines represent mean responses and grey boxes the associated statistical uncertainty (95% Credible Intervals).

### Combined effect of design element and plant origin

Our model estimates indicate that all design element by plant origin groups had different levels of indigenous insect species richness. In general, indigenous groups were associated with higher species richness than native and these with higher richness than nonnative. The single exception was native midstorey and nonnative tree canopy, which showed insect species richness levels that were not statistically different from each other (Fig. 4a; Appendix S2: Table S6). Overall, the indigenous midstorey was associated with the highest level of indigenous insect species richness, with the average plant species in this group showing 2.6 and 1.6 times more insect species than the average nonnative and native midstorey plant species, respectively (Fig. 4a; Appendix S2: Table S6). The indigenous tree canopy was associated with the second highest level of insect species richness, with the average plant species in this group showing 1.8 and 1.4 times more insect species than the average nonnative and native tree canopy plant species, respectively (Fig. 4a; Appendix S2: Table S6). Across insect functional groups, lawns showed the lowest insect species richness (Fig. 5a,c,e,g,i; Appendix S2: Table S6). The indigenous midstorey showed the highest number of indigenous insect species; however, for predators and detritivores the indigenous midstorey was not statistically different to the indigenous tree canopy (Fig. 5e,i; Appendix S2: Table S6). Other departures from the general pattern were observed for each functional group. The indigenous and native tree canopy groups did not show different levels of indigenous pollinators or parasitoid species (Fig. 5a,g; Appendix S2: Table S6), and the native tree canopy and midstorey groups did not show different levels of indigenous herbivore species (Fig. 5c; Appendix S2: Table S6). For predators, parasitoids and detritivores, the nonnative tree canopy group has a higher species richness than the native midstorey and was not statistically different than the native tree canopy – the native and nonnative midstorey groups in turn did not show different levels of associated species (Fig. 5e,g,i; Appendix S2: Table S6).

**Figure 4.**
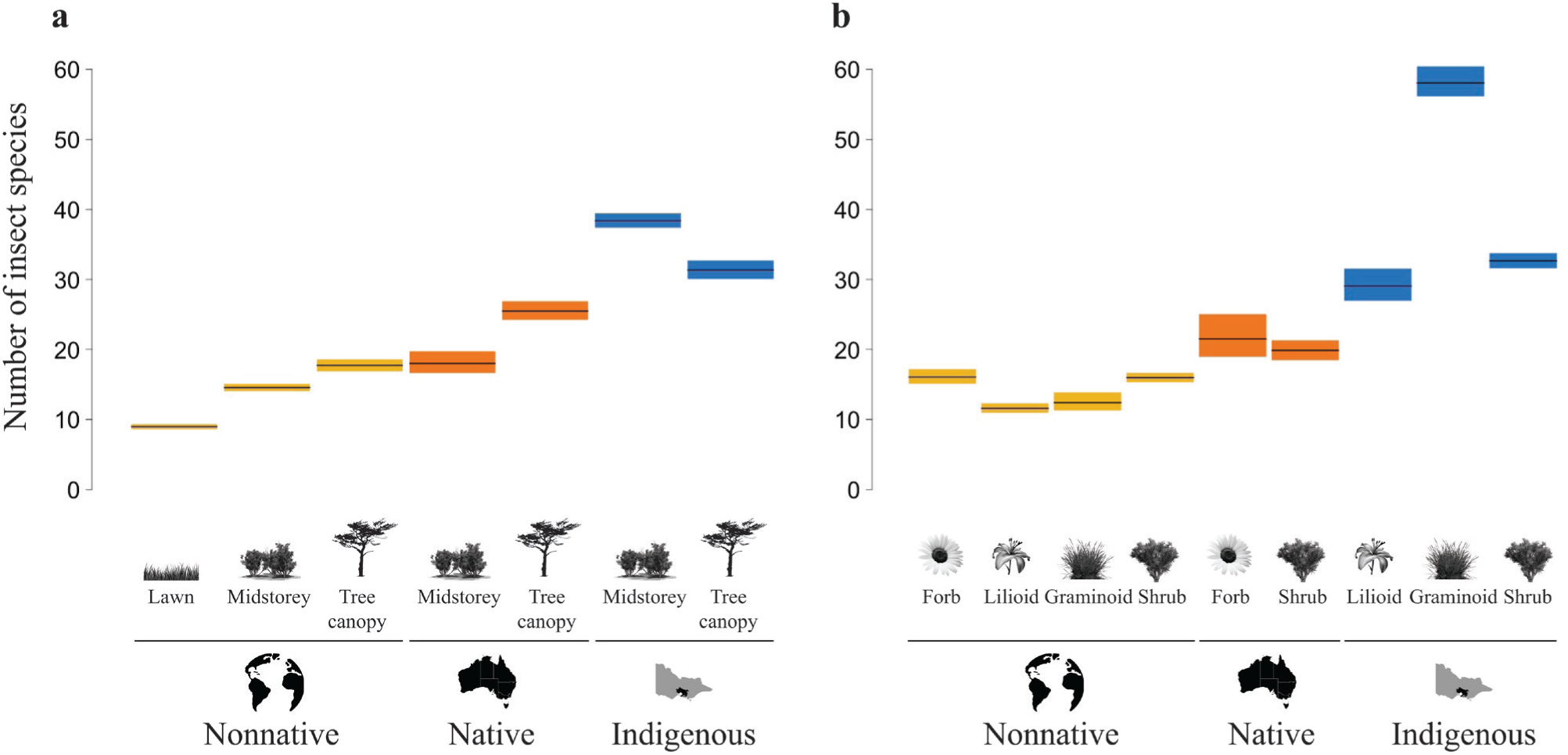
Estimated species richness of indigenous insect by planting design element by plant origin (a) and midstorey growth form by plant origin (b). Black lines represent mean responses and coloured boxes the associated statistical uncertainty (95% Credible Intervals). For ease of interpretation plant origin has been colour coded as yellow (nonnative), orange (native) or blue (indigenous).

**Figure 5.**
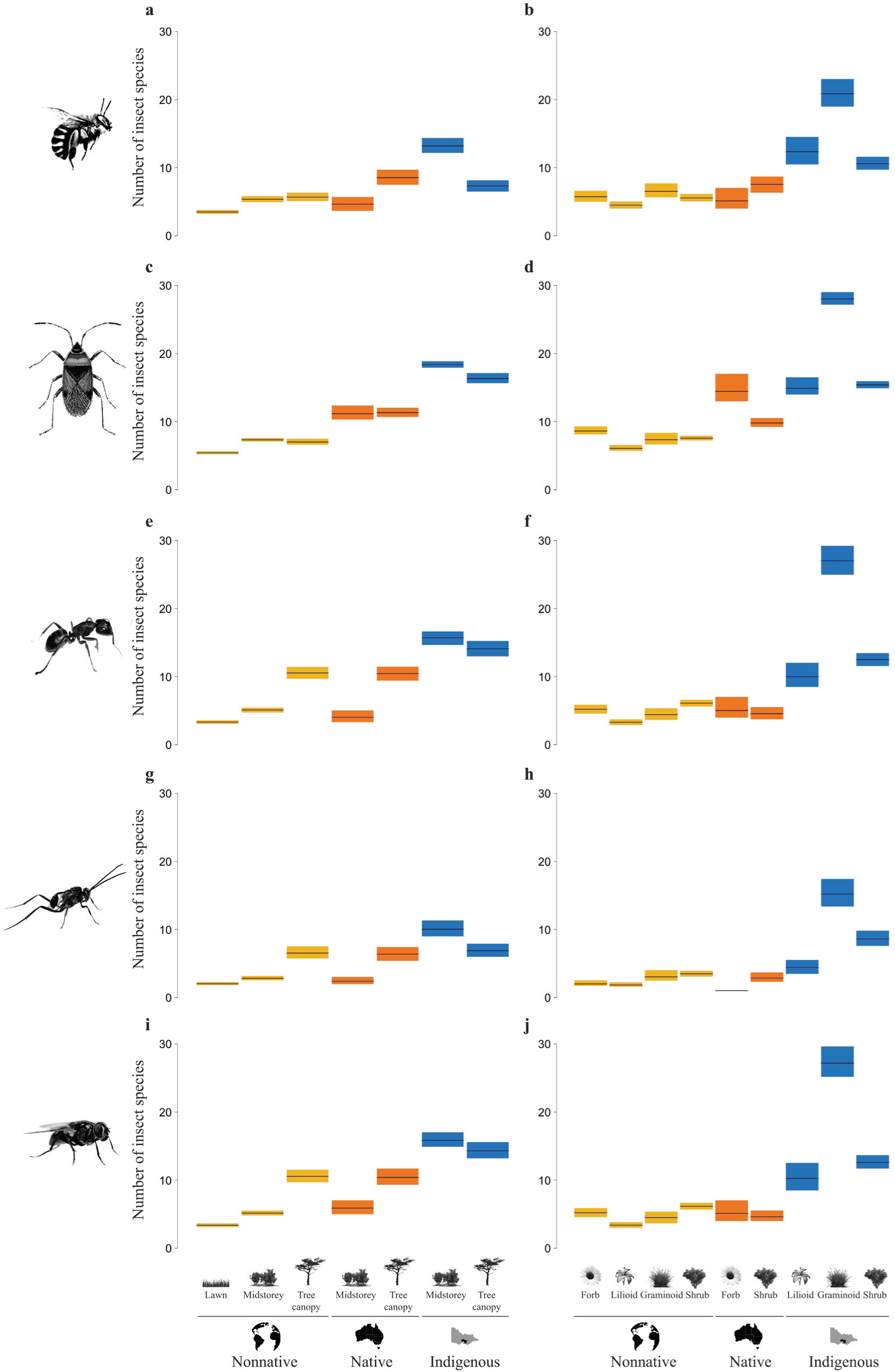
Estimated species richness of indigenous insects by planting design element by plant origin (a,c,e,g,i) and midstorey growth form by plant origin (b,d,f,h,j) for pollinators (a,b), herbivores (c,d), predators (e,f), parasitoids (g,h) and detritivores (i,j). Black lines represent mean responses and coloured boxes the associated statistical uncertainty (95% Credible Intervals). For ease of interpretation plant origin has been colour coded as yellow (nonnative), orange (native) or blue (indigenous).

### Combined effect of growth form and plant origin

Our model estimates indicate that the species richness of indigenous insects varied amongst the growth form by plant origin groups. In general, indigenous groups had higher insect species richness than did native and these had higher insect richness than nonnative (Fig. 4b; Appendix S2: Table S6). The group with the highest insect species richness was indigenous graminoids, with the average plant species in this group showing nearly five times more species than the average nonnative graminoid (Fig. 4b; Appendix S2: Table S6). Indigenous shrubs were associated with the second highest level of insect species richness, with the average indigenous shrub showing 2.1 and 1.6 times more insect species than average nonnative and native shrubs, respectively (Fig. 4b; Appendix S2: Table S6). The group with the third highest insect species richness was indigenous lilioids, with the average indigenous lilioid showing 2.5 times more insect species than the average nonnative lilioid (Fig. 4b; Appendix S2: Table S6).

Indigenous graminoids showed the highest number of indigenous insect species across all insect functional groups (Fig. 5b,d,f,h,j; Appendix S2: Table S6). In general, the indigenous growth form groups were associated with the highest number of indigenous insect species across all insect functional groups (Fig. 5b,d,f,h,j; Appendix S2: Table S6).

### Community composition and unique species

Indigenous insect community composition varied markedly across the design elements by plant origin (Fig. 6a) and growth forms by plant origin (Fig. 6b) groups. The insect composition of a few group pairs was markedly similar such as between indigenous and native midstorey (5%; Fig. 6a) and between nonnative graminoids and shrubs (5%; Fig. 6b). However, most group pairs showed moderate to high dissimilarity (> 20%) in species composition, for example, between native and nonnative tree canopy (32%; Fig. 6a) and between nonnative forbs and lilioids (40%; Fig. 6b).

**Figure 6.**
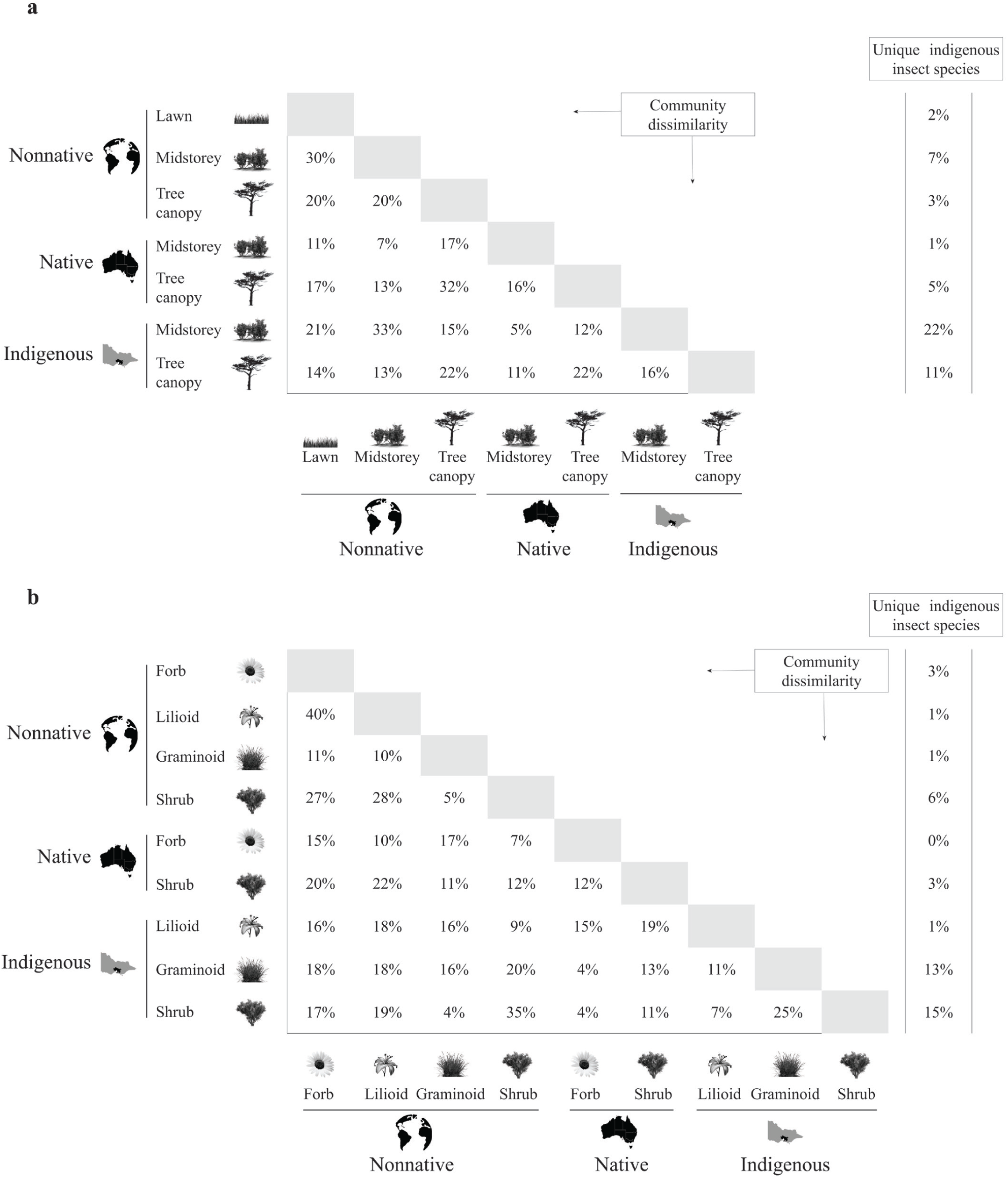
Indigenous insect community dissimilarity matrices for the planting design element by plant origin (a) and midstorey growth form by plant origin (b) groups. Percentages in the white cells were calculated using the Jaccard dissimilarity index, where 0 and 1 represents minimum and maximum dissimilarly, respectively. Values in the adjacent left columns represent the percentage of unique species observed in each group.

The number of unique insect species also varied substantially across groups (Fig. 6). Within the design element by plant origin group, up to 33% of all indigenous insect species recorded in the study were unique to the indigenous groups followed by their nonnative and native counterparts, with 12% and 6% unique species, respectively (Fig. 6a). From the design element perspective, 30% of all indigenous insect species recorded in the study were unique to the midstorey groups, followed by their tree canopy and lawn counterparts, with 19% and 2% unique species, respectively (Fig. 6a). Similarly, within the growth form by plant origin group, as much as 29% of all indigenous insect species recorded in the midstorey were unique to the three indigenous groups, followed by their nonnative and native counterparts, with 11% and 3% unique species, respectively (Fig. 6b). From the growth form perspective, 24% of all indigenous insect species recorded in the midstorey were unique to the shrub groups, followed by the graminoid, forb and lilioid groups, with 14%, 3% and 2% unique species, respectively (Fig. 6b).

## Discussion

Our findings demonstrate that taxonomically and functionally diverse indigenous insect communities occur in greenspaces in densely urbanised inner-city municipalities such as the City of Melbourne, with the potential to boost ecosystem multifunctionality (Soliveres et al. 2016) and biotic resistance against the establishment of introduced insects (Kennedy et al. 2002). We have shown that insect communities in Melbourne greenspaces are predominately composed of indigenous rather than introduced species. Our study further highlights that multi-layered, structurally complex indigenous vegetation plays a core role in sustaining high insect biodiversity in urban areas, with the indigenous midstorey and canopy key to maintaining a rich and functionally diverse indigenous insect community within this system. Within the indigenous midstorey, graminoids surprisingly support the highest indigenous insect species richness across all functional groups – particularly for herbivores, predators and detritivores – followed by indigenous shrubs and lilioids. The indigenous midstorey also hosts the largest percentage of unique species. Taken together our findings emphasise the opportunity presented by indigenous understory and midstorey plants, particularly indigenous graminoids in our study area, to promote indigenous insect biodiversity in urban greenspaces.

### Greenspace insect communities are dominated by indigenous insect species

Our results indicate that the insect community in our study area was composed predominately of indigenous species. This finding aligns with previous studies of insect richness in cities across other continents (Goertzen and Suhling 2014, Sing et al. 2016, Brown and Hartop 2017). Despite the expectation that urban environments act as hotspots for biological invasions (Cadotte et al. 2017), particularly of nonnative plants and insects (Pysek et al 2010), we found a relatively low number of introduced insect species. Interestingly however, some of these introduced species, for example the European honeybee *Apis mellifera*, were common in and widespread across the studied greenspaces. Consequently, the probability of occurrence of introduced insect species on any particular plant across the study area is similar than that of indigenous species.

### Indigenous plant species promote indigenous insect diversity

We found multiple threads of evidence to suggest that indigenous plant species sustain the highest numbers of indigenous insect species and host the largest percentage of unique species. It is generally accepted in restoration ecology that the presence of indigenous plant species promotes recolonisation by indigenous insect species (Moir et al. 2005, Nemec and Bragg 2008). In most cases, phytophagous taxa drive this trend (Procheş et al. 2008, Woodcock et al. 2009) and it is a function of the host-specificity of the insect species, provided that other factors are accounted for, such as the insect’s power of dispersal and suitable micro-climate conditions (Moir et al. 2005).

It follows, therefore, that indigenous plants should encourage the occurrence of indigenous insects, especially for herbivores, but also for other insect functional groups. Indeed, our findings distinctly show that pollinators, herbivores, predators, parasitoid and detritivores reach higher levels of species richness in association with indigenous plant species. Experimental studies also support this assumption (Ballard et al. 2013, Burghardt and Tallamy 2013, Salisbury et al. 2015, 2017). For example, using experimental plantings of tree and shrub species, Burghardt and Tallamy (2013) showed that nonnative plants supported less diverse herbivorous insect communities than indigenous plants. Similarly, in an early successional experiment the biomass, abundance and species richness of herbivorous, predatory and parasitoid insects was lower on nonnative forbs than on indigenous forbs (Ballard et al. 2013). Working specifically in an urban setting, Salisbury and colleagues (2015, 2017) experimented with the origin of flowering plants in garden borders, demonstrating that insects across a diverse range of functional groups were less abundant on nonnative than indigenous plant species. These experimental findings have been substantiated by observational approaches, particularly by studies conducted within urban environments. For instance, a study that included the species-specific responses of bees, beetles and heteropteran bugs to vegetation attributes of gardens, parks and golf courses revealed how occurrence probabilities for most insect species decreased as a function of the amount of nonnative plants present in the studied greenspaces (Threlfall et al. 2017).

### Midstorey as a key planting design element

Our results have shown that the midstorey is a highly valuable ecological asset in urban landscapes in terms of promoting insect diversity. Midstorey vegetation – which in our study also included plants species associated with the understorey – harboured nearly as many indigenous insect species as canopy vegetation. Indigenous midstorey plant species in particular promoted higher levels of overall insect richness, a pattern that was consistent across all insect functional groups. The midstorey also sustained the highest number of unique indigenous insect species. Taken together, our results indicate that the under- and midstorey are underappreciated strata with great potential for supporting insect biodiversity across urban environments. Our findings go beyond the accepted understanding that the greater structural complexity of experimental, restored or managed sites, the higher the taxonomical and functional diversity of insects and other invertebrates (Murdoch et al. 1972, Brown 1984, Majer et al. 2007, Woodcock et al. 2009, Gibb and Cunningham 2010, Mata et al. 2017, Threlfall et al. 2017, Schuldt et al. 2019). We show that within the midstorey it is the indigenous species – particularly indigenous graminoids and shrubs that distinctly outperform their native and nonnative counterparts. Our results are also consistent with previous studies that have documented how insect and other arthropod communities are highly stratified across forest strata (Basset et al. 2003, Ulyshen 2011). It is likely that greenspace midstorey vegetation supports different insect taxa due to differences in habitat structure (e.g. foliage complexity or plant surface textures), microclimate (e.g. light, temperature, wind or humidity differences), unique food resources or particular inter-specific interactions, as has been discussed for temperate deciduous forest (Ulyshen 2011). Unpacking the causal mechanisms for our results would require experimental manipulations that fell beyond the scope of this study.

### Graminoids as a key midstorey growth form

A striking result to emerge from the data is that graminoids sustain the highest number of indigenous insect species across all growth forms and indigenous graminoids show the highest species richness of indigenous insects across all growth form by plant origin groups. Three indigenous tussock grass species made particularly important contributions to supporting indigenous insect biodiversity in our study area: common tussock-grass *Poa labillardierei*, wallaby grass *Rytidosperma sp*. and kangaroo grass *Themeda triandra.* Indeed, *P. labillardierei* had the highest number of indigenous insect species across the study, with any particular tussock grass patch supporting as much as 5.4 times more indigenous insect species than the most speciose lawn complex and 1.7 times more indigenous insect species than the spotted gum *Corymbia maculata*, which was the tree species sustaining the highest number of indigenous insect species across the study. These results substantiate previous findings stressing the relevance of indigenous tussocks and other structurally complex graminoids in providing a diversity of habitat and food resources for the immature and adult life stage of insects and other invertebrates (Tscharntke and Greiler 1995, Morris 2000, Barratt et al. 2005).

Remarkably, the capacity of indigenous graminoids to support the highest levels of indigenous insect species across all studied plant groups was true for all insect functional groups. This finding extends previous studies reporting on the positive effects of tussocks and other graminoids on specific insect functional groups – predominantly on herbivores, predators and parasitoids (Dennis et al. 1998, Woodcock et al. 2007, Haaland et al. 2011), but also for pollinators (Saarinen et al. 2005, Potts et al. 2009). Our finding that graminoids, which are predominantly pollinated by wind, support more pollinators and other flower-visiting insect species than do lilioids or shrubs, which are predominantly pollinated by insects, is highly noteworthy and provides insight into function and value of non-floral resources for insect pollinators (Roulston and Goodell 2011, Requier and Leonhardt 2020). We hope that our research will serve as a base for future studies on the capacity of graminoids to provide habitat and food resources for insects in urban greenspaces, particularly of non-floral resources for pollinators and other flower-visiting species. Beyond graminoids, our data further emphasises the contributions of other midstorey growth forms. For instance, indigenous shrubs and lilioids sustained the second and third highest number of indigenous insect species across the growth from by plant origin groups. Indeed, our findings indicate that any particular indigenous shrub or lilioid species supports the same number of indigenous insect species as any particular indigenous tree species; and substantially more insect species than any particular native or nonnative tree species. However, we found that the number of unique indigenous insect species varied markedly between these two plant groups, with indigenous shrubs showing as much as 7.4 times more unique insect species than their lilioid counterparts. In general, shrubs outperformed all other midstorey growth forms, with approximately one out of every four indigenous insect species across the study being exclusively associated with shrub species.

As underlined by an increasing body of literature (Mata et al. 2017, Threlfall et al. 2017, Aguilera et al. 2019, Norton et al. 2019, Majewska and Altizer 2020), including recent reviews (Burkman and Gardiner 2014, Aronson et al. 2017), and meta-analyses (Beninde et al. 2015), the evidence we found points to the critical role that midstorey growth forms – particularly indigenous plant species – play in supporting taxonomically and functionally diverse indigenous insect communities in urban greenspaces. This finding is not only critical for insect conservation in urban environments but of direct, immediate relevance for a wide range of animals such as reptiles, birds and mammals that rely on them as a primary or complementary food source. Our findings therefore underscore the potential of a diverse, primarily indigenous understorey and midstorey strata to increase the positive biodiversity outcomes provided by structurally complex vegetation. As such, they support ideas that move beyond the stagnant approach of designing urban greenspaces predominantly or often exclusively with nonnative short turfgrass lawn and tall trees (Ikin et al. 2015, Smith et al. 2015, Parris et al. 2018, Norton et al. 2019).

### Limitations and future research

We are aware of some features of our study context and design that might have influenced our results. Firstly, the small proportion of introduced insects found in this study may be the result of recent introductions that have not yet had enough time to develop into established, large populations – a pattern that is common to both the Northern (Roques et al. 2009) and Southern Hemispheres (Ward and Edney-Browne 2015). It is therefore not inconceivable that some introduced species occurring in our study area in small, isolated populations might have gone undetected. Indeed, a recent follow up study in one of the study sites revealed the occurrence of the European firebug *Pyrrhocoris apterus* – a Palaearctic species not previously recorded in Australia (LM unpublished data). As with any observational study, our study design may have introduced unintended bias because our data collection period was purposely designed to coincide with the peak activity season for insects in our study area (i.e., the summer months). We believe however that this effect is negligible as far fewer insect species are more active over the colder periods.

This research has revealed many questions in need of further investigation. While we believe that our finding, that indigenous plants sustain substantially more indigenous insect species than their regionally native and nonnative counterparts, are transferable to other urban environments worldwide, the strong relationships we found between indigenous graminoids and indigenous insects might be less transferable. While graminoids, particularly tussock forming species, were a dominant growth form in our study area before colonial settlement other growth forms might have been more representative in other bioregions. A global study across many bioregions is needed to shed light on this question.

While we have emphasised the critical role of indigenous plants, we have also shown that indigenous insects are being found in association with a wide array of regionally native and nonnative plants. Thus, our study provides considerable insights in support of the idea that urban environments may facilitate novel resource utilisation (Valentine et al. 2020). We recommend future experimental work on this topic to clarify to what extent these associations reflect host shifting patterns unique to urban environments and to fully understand the opportunities and risks provided by novel urban resources. On a wider level, research is also needed to determine to what extent the increases in insect biodiversity provided by complementing lawns and trees with a diverse palette of midstorey growth forms, particularly indigenous species, can boost ecosystem multifunctionality in urban greenspaces – as recently demonstrated for meso- and macrofauna influencing soil multifunctionality (Tresch et al. 2019). The prospect of being able to understand the mechanistic links between, and to quantify the contributions of, increased biodiversity due to greenspace management actions and greenspace multifunctionality serves as a continuous incentive to future research.

### Implications for greenspace design and management

As our research was conducted following the science-government partnerships model (Ives and Lynch 2014), our findings have now been used by the City of Melbourne to provide practical guidance for designing greenspaces that meet the needs of both people and nature. We share these examples of applied ecological knowledge here as a demonstration of how this ecological research can inform practical actions.

In the first instance, our study provides a blueprint and stimulus for built-environment professionals, including architects, engineers, planners and designers, to conceptualise and incorporate into their practice palettes of plant species that foster a larger presence of indigenous plants over regionally native or nonnative species, whilst incorporating a broader mixture of midstorey growth forms. These features are expected to promote taxonomically and functionally diverse indigenous insect communities – even when increasing the amount of greenspace is not feasible due to other pressures (Beninde et al 2015). Integrating these plant palettes into practice may further allow built-environment professionals to plan and design complex plant communities that support and boost indigenous biodiversity in greenspaces and that will likely contribute to bring locally extinct or rare species back into urban environments (Baruch et al. 2020, Mata et al. 2020).

Another promising pathway that can be explored by greenspace professionals includes identifying locations where lawns can be converted to more complex vegetation that includes indigenous plants, particularly, at least in the bioregional context of our study, graminoids, shrub and lilioids. Simple strategies for incorporating more complex vegetation without compromising access to lawn areas include placing the plantings around the greenspace boundaries or under the canopy areas of larger trees where they can also act as a subtle exclusion zone and reduce the risk of injury due to falling limbs or branches. Placing the taller midstorey plantings away from footpaths and other infrastructure can also help meet ‘Crime Prevention Through Environmental Design’ principles by maintaining a line of site (Piroozfar et al. 2019).

### Making a difference – implications for policy and beyond

Through our study we have gained considerable insights that advance knowledge of plant-insect relationships in urban greenspaces. However, to bring about beneficial outcomes for urban landscapes, this scientific evidence must be embedded into policy and, ultimately, operationalised into practice. Indeed, our research was conducted as part of ‘The Little Things that Run the City’ (Mata et al. 2015, 2016), and two of our co-authors were working for the City of Melbourne during the formation and early analysis of this research. Not unexpectedly, project findings have been contributing to inform City of Melbourne decision-making and policy, including the Council’s ‘Nature in the City Strategy’ (City of Melbourne 2017), which includes goals to increase indigenous biodiversity and specific targets to increase plant-related midstorey habitat for insects and other taxa. Project findings have also been applied to develop an insect biodiversity educational portal (http://biodiversity.melbourne.vic.gov.au/insects/) and a children’s book (Cranney et al. 2017). These non-academic outcomes highlight the value of our work and provide encouragement for future partnerships between industry professionals and researchers advocating for and evidencing the value of urban biodiversity.

Another significant approach that could be used by local governments to advance practice aimed at promoting insect biodiversity in urban environments is to incentivise the translation of research findings into landscape design guidelines. These non-mandatory documents, which have traditionally focused on aesthetic outcomes at the expense of biodiversity, can distinctly influence outcomes on the ground. Ultimately however, contractors and consultants responsible for delivering capital works projects will overlook these if the required plants are not readily available when needed. Therefore, the crucial role played by plant nurseries should not be overlooked – an industry where supply tends to influence demand, with growers often limiting production to reliable, profitable and easy to grow plants. Influencing plant supply will require a dedicated engagement with the nursery industry to broaden production to include a wider range of plant species known to support insect biodiversity as those provided evidence for in this study.

Finally, while government officers may apply research findings to design greenspaces capable of supporting diverse insect communities, they must synergistically encourage operational and maintenance programs that can enable these communities to thrive. On ground operational and maintenance teams, that practice greenspace management on a regular basis, have the potential to be involved in identifying opportunities and challenges for supporting insects – and the plants they are more closely associated with – which may expand beyond the obvious choices available to or envisaged by office-based decision-makers. They are, for example, uniquely positioned to readily transfer knowledge on where the optimal soil and microhabitat conditions required for focal plant species within a given greenspace are met – a necessary prerequisite for these to be able to deliver the resources needed by insects to become established and thrive.

## Supporting information

Multi-species hierarchical metacommunity model

## Acknowledgments

This study was funded by the City of Melbourne – thanks to Amy Rogers, Lingna Zhang and other colleagues from the Urban Sustainability Branch for their support and enthusiasm. We would also like to acknowledge funding from RMIT University’s Strategic Projects in Urban Research (SPUR) Fund, the National Environmental Science Programme - Clean Air and Urban Landscapes Hub (NESP - CAUL) and the Australian Research Council - Centre of Excellence for Environmental Decisions (CEED). Thanks to Adam Ślipiński, Ascelin Gordon, David Heathcote, Dayanthi Nugegoda, Flickr community, Friends of Westgate Park, Jeff Shimeta, Laura Stark, Marc Kéry, Michelle Freeman, Rolf Oberprieler, Royal Park managers, Shannon Fernandes, Timothy New and Xavier Francoeur for their invaluable contributions to the study.

## Notes

### Competing Interest Statement

The authors have declared no competing interest.

