## Appendix 1 for "Indigenous plants promote insect biodiversity in urban greenspaces"

### Appendix S1

A schematic representation of our study design is given in Figure S1. The number of plots established in each park (Table S1) varied as a function of the park's area and as a function of the planting design elements (lawn, midstorey and tree canopy) and midstorey growth forms (forb, lilioid, graminoid and shrub) present in each park. In an ideal experimental design with no resource constraints we would have aimed to survey at least 60% of the park's total area (representing equals amounts of the lawn and tree canopy design elements, as well as the four midstorey growth forms). In our study, this would have worked out well for the smallest parks (e.g. Garrard Street Reserve at ~ $10^3$ $m^2$) but would have however become widely unfeasible for large parks (e.g. Royal Park at ~ $10^6$ $m^2$). We therefore estimated the total area to be surveyed in each park using a logarithmic function. Specifically:

$S = 6 \times 100 \times 2^{(\log_{10} A) - 3}$ [Equation S1]

where $S$ equals the total area to be surveyed and $A$ the park's area. This formula satisfies the condition that the total area to be surveyed in a 1,000 $m^2$ site is 600 $m^2$ (60%), whilst yielding proportionally smaller survey areas as park area increases. Likewise, the ideal lawn, midstorey (forb, lilioid, graminoid and shrub) and tree canopy plots would have had an area of 100 $m^2$ (10 x 10 m). In practice, however, the size of each plot $Ps$ was determined by:

$Ps = S / (6 \times Pn)$ [Equation S2]

Where the number of plots $Pn$ of each design element/growth form to be established in each park was defined as:

$Pn = \text{Integer } S / 600$ [Equation S3]

This specification allowed plot size to vary between 75 $m^2$ and 150 $m^2$, and the number of plots of each design element/growth form to be established in each park to vary between one and nine.

The number of observation minutes $DOn$ and sweeps per plant $SNn$ was standardised as a proportion of the given plant's volume $Pv$ ($m^3$) through the following formulas:

$DOn = 1 \text{ min} \times Pv$ [Equation S4]

$SNn = 5 \text{ sweeps} \times Pv$ [Equation S5]

which contributed to make our per plot survey effort proportional to the plot's vegetation volume.

Direct observations were always conducted prior to sweep-netting and consisted on a trained field researcher walking through the plot while taking notes of and/or collecting any insect species observed making contact with the above-ground structure of the targeted plant species (or lawn complexes in the case of lawn plots).

Direct observations were conducted by researchers KC, DD, CI, LM, EP and MP, and a limited number of trained volunteers (Xavier Francoeur, Michelle Freeman and Ascelin Gordon). On occasions insects observed during the direct observations protocol were also photographed using a digital reflex camera equipped with a 100 mm macro lens. These and other related photos are publicly available at the lead author's Flickr photostream:

https://www.flickr.com/photos/dingilingi/albums/72157651179731788

and have been uploaded to the iNaturalist project The Little Things that Run the City:

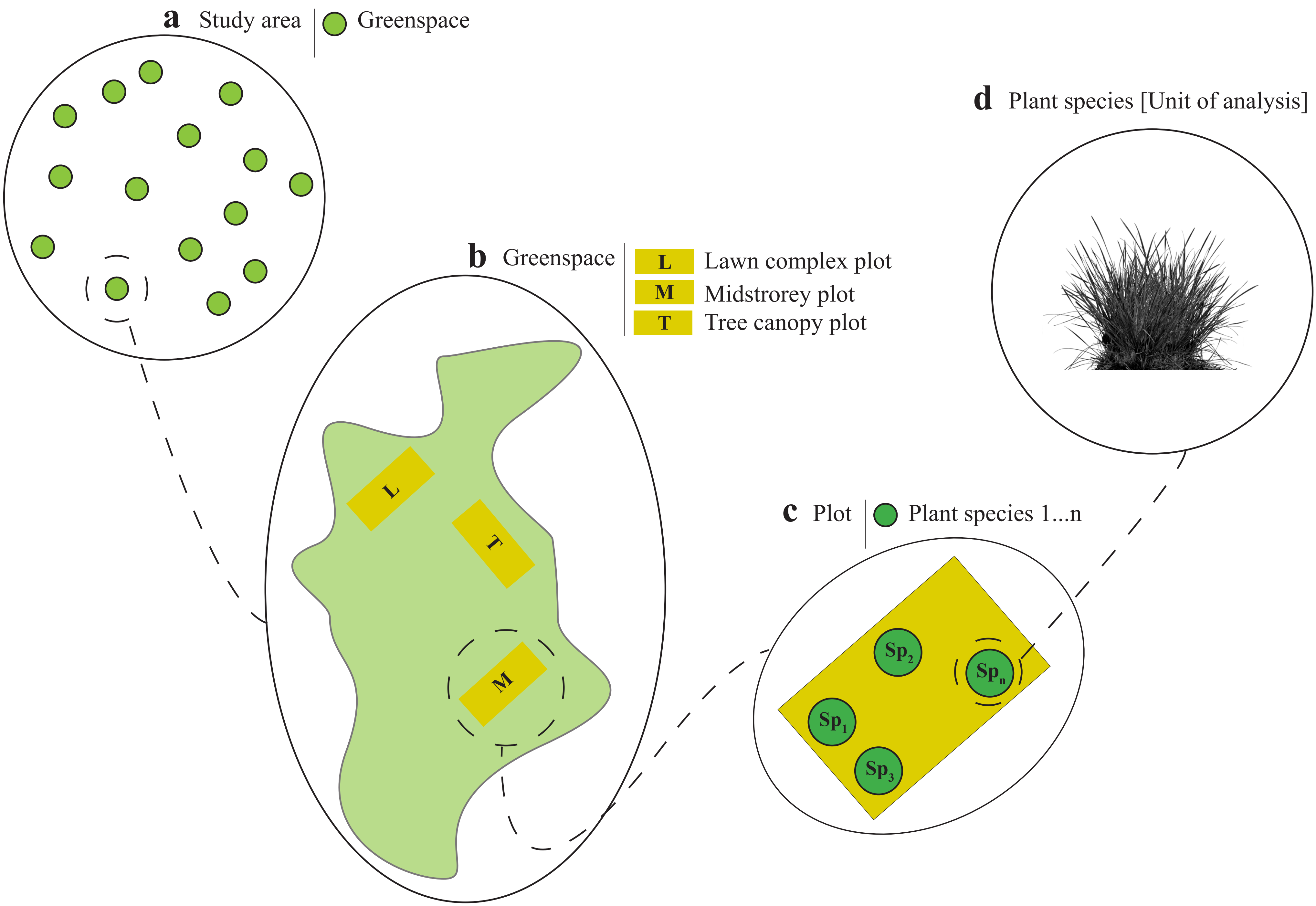


**Figure S1.** Schematic representation of our study design. The study was conducted across 15 public parks (a). Within each park we established at least two plots, for a total of 130 plots; the number of plots established in each park, and their size, varied as a function of the park's area and the planting design element (lawn, midstorey and tree canopy) and midstorey growth forms (forb, lilioid, graminoid and shrub) present in each park (b). Within each plot we located and identified all plant species (c). In total, our study comprised 133 plant species, genera or species complexes (d).

**Table S1.** Name and area of the 15 public parks surveyed in our study, including the area to be surveyed, plot size and number of plots per planting design element/ growth form as given by eqautions S1-S3, and the actual number of plots of each type established in each park.

| Park name | Area ($m^2$)[1] | Area to be surveyed *S* ($m^2$) | Plot size *Ps* ($m^2$) | Number of plots Pn per planting design element/ growth form | Lawn complex plots | Midstorey - Forb plots | Midstorey - Lilioid plots | Midstorey - Graminoid plots | Midstorey - Shrub plots | Midstorey - Mix plots | Tree canaopy plots |
|---|---|---|---|---|---|---|---|---|---|---|---|
| Royal Park | 1,261,946 | 5,148 | 95 | 9 | 9 | | | 9 | 9 | | 9 |
| Princes Park | 329,280 | 3,436 | 95 | 6 | 6 | | 1 | 3 | 1 | | 6 |
| Westgate Park | 239,791 | 3,123 | 104 | 5 | 4 | | | 5 | 5 | | 5 |
| Fitzroy-Treasury Gardens | 175,776 | 2,844 | 95 | 5 | 5 | | | | | 5 | 5 |
| Carlton Gardens South | 47,244 | 1,915 | 106 | 3 | 3 | | | | | 3 | 3 |
| University Square | 12,768 | 1,292 | 108 | 2 | 2 | | | | | | 1 |
| Lincoln Square | 9,865 | 1,195 | 100 | 2 | 2 | | | 1 | 1 | | 2 |
| Arglye Square | 8,335 | 1,136 | 95 | 2 | 2 | | | | | 2 | 2 |
| Women's Peace Gardens | 5,684 | 1,012 | 84 | 2 | 2 | | | | | 2 | 2 |
| Gardiner Reserve | 3,655 | 886 | 148 | 1 | 1 | | | | | | 1 |
| Pleasance Gardens | 3,404 | 868 | 145 | 1 | 1 | | | | | | 1 |
| Murchison Square | 3,294 | 859 | 143 | 1 | 1 | | | | | | 1 |
| State Library of Victoria | 2,400 | 781 | 130 | 1 | 1 | | | | | 1 | 1 |
| Canning/Neill Street Reserve | 1,601 | 691 | 115 | 1 | 1 | | | | | | 1 |
| Garrard Street Reserve | 1,081 | 614 | 102 | 1 | 1 | | | | | | 1 |

1 This figure excludes areas of the park covered by imprevious surfaces

https://www.inaturalist.org/projects/the-little-things-that-run-the-city where they have become public records.

Plant species (Appendix S2: Table S3) within midstorey and tree canopy were sweep-netted individually, while plant species in lawn plots were aggregated into one single sweep-net event. Each lawn plot was considered an independent lawn complex (Appendix S2: Table S3). For tree species we only surveyed to about 3 m in height, assisted, if necessary, by a field step ladder. Most if not all tree species selected for this study had substantial low branching canopies that reached to between 10 and 100 cm of the ground. For sweep-netting we employed an entomological net with a central rod of 90 cm, a bag diameter of 50 cm and a bag depth of 55 cm. In order to minimise collector bias, all sweep-netting was conducted by a single researcher (LM).

Our survey protocol explicitly avoided collecting the immature stages of insect species. Moreover, we put considerable effort in minimising the amount of adult insect material collected. Known species (e.g. the European honeybee *Apis mellifera* and the Passionvine hopper *Scolypopa australis*), either directly observed or sampled in the net, were recorded in situ and released. Only when absolutely necessary for posterior identification were specimens collected, in which case they were directly sampled with a collecting vial or extracted into a collecting vial through an entomological pooter. Specimens were then transferred to a labeled storing vial containing a preservative liquid (70% Ethanol). The assignation of insect species to functional groups also considered the function of the immature stage(s).
