## Appendix 2 for "Indigenous plants promote insect biodiversity in urban greenspaces"

**Appendix S2**

**Figure S1 (Next page).** Plant species by species richness of indigenous insect species (Log10 scale) as estimated under the multi-species community model. Dots represent mean responses and horizontal lines the associated statistical uncertainty (95% Credible Intervals). For ease of interpretation plant origin has been colour coded as yellow (nonnative), orange (native) or blue (indigenous).

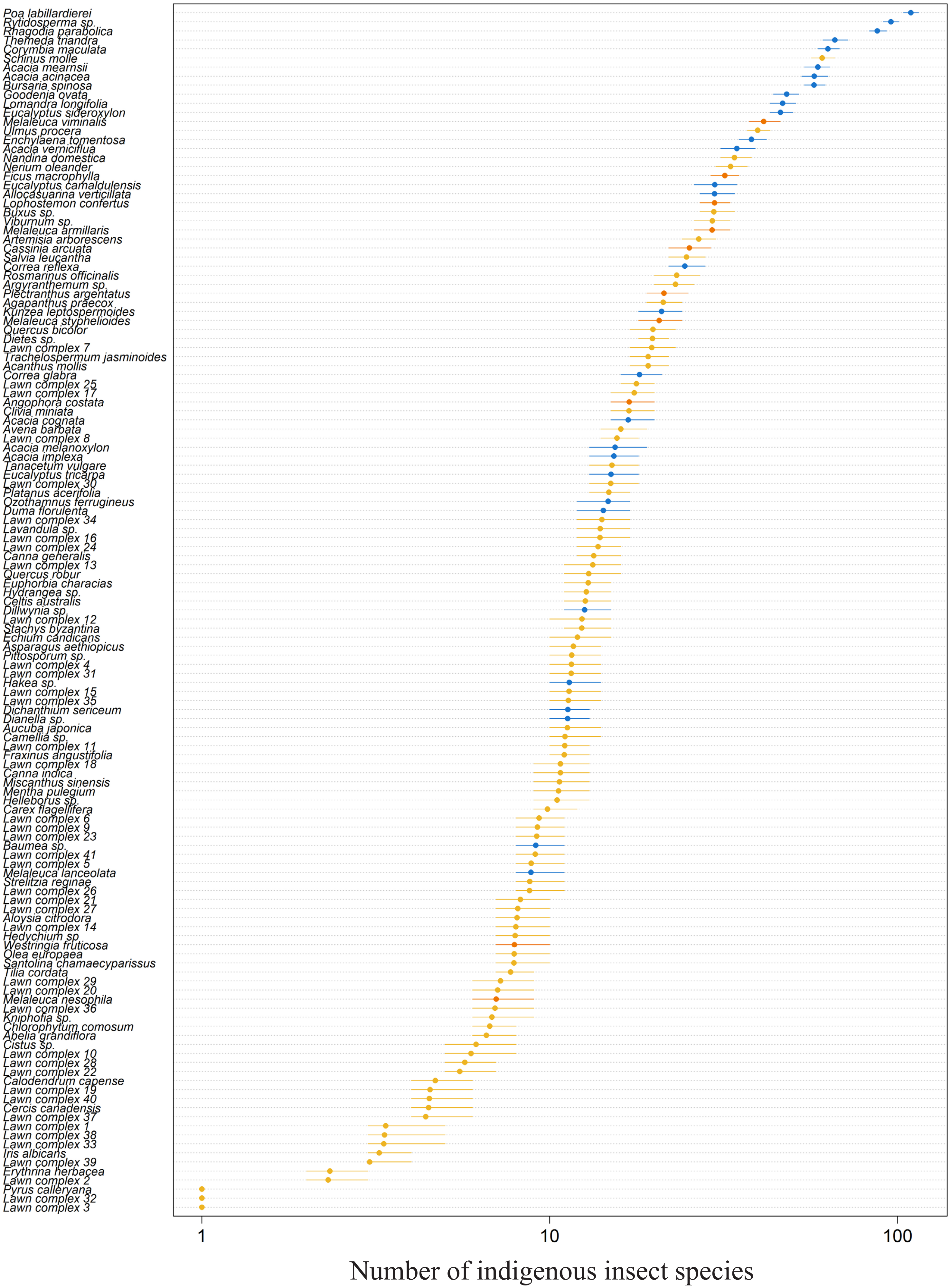
Poa labillardierei
Rytidosperma sp.
Rhagodia parabolica
Themeda triandra
Corymbia maculata
Schinus molle
Acacia mearnsii
Acacia acinacea
Bursaria spinosa
Goodenia ovata
Lomandra longifolia
Eucalyptus sideroxylon
Melaleuca viminalis
Ulmus procera
Enchylaena tomentosa
Acacia verniciflua
Nandina domestica
Nerium oleander
Ficus macrophylla
Eucalyptus camaldulensis
Allocasuarina verticillata
Lophostemon confertus
Buxus sp.
Viburnum sp.
Melaleuca armillaris
Artemisia arborescens
Cassinia arcuata
Salvia leucantha
Correa reflexa
Rosmarinus officinalis
Argyranthemum sp.
Plectranthus argentatus
Agapanthus praecox
Kunzea leptospermoides
Melaleuca styphelioides
Quercus bicolor
Dietes sp.
Lawn complex 7
Trachelospermum jasminoides
Acanthus mollis
Correa glabra
Lawn complex 25
Lawn complex 17
Angophora costata
Clivia miniata
Acacia cognata
Avena barbata
Lawn complex 8
Acacia melanoxylon
Acacia implexa
Tanacetum vulgare
Eucalyptus tricarpa
Lawn complex 30
Platanus acerifolia
Ozothamnus ferrugineus
Duma florulenta
Lawn complex 34
Lavandula sp.
Lawn complex 16
Lawn complex 24
Canna generalis
Lawn complex 13
Quercus robur
Euphorbia characias
Hydrangea sp.
Celtis australis
Dillwynia sp.
Lawn complex 12
Stachys byzantina
Echium candicans
Asparagus aethiopicus
Pittosporum sp.
Lawn complex 4
Lawn complex 31
Hakea sp.
Lawn complex 15
Lawn complex 35
Dichanthium sericeum
Dianella sp.
Aucuba japonica
Camellia sp.
Lawn complex 11
Fraxinus angustifolia
Lawn complex 18
Canna indica
Miscanthus sinensis
Mentha pulegium
Helleborus sp.
Carex flagellifera
Lawn complex 6
Lawn complex 9
Lawn complex 23
Baumea sp.
Lawn complex 41
Lawn complex 5
Melaleuca lanceolata
Strelitzia reginae
Lawn complex 26
Lawn complex 21
Lawn complex 27
Aloysia citrodora
Lawn complex 14
Hedychium sp
Westringia fruticosa
Olea europaea
Santolina chamaecyparissus
Tilia cordata
Lawn complex 29
Lawn complex 20
Melaleuca nesophila
Lawn complex 36
Kniphofia sp.
Chlorophytum comosum
Abelia grandiflora
Cistus sp.
Lawn complex 10
Lawn complex 28
Lawn complex 22
Calodendrum capense
Lawn complex 19
Lawn complex 40
Cercis canadensis
Lawn complex 37
Lawn complex 1
Lawn complex 38
Lawn complex 33
Iris albicans
Lawn complex 39
Erythrina herbacea
Lawn complex 2
Pyrus calleryana
Lawn complex 32
Lawn complex 3
1
10
100
Number of indigenous insect species

**Table S1.** The 552 insect species observed in th study, their taxonomical and functional associations, and the posterior estimates for their species-specific probabilities of occurrence and detection as estimated under the multi-species community model. Introduced insect species are indicated with an *. POL: Pollinator or flower-visiting; HER: Herbivore; PRE: Predator; PAR: Parasitoid; DET: Detritivore.

| Insect group \| Species/morphospecies | Family | POL | HER | PRE | PAR | DET | Probability of occurrence | | | | Probability of detection | | | |
|---|---|---|---|---|---|---|---|---|---|---|---|---|---|---|
| | | | | | | | Mean | SD | 2.50% | 97.50% | Mean | SD | 2.50% | 97.50% |
| **Ants [Hymenoptera: Apocrita: Aculeata: Vespoidea: Formicidae]** | | | | | | | | | | | | | | |
| *Linepithema humile* * | Formicidae | 0 | 1 | 1 | 0 | 1 | 0.519 | 0.077 | 0.376 | 0.679 | 0.202 | 0.024 | 0.157 | 0.252 |
| *Camponotus consobrinus* | Formicidae | 0 | 1 | 1 | 0 | 1 | 0.127 | 0.100 | 0.018 | 0.395 | 0.022 | 0.016 | 0.004 | 0.065 |
| *Colobopsis gasseri* | Formicidae | 0 | 0 | 1 | 0 | 1 | 0.189 | 0.120 | 0.045 | 0.509 | 0.029 | 0.019 | 0.007 | 0.077 |
| *Iridomyrmex sp. 1* | Formicidae | 1 | 1 | 1 | 0 | 1 | 0.705 | 0.060 | 0.587 | 0.825 | 0.278 | 0.022 | 0.235 | 0.322 |
| *Melophorus sp. 1 (frugutae group)* | Formicidae | 0 | 0 | 1 | 0 | 1 | 0.116 | 0.084 | 0.024 | 0.327 | 0.041 | 0.027 | 0.010 | 0.114 |
| *Melophorus sp. 2 (turneri group)* | Formicidae | 0 | 0 | 1 | 0 | 1 | 0.098 | 0.076 | 0.016 | 0.302 | 0.036 | 0.024 | 0.007 | 0.095 |
| *Monomorium sp. 1 (sydneyense group)* | Formicidae | 0 | 0 | 1 | 0 | 1 | 0.162 | 0.103 | 0.042 | 0.419 | 0.038 | 0.020 | 0.011 | 0.086 |
| *Notoncus sp. 1 (ectatommoides group)* | Formicidae | 0 | 0 | 1 | 0 | 1 | 0.121 | 0.091 | 0.021 | 0.356 | 0.027 | 0.024 | 0.005 | 0.091 |
| *Nylanderia rosae* | Formicidae | 1 | 1 | 1 | 0 | 1 | 0.299 | 0.086 | 0.166 | 0.503 | 0.125 | 0.037 | 0.064 | 0.206 |
| *Ochetellus sp. 1 (glaber¬†group)* | Formicidae | 0 | 0 | 1 | 0 | 1 | 0.352 | 0.138 | 0.158 | 0.690 | 0.058 | 0.023 | 0.024 | 0.109 |
| *Pheidole sp. 1* | Formicidae | 0 | 1 | 1 | 0 | 1 | 0.343 | 0.130 | 0.147 | 0.662 | 0.059 | 0.026 | 0.023 | 0.124 |
| *Pheidole vigilans* | Formicidae | 0 | 1 | 1 | 0 | 1 | 0.180 | 0.092 | 0.062 | 0.418 | 0.065 | 0.032 | 0.021 | 0.144 |
| *Prolasius sp. 1 (nitidissimus group)* | Formicidae | 0 | 1 | 1 | 0 | 1 | 0.522 | 0.109 | 0.332 | 0.756 | 0.086 | 0.021 | 0.052 | 0.133 |
| *Rhytidoponera metallica* | Formicidae | 0 | 1 | 1 | 0 | 1 | 0.154 | 0.103 | 0.032 | 0.430 | 0.025 | 0.017 | 0.006 | 0.069 |
| *Rhytidoponera victoriae* | Formicidae | 0 | 0 | 1 | 0 | 1 | 0.228 | 0.106 | 0.087 | 0.518 | 0.057 | 0.022 | 0.023 | 0.110 |
| *Tetramorium bicarinatum* | Formicidae | 0 | 0 | 1 | 0 | 1 | 0.199 | 0.114 | 0.059 | 0.510 | 0.051 | 0.029 | 0.014 | 0.126 |
| *Turneria sp. nov.* | Formicidae | 0 | 0 | 1 | 0 | 1 | 0.114 | 0.084 | 0.020 | 0.333 | 0.029 | 0.027 | 0.005 | 0.096 |
| **Bees [Hymenoptera: Apocrita: Aculeata: Apoidea]** | | | | | | | | | | | | | | |
| *Apis mellifera* * | Apidae | 1 | 1 | 0 | 0 | 0 | 0.367 | 0.103 | 0.204 | 0.607 | 0.113 | 0.032 | 0.059 | 0.184 |
| *Callohesma sp. 1* | Colletidae | 1 | 1 | 0 | 0 | 0 | 0.122 | 0.090 | 0.020 | 0.364 | 0.021 | 0.015 | 0.004 | 0.060 |
| *Colletidae 1* | Colletidae | 1 | 1 | 0 | 0 | 0 | 0.113 | 0.094 | 0.018 | 0.382 | 0.029 | 0.025 | 0.005 | 0.096 |
| *Crabronidae 1* | Crabronidae | 1 | 1 | 1 | 0 | 0 | 0.147 | 0.097 | 0.030 | 0.400 | 0.025 | 0.016 | 0.006 | 0.067 |
| *Crabronidae 2* | Crabronidae | 1 | 1 | 1 | 0 | 0 | 0.160 | 0.109 | 0.031 | 0.441 | 0.030 | 0.025 | 0.006 | 0.095 |
| *Euryglossina sp. 1* | Colletidae | 1 | 1 | 0 | 0 | 0 | 0.101 | 0.081 | 0.017 | 0.317 | 0.031 | 0.026 | 0.005 | 0.103 |
| *Homalictus bisbanensis* | Halictidae | 1 | 1 | 0 | 0 | 0 | 0.115 | 0.093 | 0.018 | 0.371 | 0.029 | 0.025 | 0.005 | 0.099 |
| *Homalictus punctatus* | Halticidae | 1 | 1 | 0 | 0 | 0 | 0.111 | 0.084 | 0.018 | 0.336 | 0.029 | 0.025 | 0.005 | 0.095 |
| *Homalictus sphecodoides* | Halticidae | 1 | 1 | 0 | 0 | 0 | 0.271 | 0.122 | 0.089 | 0.568 | 0.038 | 0.017 | 0.014 | 0.081 |
| *Hylaeus sp. 1* | Colletidae | 1 | 1 | 0 | 0 | 0 | 0.123 | 0.086 | 0.021 | 0.350 | 0.026 | 0.022 | 0.005 | 0.084 |
| *Hyphesma atromicans* | Colletidae | 1 | 1 | 0 | 0 | 0 | 0.129 | 0.094 | 0.021 | 0.367 | 0.022 | 0.016 | 0.005 | 0.065 |
| *Lasioglossum clelandi* | Halictidae | 1 | 1 | 0 | 0 | 0 | 0.110 | 0.080 | 0.020 | 0.319 | 0.025 | 0.019 | 0.005 | 0.074 |
| *Lasioglossum cognatum* | Halictidae | 1 | 1 | 0 | 0 | 0 | 0.110 | 0.081 | 0.017 | 0.330 | 0.025 | 0.019 | 0.005 | 0.073 |
| *Lasioglossum hemichalceum* | Halictidae | 1 | 1 | 0 | 0 | 0 | 0.213 | 0.117 | 0.052 | 0.503 | 0.024 | 0.014 | 0.006 | 0.061 |
| *Lasioglossum quadratum* | Halictidae | 1 | 1 | 0 | 0 | 0 | 0.112 | 0.085 | 0.017 | 0.346 | 0.024 | 0.018 | 0.005 | 0.070 |
| *Lipotriches flavoviridis* | Halictidae | 1 | 1 | 0 | 0 | 0 | 0.130 | 0.114 | 0.018 | 0.477 | 0.023 | 0.018 | 0.004 | 0.072 |
| **Beetles [Coleoptera]** | | | | | | | | | | | | | | |
| *Apinocis variipennis* * | Curculionidae | 0 | 1 | 0 | 0 | 0 | 0.146 | 0.124 | 0.024 | 0.533 | 0.024 | 0.019 | 0.004 | 0.071 |
| *Archaeocrypticus topali* * | Archaeocrypticidae | 0 | 1 | 0 | 0 | 1 | 0.142 | 0.111 | 0.018 | 0.425 | 0.026 | 0.025 | 0.004 | 0.095 |
| *Atrichonotus sordidus* * | Curculionidae | 0 | 1 | 0 | 0 | 0 | 0.150 | 0.118 | 0.022 | 0.474 | 0.025 | 0.022 | 0.004 | 0.087 |
| *Derelomini 1* * | Curculionidae | 0 | 1 | 0 | 0 | 0 | 0.148 | 0.126 | 0.023 | 0.510 | 0.023 | 0.019 | 0.004 | 0.075 |
| *Diachus auratus* * | Chrysomelidae | 0 | 1 | 0 | 0 | 0 | 0.237 | 0.101 | 0.095 | 0.485 | 0.066 | 0.028 | 0.024 | 0.133 |
| *Heteronychus arator* * | Scarabaeidae | 0 | 1 | 0 | 0 | 1 | 0.148 | 0.132 | 0.022 | 0.564 | 0.026 | 0.024 | 0.004 | 0.089 |
| *Hippodamia variegata* * | Coccinellidae | 0 | 0 | 1 | 0 | 0 | 0.241 | 0.127 | 0.069 | 0.552 | 0.042 | 0.021 | 0.014 | 0.095 |
| *Leucohimatium arundinaceum* * | Erotylidae | 0 | 1 | 0 | 0 | 1 | 0.173 | 0.110 | 0.046 | 0.491 | 0.058 | 0.035 | 0.015 | 0.154 |
| *Listronotus sp. 1 (bonariensis group)* * | Curculionidae | 0 | 1 | 0 | 0 | 0 | 0.239 | 0.131 | 0.064 | 0.565 | 0.035 | 0.025 | 0.008 | 0.102 |
| *Meligethes sp. 1* * | Nitidulidae | 0 | 0 | 0 | 0 | 1 | 0.172 | 0.112 | 0.043 | 0.496 | 0.040 | 0.026 | 0.010 | 0.107 |
| *Naupactus cervinus* * | Curculionidae | 0 | 1 | 0 | 0 | 0 | 0.331 | 0.157 | 0.109 | 0.693 | 0.044 | 0.023 | 0.014 | 0.100 |
| *Phlyctinus callosus* * | Curculionidae | 0 | 1 | 0 | 0 | 0 | 0.169 | 0.121 | 0.031 | 0.512 | 0.029 | 0.024 | 0.006 | 0.098 |
| *Phyllotreta undulata* * | Chrysomelidae | 0 | 1 | 0 | 0 | 0 | 0.158 | 0.128 | 0.026 | 0.525 | 0.021 | 0.017 | 0.004 | 0.066 |
| *Sitona discoideus* * | Curculionidae | 0 | 1 | 0 | 0 | 0 | 0.165 | 0.122 | 0.023 | 0.503 | 0.020 | 0.015 | 0.004 | 0.061 |
| *Xanthogaleruca luteola* * | Chrysomelidae | 0 | 1 | 0 | 0 | 0 | 0.275 | 0.125 | 0.110 | 0.624 | 0.064 | 0.026 | 0.024 | 0.123 |
| *Acalonoma sp. 1 (pusillum group)* | Curculionidae | 0 | 1 | 0 | 0 | 0 | 0.121 | 0.086 | 0.018 | 0.345 | 0.024 | 0.020 | 0.005 | 0.075 |
| *Aderidae 1* | Aderidae | 0 | 1 | 0 | 0 | 1 | 0.312 | 0.154 | 0.095 | 0.692 | 0.037 | 0.020 | 0.012 | 0.089 |
| *Aderus sp. 1* | Aderidae | 0 | 1 | 0 | 0 | 1 | 0.116 | 0.088 | 0.020 | 0.368 | 0.027 | 0.022 | 0.005 | 0.089 |
| *Aethina concolor* | Nitidulidae | 0 | 0 | 0 | 0 | 1 | 0.203 | 0.116 | 0.051 | 0.486 | 0.037 | 0.024 | 0.009 | 0.099 |
| *Agrypnus sp. 1* | Elateridae | 0 | 1 | 0 | 0 | 1 | 0.106 | 0.088 | 0.017 | 0.372 | 0.031 | 0.026 | 0.005 | 0.102 |
| *Aleocharinae 1* | Staphylinidae | 0 | 0 | 1 | 0 | 0 | 0.153 | 0.116 | 0.028 | 0.489 | 0.032 | 0.027 | 0.006 | 0.104 |
| *Amlearcha sp. 1* | Nitidulidae | 0 | 0 | 0 | 0 | 1 | 0.163 | 0.104 | 0.037 | 0.441 | 0.038 | 0.028 | 0.008 | 0.110 |
| *Ancyttalia sp. 1 (erichsoni group)* | Curculionidae | 0 | 1 | 0 | 0 | 0 | 0.139 | 0.089 | 0.029 | 0.369 | 0.033 | 0.026 | 0.007 | 0.103 |
| *Anthicidae 1* | Anthicidae | 0 | 1 | 1 | 0 | 1 | 0.203 | 0.119 | 0.050 | 0.518 | 0.026 | 0.016 | 0.007 | 0.066 |
| *Anthicus obliquifasciatus* | Anthicidae | 0 | 1 | 1 | 0 | 1 | 0.152 | 0.097 | 0.031 | 0.396 | 0.028 | 0.020 | 0.006 | 0.079 |
| *Anthrenocerus sp. 1* | Dermestidae | 1 | 1 | 0 | 0 | 1 | 0.142 | 0.098 | 0.029 | 0.405 | 0.030 | 0.021 | 0.007 | 0.083 |
| *Anthrenus verbasci* | Dermestidae | 1 | 1 | 0 | 0 | 1 | 0.121 | 0.087 | 0.019 | 0.359 | 0.026 | 0.021 | 0.005 | 0.081 |
| *Apion sp. 1* | Brentidae | 0 | 1 | 0 | 0 | 0 | 0.108 | 0.084 | 0.019 | 0.337 | 0.028 | 0.023 | 0.005 | 0.088 |
| *Apion sp. 2* | Brentidae | 0 | 1 | 0 | 0 | 0 | 0.125 | 0.101 | 0.023 | 0.400 | 0.022 | 0.016 | 0.004 | 0.063 |
| *Apolinus sp. 1* | Coccinellidae | 0 | 0 | 1 | 0 | 0 | 0.172 | 0.116 | 0.038 | 0.469 | 0.036 | 0.025 | 0.008 | 0.103 |
| *Araecerus sp. 1 (lindensis group)* | Anthribidae | 0 | 1 | 0 | 0 | 1 | 0.254 | 0.140 | 0.072 | 0.581 | 0.024 | 0.013 | 0.007 | 0.058 |
| *Balanophorus sp. 1* | Melyridae | 1 | 0 | 1 | 0 | 0 | 0.172 | 0.125 | 0.028 | 0.504 | 0.027 | 0.021 | 0.006 | 0.089 |
| *Bruchidius sp. 1* | Chrysomelidae | 0 | 1 | 0 | 0 | 0 | 0.146 | 0.099 | 0.028 | 0.408 | 0.029 | 0.021 | 0.006 | 0.085 |
| *Chaetocnema sp. 1* | Chrysomelidae | 0 | 1 | 0 | 0 | 0 | 0.421 | 0.096 | 0.261 | 0.638 | 0.113 | 0.025 | 0.068 | 0.166 |
| *Chaetocnema sp. 2* | Chrysomelidae | 0 | 1 | 0 | 0 | 0 | 0.143 | 0.108 | 0.039 | 0.414 | 0.089 | 0.060 | 0.018 | 0.240 |
| *Chauliognathus lugubris* | Cantharidae | 1 | 1 | 1 | 0 | 0 | 0.292 | 0.134 | 0.103 | 0.615 | 0.033 | 0.015 | 0.012 | 0.071 |
| *Coccinella transversalis* | Coccinellidae | 0 | 0 | 1 | 0 | 0 | 0.169 | 0.121 | 0.032 | 0.509 | 0.026 | 0.018 | 0.005 | 0.071 |
| *Conoderus sp. 1* | Elateridae | 0 | 1 | 0 | 0 | 1 | 0.151 | 0.105 | 0.031 | 0.434 | 0.030 | 0.022 | 0.006 | 0.087 |
| *Corticaria sp.1* | Latriidae | 0 | 1 | 0 | 0 | 1 | 0.147 | 0.105 | 0.028 | 0.414 | 0.030 | 0.022 | 0.006 | 0.090 |
| *Cortinicara sp.1* | Latriidae | 0 | 1 | 0 | 0 | 1 | 0.877 | 0.035 | 0.805 | 0.940 | 0.567 | 0.021 | 0.527 | 0.608 |
| *Corylophidae 1* | Corylophidae | 0 | 1 | 0 | 0 | 1 | 0.121 | 0.101 | 0.018 | 0.403 | 0.022 | 0.016 | 0.004 | 0.063 |
| *Cryptamorpha sp. 1* | Silvnidae | 0 | 1 | 0 | 0 | 1 | 0.154 | 0.108 | 0.032 | 0.450 | 0.031 | 0.024 | 0.006 | 0.096 |
| *Cryptolaemus montrouzeri* | Coccinellidae | 0 | 0 | 1 | 0 | 0 | 0.111 | 0.087 | 0.018 | 0.351 | 0.026 | 0.021 | 0.005 | 0.081 |
| *Curculionidae 1* | Curculionidae | 0 | 1 | 0 | 0 | 0 | 0.104 | 0.078 | 0.017 | 0.310 | 0.030 | 0.024 | 0.006 | 0.097 |
| *Curculionidae 2* | Curculionidae | 0 | 1 | 0 | 0 | 0 | 0.136 | 0.100 | 0.022 | 0.409 | 0.022 | 0.017 | 0.004 | 0.067 |
| *Cydmaea sp. 1* | Curculionidae | 0 | 1 | 0 | 0 | 0 | 0.157 | 0.112 | 0.030 | 0.457 | 0.030 | 0.023 | 0.006 | 0.092 |
| *Dasytinae 1* | Melyridae | 1 | 0 | 1 | 0 | 0 | 0.112 | 0.089 | 0.017 | 0.356 | 0.026 | 0.019 | 0.005 | 0.075 |
| *Dicranolaius bellulus* | Melyridae | 1 | 0 | 1 | 0 | 0 | 0.267 | 0.130 | 0.087 | 0.573 | 0.029 | 0.015 | 0.009 | 0.067 |
| *Diomus notescens* | Coccinellidae | 0 | 0 | 1 | 0 | 0 | 0.307 | 0.133 | 0.112 | 0.618 | 0.051 | 0.024 | 0.017 | 0.108 |
| *Diomus sp. 1* | Coccinellidae | 0 | 0 | 1 | 0 | 0 | 0.345 | 0.083 | 0.206 | 0.523 | 0.118 | 0.025 | 0.074 | 0.171 |
| *Diomus sp. 2* | Coccinellidae | 0 | 0 | 1 | 0 | 0 | 0.364 | 0.159 | 0.121 | 0.732 | 0.036 | 0.017 | 0.013 | 0.079 |
| *Diomus sp. 3* | Coccinellidae | 0 | 0 | 1 | 0 | 0 | 0.301 | 0.087 | 0.159 | 0.505 | 0.097 | 0.025 | 0.055 | 0.152 |
| *Diomus sp. 4* | Coccinellidae | 0 | 0 | 1 | 0 | 0 | 0.177 | 0.095 | 0.050 | 0.412 | 0.059 | 0.036 | 0.015 | 0.151 |
| *Diomus sp. 5* | Coccinellidae | 0 | 0 | 1 | 0 | 0 | 0.200 | 0.101 | 0.074 | 0.471 | 0.070 | 0.031 | 0.023 | 0.143 |
| *Ditropidus sp. 1 (aeneolus group)* | Chrysomelidae | 0 | 1 | 0 | 0 | 0 | 0.090 | 0.076 | 0.014 | 0.320 | 0.046 | 0.037 | 0.008 | 0.140 |
| *Ditropidus sp. 2* | Chrysomelidae | 0 | 1 | 0 | 0 | 0 | 0.079 | 0.057 | 0.018 | 0.243 | 0.087 | 0.059 | 0.016 | 0.239 |
| *Ditropidus sp. 3* | Chrysomelidae | 0 | 1 | 0 | 0 | 0 | 0.232 | 0.126 | 0.062 | 0.545 | 0.031 | 0.019 | 0.008 | 0.079 |
| *Ditropidus sp. 4* | Chrysomelidae | 0 | 1 | 0 | 0 | 0 | 0.097 | 0.066 | 0.016 | 0.266 | 0.031 | 0.027 | 0.006 | 0.102 |
| *Dryophiloides sp. 1* | Ptinidae | 0 | 0 | 0 | 0 | 1 | 0.167 | 0.108 | 0.033 | 0.450 | 0.025 | 0.018 | 0.005 | 0.069 |

| | | | | | | | | | | | | | | |
|---|---|---|---|---|---|---|---|---|---|---|---|---|---|---|
| *Dryophiloides sp. 2* | Ptinidae | 0 | 0 | 0 | 0 | 1 | 0.074 | 0.054 | 0.016 | 0.215 | 0.093 | 0.055 | 0.023 | 0.233 |
| *Dryophiloides sp. 3* | Ptinidae | 0 | 0 | 0 | 0 | 1 | 0.133 | 0.088 | 0.031 | 0.366 | 0.052 | 0.035 | 0.011 | 0.145 |
| *Eboo sp. 1* | Chrysomelidae | 0 | 1 | 0 | 0 | 0 | 0.182 | 0.109 | 0.036 | 0.440 | 0.021 | 0.014 | 0.005 | 0.057 |
| *Ecnolagria sp. 1* | Tenebrionidae | 0 | 1 | 0 | 0 | 1 | 0.227 | 0.131 | 0.053 | 0.557 | 0.023 | 0.014 | 0.006 | 0.059 |
| *Elateridae 1* | Elateridae | 0 | 1 | 0 | 0 | 1 | 0.135 | 0.090 | 0.029 | 0.370 | 0.035 | 0.028 | 0.007 | 0.109 |
| *Emblesis sp. 1* | Curculionidae | 0 | 1 | 0 | 0 | 0 | 0.344 | 0.138 | 0.127 | 0.658 | 0.038 | 0.017 | 0.016 | 0.080 |
| *Emblesis sp. 2* | Curculionidae | 0 | 1 | 0 | 0 | 0 | 0.256 | 0.110 | 0.097 | 0.516 | 0.047 | 0.023 | 0.016 | 0.105 |
| *Epamoebus sp. 1* | Curculionidae | 0 | 1 | 0 | 0 | 0 | 0.160 | 0.099 | 0.035 | 0.397 | 0.038 | 0.028 | 0.009 | 0.112 |
| *Epuraea sp. 1* | Nitidulidae | 0 | 0 | 0 | 0 | 1 | 0.132 | 0.078 | 0.040 | 0.352 | 0.095 | 0.049 | 0.026 | 0.209 |
| *Euciodes sp. 1 (suturalis group)* | Anthribidae | 0 | 1 | 0 | 0 | 1 | 0.263 | 0.132 | 0.074 | 0.574 | 0.031 | 0.015 | 0.010 | 0.068 |
| *Eurispa sp. 1* | Chrysomelidae | 0 | 1 | 0 | 0 | 0 | 0.133 | 0.091 | 0.022 | 0.354 | 0.021 | 0.016 | 0.004 | 0.062 |
| *Euthyphasis parva* | Curculionidae | 0 | 1 | 0 | 0 | 0 | 0.115 | 0.081 | 0.023 | 0.340 | 0.046 | 0.032 | 0.010 | 0.130 |
| *Floydwernerius australis* | Anthicidae | 0 | 1 | 1 | 0 | 1 | 0.117 | 0.085 | 0.019 | 0.343 | 0.022 | 0.016 | 0.005 | 0.064 |
| *Harmonia conformis* | Coccinellidae | 0 | 0 | 1 | 0 | 0 | 0.421 | 0.151 | 0.168 | 0.737 | 0.035 | 0.016 | 0.014 | 0.073 |
| *Helcogaster varius* | Melyridae | 1 | 0 | 1 | 0 | 0 | 0.126 | 0.099 | 0.018 | 0.401 | 0.024 | 0.019 | 0.004 | 0.073 |
| *Helochares tristis* | Hydrophilidae | 0 | 0 | 1 | 0 | 1 | 0.326 | 0.136 | 0.117 | 0.638 | 0.033 | 0.015 | 0.012 | 0.070 |
| *Hispellinus multispinosus* | Chrysomelidae | 0 | 1 | 0 | 0 | 0 | 0.085 | 0.069 | 0.013 | 0.276 | 0.052 | 0.045 | 0.009 | 0.175 |
| *Idaethina sp. 1 (pilistriata group)* | Nitidulidae | 0 | 0 | 0 | 0 | 1 | 0.123 | 0.093 | 0.023 | 0.348 | 0.047 | 0.035 | 0.010 | 0.142 |
| *Idaethina sp. 2* | Nitidulidae | 0 | 0 | 0 | 0 | 1 | 0.109 | 0.080 | 0.019 | 0.323 | 0.028 | 0.023 | 0.005 | 0.091 |
| *Illeis galbula* | Coccinellidae | 0 | 0 | 1 | 0 | 0 | 0.154 | 0.120 | 0.025 | 0.489 | 0.033 | 0.028 | 0.006 | 0.103 |
| *Lemidia concinna* | Cleridae | 1 | 1 | 1 | 0 | 0 | 0.118 | 0.096 | 0.019 | 0.398 | 0.026 | 0.021 | 0.005 | 0.083 |
| *Leptopius sp. 1 (robustus group)* | Curculionidae | 0 | 1 | 0 | 0 | 0 | 0.121 | 0.082 | 0.021 | 0.328 | 0.022 | 0.017 | 0.004 | 0.071 |
| *Litochrus sp. 1* | Phalacridae | 1 | 1 | 0 | 0 | 1 | 0.180 | 0.128 | 0.037 | 0.543 | 0.033 | 0.023 | 0.007 | 0.095 |
| *Litochrus sp. 2* | Phalacridae | 1 | 1 | 0 | 0 | 1 | 0.137 | 0.111 | 0.024 | 0.429 | 0.021 | 0.015 | 0.004 | 0.061 |
| *Lygesis mendica* | Cerambycidae | 0 | 1 | 0 | 0 | 0 | 0.122 | 0.100 | 0.020 | 0.414 | 0.027 | 0.023 | 0.005 | 0.088 |
| *Mecyclothorax sp. 1 (ambiguus/punctipennis group)* | Carabidae | 0 | 0 | 1 | 0 | 0 | 0.109 | 0.084 | 0.017 | 0.341 | 0.028 | 0.023 | 0.005 | 0.086 |
| *Melanterius sp. 1* | Curculionidae | 0 | 1 | 0 | 0 | 0 | 0.108 | 0.085 | 0.019 | 0.353 | 0.029 | 0.026 | 0.005 | 0.097 |
| *Melobasis sp. 1 (pupurascens group)* | Buprestidae | 1 | 1 | 0 | 0 | 0 | 0.113 | 0.079 | 0.018 | 0.322 | 0.027 | 0.022 | 0.005 | 0.085 |
| *Metopum sp. 1* | Attelabidae | 0 | 1 | 0 | 0 | 0 | 0.091 | 0.073 | 0.015 | 0.286 | 0.046 | 0.037 | 0.008 | 0.145 |
| *Micraonychus sp. 1* | Curculionidae | 0 | 1 | 0 | 0 | 0 | 0.119 | 0.103 | 0.019 | 0.404 | 0.028 | 0.024 | 0.005 | 0.093 |
| *Micraspis furcifera* | Coccinellidae | 0 | 0 | 1 | 0 | 0 | 0.132 | 0.097 | 0.020 | 0.391 | 0.022 | 0.016 | 0.004 | 0.062 |
| *Misophrice sp. 1* | Curculionidae | 0 | 1 | 0 | 0 | 0 | 0.105 | 0.078 | 0.017 | 0.314 | 0.029 | 0.025 | 0.005 | 0.092 |
| *Misophrice sp. 2* | Curculionidae | 0 | 1 | 0 | 0 | 0 | 0.120 | 0.106 | 0.018 | 0.418 | 0.026 | 0.020 | 0.005 | 0.078 |
| *Monolepta picticollis* | Chrysomelidae | 0 | 1 | 0 | 0 | 0 | 0.118 | 0.103 | 0.016 | 0.386 | 0.027 | 0.022 | 0.005 | 0.083 |
| *Monolepta sp. 1 (froggatti group)* | Chrysomelidae | 0 | 1 | 0 | 0 | 0 | 0.076 | 0.054 | 0.016 | 0.229 | 0.092 | 0.064 | 0.017 | 0.265 |
| *Monolepta sp. 2 (christinae group)* | Chrysomelidae | 0 | 1 | 0 | 0 | 0 | 0.127 | 0.114 | 0.020 | 0.470 | 0.027 | 0.024 | 0.004 | 0.091 |
| *Monolepta sp. 4 (ordiaria group)* | Chrysomelidae | 0 | 1 | 0 | 0 | 0 | 0.125 | 0.099 | 0.019 | 0.395 | 0.023 | 0.017 | 0.004 | 0.069 |
| *Mordella sp. 1* | Mordellidae | 1 | 1 | 0 | 0 | 1 | 0.181 | 0.101 | 0.043 | 0.416 | 0.032 | 0.022 | 0.008 | 0.095 |
| *Mordella sp. 2* | Mordellidae | 1 | 1 | 0 | 0 | 1 | 0.114 | 0.086 | 0.018 | 0.351 | 0.027 | 0.022 | 0.005 | 0.090 |
| *Mordellistena sp. 1* | Mordellidae | 1 | 1 | 0 | 0 | 1 | 0.202 | 0.111 | 0.061 | 0.481 | 0.047 | 0.025 | 0.014 | 0.112 |
| *Neolaemosaccus chadwicki* | Curculionidae | 0 | 1 | 0 | 0 | 0 | 0.170 | 0.110 | 0.034 | 0.441 | 0.024 | 0.016 | 0.006 | 0.067 |
| *Nephus sp. 1* | Coccinellidae | 0 | 0 | 1 | 0 | 0 | 0.157 | 0.111 | 0.027 | 0.456 | 0.030 | 0.023 | 0.006 | 0.089 |
| *Nocar depressiusculus* | Tenebrionidae | 0 | 1 | 0 | 0 | 1 | 0.132 | 0.108 | 0.020 | 0.415 | 0.024 | 0.020 | 0.005 | 0.079 |
| *Notiobia sp. 1* | Carabidae | 0 | 0 | 1 | 0 | 0 | 0.115 | 0.088 | 0.019 | 0.348 | 0.029 | 0.025 | 0.005 | 0.096 |
| *Omonadus sp. 1 (hesperi group)* | Anthicidae | 0 | 1 | 1 | 0 | 1 | 0.178 | 0.105 | 0.045 | 0.451 | 0.044 | 0.029 | 0.010 | 0.121 |
| *Orthorhinus klugii* | Curculionidae | 0 | 1 | 0 | 0 | 0 | 0.249 | 0.140 | 0.064 | 0.587 | 0.037 | 0.024 | 0.009 | 0.100 |
| *Paratillus carus* | Cleridae | 1 | 1 | 1 | 0 | 0 | 0.118 | 0.094 | 0.017 | 0.386 | 0.027 | 0.024 | 0.005 | 0.090 |
| *Paropsides sp. 1* | Chrysomelidae | 0 | 1 | 0 | 0 | 0 | 0.084 | 0.063 | 0.015 | 0.252 | 0.048 | 0.039 | 0.008 | 0.152 |
| *Paropsisterna sp. 1* | Chrysomelidae | 0 | 1 | 0 | 0 | 0 | 0.109 | 0.079 | 0.019 | 0.320 | 0.029 | 0.025 | 0.005 | 0.098 |
| *Paropsisterna sp. 2* | Chrysomelidae | 0 | 1 | 0 | 0 | 0 | 0.131 | 0.107 | 0.018 | 0.439 | 0.022 | 0.016 | 0.004 | 0.066 |
| *Phalacrinus sp. 1 (corruscans group)* | Phalacridae | 1 | 1 | 0 | 0 | 1 | 0.342 | 0.143 | 0.124 | 0.669 | 0.032 | 0.014 | 0.012 | 0.065 |
| *Rhinophthalmus sp. 1* | Cerambycidae | 0 | 1 | 0 | 0 | 0 | 0.112 | 0.084 | 0.019 | 0.345 | 0.028 | 0.025 | 0.005 | 0.097 |
| *Rhyparida sp. 1* | Chrysomelidae | 0 | 1 | 0 | 0 | 0 | 0.103 | 0.085 | 0.016 | 0.311 | 0.029 | 0.026 | 0.005 | 0.096 |
| *Rhyzobius sp. 1* | Coccinellidae | 0 | 0 | 1 | 0 | 0 | 0.461 | 0.151 | 0.216 | 0.784 | 0.046 | 0.018 | 0.021 | 0.089 |
| *Rhyzobius sp. 2* | Coccinellidae | 0 | 0 | 1 | 0 | 0 | 0.334 | 0.143 | 0.119 | 0.652 | 0.040 | 0.021 | 0.013 | 0.091 |
| *Rhyzobius sp. 3* | Coccinellidae | 0 | 0 | 1 | 0 | 0 | 0.107 | 0.082 | 0.017 | 0.326 | 0.030 | 0.025 | 0.005 | 0.101 |
| *Rodolia sp. 1* | Coccinellidae | 0 | 0 | 1 | 0 | 0 | 0.123 | 0.094 | 0.019 | 0.374 | 0.023 | 0.017 | 0.005 | 0.069 |
| *Sarothrocrepis civica* | Carabidae | 0 | 0 | 1 | 0 | 0 | 0.116 | 0.091 | 0.022 | 0.357 | 0.024 | 0.019 | 0.004 | 0.073 |
| *Serangium maculigenum* | Coccinellidae | 0 | 0 | 1 | 0 | 0 | 0.443 | 0.103 | 0.265 | 0.653 | 0.099 | 0.024 | 0.058 | 0.151 |
| *Serangium mysticum* | Coccinellidae | 0 | 0 | 1 | 0 | 0 | 0.159 | 0.075 | 0.056 | 0.346 | 0.097 | 0.046 | 0.033 | 0.207 |
| *Staphylininae 1* | Staphylinidae | 0 | 0 | 1 | 0 | 0 | 0.120 | 0.091 | 0.021 | 0.370 | 0.023 | 0.017 | 0.005 | 0.068 |
| *Stethorus sp. 1* | Coccinellidae | 0 | 0 | 1 | 0 | 0 | 0.159 | 0.095 | 0.049 | 0.432 | 0.054 | 0.026 | 0.018 | 0.119 |
| *Symbothynus squalidus* | Curculionidae | 0 | 1 | 0 | 0 | 0 | 0.089 | 0.062 | 0.017 | 0.245 | 0.035 | 0.022 | 0.008 | 0.091 |
| *Syzeton sp. 1 (abnormis group)* | Aderidae | 0 | 1 | 0 | 0 | 1 | 0.101 | 0.078 | 0.017 | 0.324 | 0.029 | 0.024 | 0.005 | 0.093 |
| *Tarsostenus hilaris* | Cleridae | 1 | 1 | 1 | 0 | 0 | 0.155 | 0.103 | 0.031 | 0.415 | 0.028 | 0.021 | 0.006 | 0.083 |
| *Trachymela sp. 1* | Chrysomelidae | 0 | 1 | 0 | 0 | 0 | 0.122 | 0.084 | 0.022 | 0.353 | 0.042 | 0.028 | 0.010 | 0.116 |
| **Cicadas [Hemiptera: Auchenorrhyncha: Cicadomorpha: Cicadoidea]** | | | | | | | | | | | | | | |
| *Cicadidae 1* | Cicadidae | 0 | 1 | 0 | 0 | 0 | 0.116 | 0.090 | 0.020 | 0.356 | 0.026 | 0.020 | 0.005 | 0.079 |
| **Flies [Diptera: Brachycera]** | | | | | | | | | | | | | | |
| *Agromyzidae 1* | Agromyzidae | 0 | 1 | 0 | 0 | 0 | 0.219 | 0.114 | 0.074 | 0.518 | 0.062 | 0.032 | 0.017 | 0.138 |
| *Agromyzidae 2* | Agromyzidae | 0 | 1 | 0 | 0 | 0 | 0.394 | 0.123 | 0.189 | 0.661 | 0.059 | 0.019 | 0.030 | 0.103 |
| *Agromyzidae 3* | Agromyzidae | 0 | 1 | 0 | 0 | 0 | 0.205 | 0.101 | 0.065 | 0.457 | 0.044 | 0.024 | 0.013 | 0.104 |
| *Agromyzidae 4* | Agromyzidae | 0 | 1 | 0 | 0 | 0 | 0.197 | 0.115 | 0.052 | 0.500 | 0.035 | 0.018 | 0.011 | 0.080 |
| *Asilidae 1* | Asilidae | 0 | 0 | 1 | 0 | 0 | 0.128 | 0.098 | 0.021 | 0.398 | 0.024 | 0.020 | 0.004 | 0.077 |
| *Asilidae 2* | Asilidae | 0 | 0 | 1 | 0 | 0 | 0.204 | 0.108 | 0.052 | 0.465 | 0.037 | 0.021 | 0.011 | 0.090 |
| *Asilidae 3* | Asilidae | 0 | 0 | 1 | 0 | 0 | 0.146 | 0.090 | 0.029 | 0.368 | 0.029 | 0.021 | 0.006 | 0.085 |
| *Calliphoridae 1* | Calliphoridae | 1 | 0 | 0 | 0 | 1 | 0.111 | 0.082 | 0.020 | 0.325 | 0.028 | 0.021 | 0.005 | 0.087 |
| *Calliphoridae 2* | Calliphoridae | 1 | 0 | 0 | 0 | 1 | 0.237 | 0.123 | 0.064 | 0.542 | 0.028 | 0.016 | 0.008 | 0.069 |
| *Calliphoridae 3* | Calliphoridae | 1 | 0 | 0 | 0 | 1 | 0.314 | 0.128 | 0.129 | 0.614 | 0.054 | 0.025 | 0.020 | 0.115 |
| *Calliphoridae 4* | Calliphoridae | 1 | 0 | 0 | 0 | 1 | 0.210 | 0.143 | 0.046 | 0.619 | 0.042 | 0.033 | 0.009 | 0.135 |
| *Calliphoridae 5* | Calliphoridae | 1 | 0 | 0 | 0 | 1 | 0.173 | 0.116 | 0.031 | 0.465 | 0.024 | 0.016 | 0.005 | 0.065 |
| *Calliphoridae 6* | Calliphoridae | 1 | 0 | 0 | 0 | 1 | 0.104 | 0.076 | 0.017 | 0.309 | 0.028 | 0.023 | 0.005 | 0.088 |
| *Calliphoridae 7* | Calliphoridae | 1 | 0 | 0 | 0 | 1 | 0.116 | 0.093 | 0.019 | 0.357 | 0.029 | 0.027 | 0.005 | 0.103 |
| *Chloropidae 1* | Chloropidae | 0 | 0 | 0 | 0 | 1 | 0.312 | 0.129 | 0.119 | 0.597 | 0.036 | 0.016 | 0.014 | 0.076 |
| *Chloropidae 2* | Chloropidae | 0 | 0 | 0 | 0 | 1 | 0.271 | 0.109 | 0.104 | 0.515 | 0.043 | 0.019 | 0.018 | 0.088 |
| *Chloropidae 3* | Chloropidae | 0 | 0 | 0 | 0 | 1 | 0.235 | 0.131 | 0.057 | 0.547 | 0.030 | 0.019 | 0.008 | 0.081 |
| *Chloropidae 4* | Chloropidae | 0 | 0 | 0 | 0 | 1 | 0.482 | 0.147 | 0.233 | 0.810 | 0.040 | 0.014 | 0.018 | 0.073 |
| *Chloropidae 5* | Chloropidae | 0 | 0 | 0 | 0 | 1 | 0.321 | 0.158 | 0.103 | 0.700 | 0.043 | 0.025 | 0.012 | 0.108 |
| *Chloropidae 6* | Chloropidae | 0 | 0 | 0 | 0 | 1 | 0.395 | 0.147 | 0.164 | 0.725 | 0.035 | 0.014 | 0.015 | 0.070 |
| *Chloropidae 7* | Chloropidae | 0 | 0 | 0 | 0 | 1 | 0.315 | 0.122 | 0.134 | 0.617 | 0.059 | 0.021 | 0.027 | 0.106 |
| *Chloropidae 8* | Chloropidae | 0 | 0 | 0 | 0 | 1 | 0.588 | 0.134 | 0.329 | 0.839 | 0.037 | 0.012 | 0.019 | 0.065 |
| *Chloropidae 9* | Chloropidae | 0 | 0 | 0 | 0 | 1 | 0.328 | 0.139 | 0.127 | 0.645 | 0.038 | 0.018 | 0.013 | 0.084 |
| *Chloropidae 10* | Chloropidae | 0 | 0 | 0 | 0 | 1 | 0.105 | 0.075 | 0.017 | 0.297 | 0.029 | 0.024 | 0.005 | 0.092 |
| *Dolichopodidae 1* | Dolichopodidae | 0 | 0 | 1 | 0 | 0 | 0.133 | 0.093 | 0.026 | 0.383 | 0.034 | 0.027 | 0.006 | 0.105 |
| *Dolichopodidae 2* | Dolichopodidae | 0 | 0 | 1 | 0 | 0 | 0.329 | 0.156 | 0.104 | 0.691 | 0.036 | 0.018 | 0.012 | 0.080 |
| *Dolichopodidae 3* | Dolichopodidae | 0 | 0 | 1 | 0 | 0 | 0.450 | 0.143 | 0.203 | 0.769 | 0.039 | 0.014 | 0.018 | 0.072 |
| *Dolichopodidae 4* | Dolichopodidae | 0 | 0 | 1 | 0 | 0 | 0.108 | 0.087 | 0.019 | 0.354 | 0.030 | 0.027 | 0.005 | 0.105 |
| *Dolichopodidae 5* | Dolichopodidae | 0 | 0 | 1 | 0 | 0 | 0.235 | 0.145 | 0.057 | 0.622 | 0.030 | 0.019 | 0.008 | 0.078 |
| *Dolichopodidae 6* | Dolichopodidae | 0 | 0 | 1 | 0 | 0 | 0.110 | 0.089 | 0.016 | 0.343 | 0.030 | 0.026 | 0.005 | 0.095 |
| *Drosophilidae 1* | Drosophilidae | 0 | 0 | 0 | 0 | 1 | 0.337 | 0.150 | 0.105 | 0.669 | 0.027 | 0.014 | 0.009 | 0.062 |
| *Drosophilidae 2* | Drosophilidae | 0 | 0 | 0 | 0 | 1 | 0.147 | 0.095 | 0.031 | 0.391 | 0.028 | 0.019 | 0.006 | 0.077 |
| *Drosophilidae 3* | Drosophilidae | 0 | 0 | 0 | 0 | 1 | 0.260 | 0.147 | 0.065 | 0.622 | 0.025 | 0.014 | 0.007 | 0.059 |

| | | | | | | | | | | | | | | |
|---|---|---|---|---|---|---|---|---|---|---|---|---|---|---|
| *Drosophilidae 4* | Drosophilidae | 0 | 0 | 0 | 0 | 1 | 0.126 | 0.092 | 0.020 | 0.372 | 0.024 | 0.019 | 0.005 | 0.074 |
| *Empididae 1* | Empididae | 0 | 0 | 1 | 0 | 0 | 0.107 | 0.080 | 0.018 | 0.328 | 0.029 | 0.025 | 0.005 | 0.099 |
| *Empididae 2* | Empididae | 0 | 0 | 1 | 0 | 0 | 0.112 | 0.086 | 0.017 | 0.323 | 0.029 | 0.025 | 0.005 | 0.098 |
| *Ephydridae 1* | Ephydridae | 0 | 0 | 0 | 0 | 1 | 0.101 | 0.064 | 0.018 | 0.257 | 0.025 | 0.018 | 0.005 | 0.074 |
| *Ephydridae 2* | Ephydridae | 0 | 0 | 0 | 0 | 1 | 0.142 | 0.106 | 0.023 | 0.447 | 0.019 | 0.014 | 0.004 | 0.057 |
| *Ephydridae 3* | Ephydridae | 0 | 0 | 0 | 0 | 1 | 0.133 | 0.057 | 0.049 | 0.278 | 0.100 | 0.033 | 0.048 | 0.172 |
| *Ephydridae 4* | Ephydridae | 0 | 0 | 0 | 0 | 1 | 0.163 | 0.101 | 0.042 | 0.441 | 0.053 | 0.033 | 0.013 | 0.134 |
| *Ephydridae 5* | Ephydridae | 0 | 0 | 0 | 0 | 1 | 0.179 | 0.119 | 0.036 | 0.488 | 0.021 | 0.014 | 0.005 | 0.058 |
| *Ephydridae 6* | Ephydridae | 0 | 0 | 0 | 0 | 1 | 0.224 | 0.127 | 0.056 | 0.533 | 0.031 | 0.020 | 0.008 | 0.086 |
| *Heleomyzidae 1* | Heleomyzidae | 0 | 0 | 0 | 0 | 1 | 0.155 | 0.109 | 0.031 | 0.456 | 0.027 | 0.018 | 0.006 | 0.073 |
| *Hydrellia tritici* | Ephydridae | 1 | 1 | 0 | 0 | 1 | 0.809 | 0.047 | 0.713 | 0.892 | 0.344 | 0.022 | 0.302 | 0.387 |
| *Lauxaniidae 1* | Lauxaniidae | 0 | 0 | 0 | 0 | 1 | 0.517 | 0.090 | 0.356 | 0.711 | 0.132 | 0.022 | 0.092 | 0.178 |
| *Lauxaniidae 2* | Lauxaniidae | 0 | 0 | 0 | 0 | 1 | 0.747 | 0.087 | 0.574 | 0.901 | 0.092 | 0.014 | 0.067 | 0.122 |
| *Lauxaniidae 3* | Lauxaniidae | 0 | 0 | 0 | 0 | 1 | 0.323 | 0.152 | 0.091 | 0.659 | 0.023 | 0.013 | 0.008 | 0.055 |
| *Lauxaniidae 4* | Lauxaniidae | 0 | 0 | 0 | 0 | 1 | 0.263 | 0.101 | 0.117 | 0.525 | 0.078 | 0.030 | 0.031 | 0.149 |
| *Lauxaniidae 5* | Lauxaniidae | 0 | 0 | 0 | 0 | 1 | 0.353 | 0.127 | 0.144 | 0.644 | 0.049 | 0.017 | 0.024 | 0.091 |
| *Lauxaniidae 6* | Lauxaniidae | 0 | 0 | 0 | 0 | 1 | 0.128 | 0.058 | 0.047 | 0.271 | 0.109 | 0.040 | 0.045 | 0.199 |
| *Lauxaniidae 7* | Lauxaniidae | 0 | 0 | 0 | 0 | 1 | 0.301 | 0.085 | 0.170 | 0.499 | 0.109 | 0.026 | 0.063 | 0.166 |
| *Lauxaniidae 8* | Lauxaniidae | 0 | 0 | 0 | 0 | 1 | 0.305 | 0.130 | 0.102 | 0.595 | 0.030 | 0.016 | 0.011 | 0.073 |
| *Lauxaniidae 9* | Lauxaniidae | 0 | 0 | 0 | 0 | 1 | 0.137 | 0.088 | 0.030 | 0.367 | 0.031 | 0.023 | 0.007 | 0.096 |
| *Lauxaniidae 10* | Lauxaniidae | 0 | 0 | 0 | 0 | 1 | 0.138 | 0.092 | 0.027 | 0.380 | 0.033 | 0.026 | 0.007 | 0.102 |
| *Lauxaniidae 11* | Lauxaniidae | 0 | 0 | 0 | 0 | 1 | 0.115 | 0.083 | 0.017 | 0.335 | 0.025 | 0.021 | 0.005 | 0.083 |
| *Lonchopteridae 1* | Lonchopteridae | 0 | 0 | 0 | 0 | 1 | 0.434 | 0.144 | 0.199 | 0.769 | 0.046 | 0.019 | 0.019 | 0.091 |
| *Muscidae 1* | Muscidae | 1 | 0 | 0 | 0 | 1 | 0.622 | 0.080 | 0.476 | 0.793 | 0.178 | 0.021 | 0.137 | 0.222 |
| *Muscidae 2* | Muscidae | 1 | 0 | 0 | 0 | 1 | 0.293 | 0.148 | 0.083 | 0.656 | 0.028 | 0.015 | 0.009 | 0.069 |
| *Muscidae 3* | Muscidae | 1 | 0 | 0 | 0 | 1 | 0.581 | 0.119 | 0.356 | 0.822 | 0.056 | 0.013 | 0.033 | 0.086 |
| *Muscidae 4* | Muscidae | 1 | 0 | 0 | 0 | 1 | 0.244 | 0.123 | 0.081 | 0.552 | 0.052 | 0.031 | 0.015 | 0.126 |
| *Muscidae 5* | Muscidae | 1 | 0 | 0 | 0 | 1 | 0.112 | 0.089 | 0.017 | 0.346 | 0.029 | 0.025 | 0.004 | 0.094 |
| *Muscidae 6* | Muscidae | 1 | 0 | 0 | 0 | 1 | 0.194 | 0.118 | 0.047 | 0.494 | 0.039 | 0.027 | 0.010 | 0.109 |
| *Muscidae 7* | Muscidae | 1 | 0 | 0 | 0 | 1 | 0.211 | 0.123 | 0.054 | 0.538 | 0.037 | 0.022 | 0.011 | 0.093 |
| *Muscidae 8* | Muscidae | 1 | 0 | 0 | 0 | 1 | 0.125 | 0.089 | 0.021 | 0.357 | 0.024 | 0.019 | 0.005 | 0.077 |
| *Muscidae 9* | Muscidae | 1 | 0 | 0 | 0 | 1 | 0.130 | 0.086 | 0.026 | 0.347 | 0.035 | 0.027 | 0.007 | 0.105 |
| *Muscidae 10* | Muscidae | 1 | 0 | 0 | 0 | 1 | 0.140 | 0.120 | 0.019 | 0.492 | 0.022 | 0.017 | 0.004 | 0.068 |
| *Parentia sp. 1* | Dolichopodidae | 0 | 0 | 1 | 0 | 0 | 0.358 | 0.141 | 0.127 | 0.688 | 0.039 | 0.019 | 0.014 | 0.087 |
| *Phoridae 1* | Phoridae | 1 | 1 | 0 | 0 | 1 | 0.476 | 0.146 | 0.233 | 0.793 | 0.046 | 0.015 | 0.022 | 0.082 |
| *Phoridae 2* | Phoridae | 1 | 1 | 0 | 0 | 1 | 0.181 | 0.112 | 0.042 | 0.474 | 0.031 | 0.020 | 0.007 | 0.083 |
| *Phoridae 3* | Phoridae | 1 | 1 | 0 | 0 | 1 | 0.154 | 0.102 | 0.035 | 0.429 | 0.044 | 0.029 | 0.010 | 0.119 |
| *Phoridae 4* | Phoridae | 1 | 1 | 0 | 0 | 1 | 0.410 | 0.122 | 0.217 | 0.689 | 0.058 | 0.018 | 0.030 | 0.097 |
| *Phoridae 5* | Phoridae | 1 | 1 | 0 | 0 | 1 | 0.189 | 0.110 | 0.046 | 0.465 | 0.028 | 0.017 | 0.007 | 0.073 |
| *Phoridae 6* | Phoridae | 1 | 1 | 0 | 0 | 1 | 0.337 | 0.153 | 0.128 | 0.695 | 0.037 | 0.019 | 0.013 | 0.086 |
| *Phoridae 7* | Phoridae | 1 | 1 | 0 | 0 | 1 | 0.119 | 0.098 | 0.019 | 0.380 | 0.027 | 0.021 | 0.005 | 0.085 |
| *Pipunculidae 1* | Pipunculidae | 1 | 0 | 1 | 1 | 0 | 0.239 | 0.121 | 0.068 | 0.535 | 0.036 | 0.019 | 0.011 | 0.083 |
| *Pipunculidae 2* | Platypezidae | 1 | 0 | 1 | 1 | 0 | 0.167 | 0.123 | 0.029 | 0.490 | 0.027 | 0.020 | 0.006 | 0.082 |
| *Pyrgotidae 1* | Pyrgotidae | 0 | 0 | 1 | 1 | 0 | 0.191 | 0.111 | 0.048 | 0.486 | 0.028 | 0.018 | 0.007 | 0.075 |
| *Pyrgotidae 2* | Pyrgotidae | 0 | 0 | 1 | 1 | 0 | 0.130 | 0.091 | 0.022 | 0.361 | 0.022 | 0.017 | 0.004 | 0.064 |
| *Rivellia sp. 1* | Platystomatidae | 1 | 0 | 0 | 0 | 1 | 0.390 | 0.112 | 0.207 | 0.630 | 0.073 | 0.020 | 0.040 | 0.118 |
| *Rivellia sp. 2* | Platystomatidae | 1 | 0 | 0 | 0 | 1 | 0.377 | 0.153 | 0.152 | 0.723 | 0.032 | 0.013 | 0.013 | 0.062 |
| *Rivellia sp. 3* | Platystomatidae | 1 | 0 | 0 | 0 | 1 | 0.284 | 0.152 | 0.080 | 0.662 | 0.022 | 0.012 | 0.007 | 0.051 |
| *Scenopinidae 1* | Scenopinidae | 0 | 0 | 1 | 0 | 0 | 0.145 | 0.081 | 0.042 | 0.345 | 0.063 | 0.031 | 0.020 | 0.140 |
| *Scenopinidae 2* | Scenopinidae | 0 | 0 | 1 | 0 | 0 | 0.179 | 0.120 | 0.036 | 0.501 | 0.023 | 0.015 | 0.005 | 0.061 |
| *Sepsidae 1* | Sepsidae | 0 | 0 | 0 | 0 | 1 | 0.412 | 0.141 | 0.192 | 0.754 | 0.047 | 0.018 | 0.021 | 0.090 |
| *Stratiomyidae 1* | Stratiomyidae | 1 | 0 | 0 | 0 | 1 | 0.226 | 0.131 | 0.057 | 0.583 | 0.031 | 0.019 | 0.008 | 0.079 |
| *Stratiomyidae 2* | Stratiomyidae | 1 | 0 | 0 | 0 | 1 | 0.489 | 0.153 | 0.224 | 0.799 | 0.034 | 0.013 | 0.015 | 0.065 |
| *Stratiomyidae 3* | Stratiomyidae | 1 | 0 | 0 | 0 | 1 | 0.115 | 0.087 | 0.020 | 0.360 | 0.025 | 0.020 | 0.005 | 0.077 |
| *Syrphidae 1* | Syrphidae | 1 | 1 | 0 | 0 | 0 | 0.257 | 0.129 | 0.076 | 0.562 | 0.042 | 0.025 | 0.011 | 0.106 |
| *Syrphidae 2* | Syrphidae | 1 | 1 | 0 | 0 | 0 | 0.116 | 0.093 | 0.019 | 0.379 | 0.026 | 0.021 | 0.005 | 0.080 |
| *Syrphidae 3* | Syrphidae | 1 | 1 | 0 | 0 | 0 | 0.133 | 0.109 | 0.020 | 0.430 | 0.026 | 0.023 | 0.005 | 0.086 |
| *Tachinidae 1* | Tachinidae | 1 | 0 | 1 | 1 | 0 | 0.118 | 0.085 | 0.021 | 0.340 | 0.023 | 0.017 | 0.005 | 0.066 |
| *Tachinidae 2* | Tachinidae | 1 | 0 | 1 | 1 | 0 | 0.122 | 0.094 | 0.018 | 0.364 | 0.026 | 0.021 | 0.004 | 0.079 |
| *Tachinidae 3* | Tachinidae | 1 | 0 | 1 | 1 | 0 | 0.107 | 0.080 | 0.017 | 0.325 | 0.026 | 0.021 | 0.005 | 0.083 |
| *Tachinidae 4* | Tachinidae | 1 | 0 | 1 | 1 | 0 | 0.262 | 0.140 | 0.067 | 0.588 | 0.024 | 0.013 | 0.007 | 0.056 |
| *Tachinidae 5* | Tachinidae | 1 | 0 | 1 | 1 | 0 | 0.158 | 0.120 | 0.028 | 0.528 | 0.029 | 0.021 | 0.006 | 0.087 |
| *Tachinidae 6* | Tachinidae | 1 | 0 | 1 | 1 | 0 | 0.099 | 0.075 | 0.018 | 0.287 | 0.029 | 0.024 | 0.005 | 0.094 |
| *Tephritidae 1* | Tephritidae | 0 | 1 | 0 | 0 | 0 | 0.203 | 0.084 | 0.079 | 0.408 | 0.078 | 0.029 | 0.034 | 0.146 |
| *Tephritidae 2* | Tephritidae | 0 | 1 | 0 | 0 | 0 | 0.121 | 0.095 | 0.021 | 0.403 | 0.025 | 0.019 | 0.004 | 0.075 |
| *Tephritidae 3* | Tephritidae | 0 | 1 | 0 | 0 | 0 | 0.122 | 0.089 | 0.023 | 0.353 | 0.023 | 0.018 | 0.004 | 0.069 |
| *Tephritidae 4* | Tephritidae | 0 | 1 | 0 | 0 | 0 | 0.149 | 0.093 | 0.033 | 0.384 | 0.027 | 0.019 | 0.006 | 0.079 |
| *Tephritidae 5* | Tephritidae | 0 | 1 | 0 | 0 | 0 | 0.128 | 0.102 | 0.019 | 0.430 | 0.024 | 0.018 | 0.004 | 0.072 |
| *Tephritidae 6* | Tephritidae | 0 | 1 | 0 | 0 | 0 | 0.119 | 0.093 | 0.017 | 0.373 | 0.027 | 0.024 | 0.005 | 0.088 |
| *Tephritidae 7* | Tephritidae | 0 | 1 | 0 | 0 | 0 | 0.113 | 0.088 | 0.019 | 0.342 | 0.025 | 0.020 | 0.005 | 0.076 |
| *Tephritidae 8* | Tephritidae | 0 | 1 | 0 | 0 | 0 | 0.292 | 0.142 | 0.090 | 0.632 | 0.025 | 0.013 | 0.009 | 0.056 |
| *Therevidae 1* | Therevidae | 0 | 0 | 1 | 0 | 0 | 0.289 | 0.119 | 0.112 | 0.558 | 0.040 | 0.016 | 0.017 | 0.079 |
| *Therevidae 2* | Therevidae | 0 | 0 | 1 | 0 | 0 | 0.274 | 0.122 | 0.085 | 0.549 | 0.029 | 0.014 | 0.010 | 0.063 |
| *Therevidae 3* | Therevidae | 0 | 0 | 1 | 0 | 0 | 0.157 | 0.113 | 0.030 | 0.456 | 0.028 | 0.021 | 0.005 | 0.083 |
| **Heteropteran bugs [Hempitera: Heteroptera]** | | | | | | | | | | | | | | |
| *Brentiscerus putoni* | Rhyparochromidae | 0 | 1 | 0 | 0 | 0 | 0.173 | 0.108 | 0.037 | 0.458 | 0.023 | 0.015 | 0.005 | 0.062 |
| *Campylomma liebknechti* | Miridae | 0 | 1 | 0 | 0 | 0 | 0.233 | 0.087 | 0.108 | 0.443 | 0.095 | 0.033 | 0.043 | 0.169 |
| *Cermatulus nasalis* | Pentatomidae | 0 | 0 | 1 | 0 | 0 | 0.123 | 0.099 | 0.017 | 0.394 | 0.028 | 0.025 | 0.005 | 0.096 |
| *Coridromius sp. 1* | Miridae | 0 | 1 | 0 | 0 | 0 | 0.078 | 0.029 | 0.030 | 0.143 | 0.239 | 0.055 | 0.140 | 0.355 |
| *Creontiades dilutus* | Miridae | 0 | 1 | 0 | 0 | 0 | 0.183 | 0.109 | 0.044 | 0.472 | 0.043 | 0.031 | 0.010 | 0.125 |
| *Crompus oculatus* | Lygaeidae | 0 | 1 | 0 | 0 | 0 | 0.066 | 0.050 | 0.012 | 0.199 | 0.072 | 0.051 | 0.014 | 0.206 |
| *Cryptorhamphidae 1* | Cryptorhamphidae | 0 | 1 | 0 | 0 | 0 | 0.066 | 0.046 | 0.013 | 0.193 | 0.062 | 0.034 | 0.017 | 0.148 |
| *Cuspicona sp. 1* | Pentatomidae | 0 | 1 | 0 | 0 | 0 | 0.101 | 0.072 | 0.017 | 0.298 | 0.028 | 0.022 | 0.005 | 0.088 |
| *Cuspicona sp. 2* | Pentatomidae | 0 | 1 | 0 | 0 | 0 | 0.084 | 0.061 | 0.015 | 0.243 | 0.043 | 0.032 | 0.008 | 0.128 |
| *Daerlac sp. 1* | Rhyparochromidae | 0 | 1 | 0 | 0 | 0 | 0.114 | 0.085 | 0.019 | 0.334 | 0.022 | 0.016 | 0.005 | 0.066 |
| *Dictyotus sp. 1* | Pentatomidae | 0 | 1 | 0 | 0 | 0 | 0.126 | 0.092 | 0.021 | 0.370 | 0.022 | 0.016 | 0.004 | 0.064 |
| *Dilompus sp. 1* | Artheneidae | 0 | 1 | 0 | 0 | 0 | 0.082 | 0.074 | 0.013 | 0.274 | 0.048 | 0.037 | 0.009 | 0.148 |
| *Emesinae 1* | Reduviidae | 0 | 0 | 1 | 0 | 0 | 0.105 | 0.065 | 0.025 | 0.270 | 0.049 | 0.034 | 0.011 | 0.142 |
| *Eribotes sp. 1* | Pentatomidae | 0 | 1 | 0 | 0 | 0 | 0.194 | 0.117 | 0.044 | 0.503 | 0.029 | 0.019 | 0.007 | 0.079 |
| *Eritingis sp. 1 (trivigata group)* | Tingidae | 0 | 1 | 0 | 0 | 0 | 0.128 | 0.114 | 0.018 | 0.468 | 0.028 | 0.026 | 0.004 | 0.099 |
| *Eurynysius meschioides* | Lygaeidae | 0 | 1 | 0 | 0 | 0 | 0.111 | 0.094 | 0.017 | 0.355 | 0.027 | 0.021 | 0.005 | 0.080 |
| *Eysarcoris sp. 1* | Pentatomidae | 0 | 1 | 0 | 0 | 0 | 0.162 | 0.112 | 0.032 | 0.474 | 0.026 | 0.017 | 0.006 | 0.070 |
| *Froggiatta olivinia* | Tingidae | 0 | 1 | 0 | 0 | 0 | 0.105 | 0.075 | 0.018 | 0.303 | 0.030 | 0.025 | 0.005 | 0.095 |
| *Germalus victoriae* | Geocoridae | 0 | 0 | 1 | 0 | 0 | 0.166 | 0.091 | 0.045 | 0.401 | 0.049 | 0.025 | 0.015 | 0.110 |
| *Heinsius sp. 1* | Blissidae | 0 | 1 | 0 | 0 | 0 | 0.069 | 0.053 | 0.013 | 0.213 | 0.061 | 0.035 | 0.016 | 0.151 |
| *Iphicrates spathus* | Blissidae | 0 | 1 | 0 | 0 | 0 | 0.104 | 0.070 | 0.021 | 0.291 | 0.052 | 0.030 | 0.013 | 0.127 |
| *Malandiola sp. 1 (semota group)* | Tingidae | 0 | 1 | 0 | 0 | 0 | 0.136 | 0.111 | 0.020 | 0.429 | 0.022 | 0.016 | 0.004 | 0.066 |
| *Mcateella sp. 1* | Piesmatidae | 0 | 1 | 0 | 0 | 0 | 0.099 | 0.074 | 0.015 | 0.304 | 0.029 | 0.024 | 0.005 | 0.092 |
| *Miridae 1* | Miridae | 0 | 1 | 0 | 0 | 0 | 0.190 | 0.113 | 0.047 | 0.480 | 0.042 | 0.024 | 0.012 | 0.105 |
| *Miridae 2* | Miridae | 0 | 1 | 0 | 0 | 0 | 0.121 | 0.100 | 0.019 | 0.404 | 0.028 | 0.023 | 0.005 | 0.088 |
| *Miridae 3* | Miridae | 0 | 1 | 0 | 0 | 0 | 0.044 | 0.035 | 0.009 | 0.131 | 0.138 | 0.091 | 0.028 | 0.376 |
| *Miridae 4* | Miridae | 0 | 1 | 0 | 0 | 0 | 0.049 | 0.040 | 0.009 | 0.155 | 0.133 | 0.093 | 0.025 | 0.368 |
| *Miridae 5* | Miridae | 0 | 1 | 0 | 0 | 0 | 0.124 | 0.099 | 0.018 | 0.409 | 0.026 | 0.022 | 0.005 | 0.082 |

| | | | | | | | | | | | | | | |
|---|---|---|---|---|---|---|---|---|---|---|---|---|---|---|
| *Mirinae 1* | Miridae | 0 | 1 | 0 | 0 | 0 | 0.232 | 0.128 | 0.062 | 0.547 | 0.029 | 0.017 | 0.008 | 0.072 |
| *Mirinae 2* | Miridae | 0 | 1 | 0 | 0 | 0 | 0.065 | 0.050 | 0.012 | 0.201 | 0.080 | 0.060 | 0.015 | 0.239 |
| *Mirinae 3* | Miridae | 0 | 1 | 0 | 0 | 0 | 0.134 | 0.113 | 0.019 | 0.461 | 0.026 | 0.022 | 0.004 | 0.081 |
| *Mirinae 4* | Miridae | 0 | 1 | 0 | 0 | 0 | 0.104 | 0.077 | 0.018 | 0.313 | 0.027 | 0.023 | 0.005 | 0.087 |
| *Mirinae 5* | Miridae | 0 | 1 | 0 | 0 | 0 | 0.120 | 0.094 | 0.019 | 0.383 | 0.027 | 0.023 | 0.005 | 0.089 |
| *Mutusca brevicornis* | Alydidae | 0 | 1 | 0 | 0 | 0 | 0.087 | 0.043 | 0.028 | 0.185 | 0.121 | 0.039 | 0.056 | 0.209 |
| *Nabis kinbergii* | Nabidae | 0 | 0 | 1 | 0 | 0 | 0.274 | 0.097 | 0.129 | 0.514 | 0.093 | 0.031 | 0.043 | 0.163 |
| *Nysius caledoniae* | Lygaeidae | 1 | 1 | 0 | 0 | 0 | 0.099 | 0.062 | 0.026 | 0.258 | 0.078 | 0.039 | 0.022 | 0.173 |
| *Nysius vinitor* | Lygaeidae | 1 | 1 | 0 | 0 | 0 | 0.669 | 0.093 | 0.489 | 0.845 | 0.123 | 0.019 | 0.092 | 0.163 |
| *Oechalia schellenbergii* | Pentatomidae | 0 | 0 | 1 | 0 | 0 | 0.115 | 0.094 | 0.017 | 0.368 | 0.028 | 0.024 | 0.005 | 0.093 |
| *Paromyocara punctatum* | Rhyparochromidae | 0 | 1 | 0 | 0 | 0 | 0.084 | 0.068 | 0.013 | 0.280 | 0.053 | 0.046 | 0.009 | 0.181 |
| *Pentatomidae 1* | Pentatomidae | 0 | 1 | 0 | 0 | 0 | 0.130 | 0.100 | 0.022 | 0.400 | 0.021 | 0.016 | 0.004 | 0.060 |
| *Pentatomidae 2* | Pentatomidae | 0 | 1 | 0 | 0 | 0 | 0.103 | 0.068 | 0.021 | 0.275 | 0.059 | 0.048 | 0.011 | 0.187 |
| *Pentatomidae 3* | Pentatomidae | 0 | 1 | 0 | 0 | 0 | 0.183 | 0.111 | 0.040 | 0.466 | 0.034 | 0.020 | 0.009 | 0.086 |
| *Pentatomidae 4* | Pentatomidae | 0 | 1 | 0 | 0 | 0 | 0.132 | 0.107 | 0.018 | 0.420 | 0.026 | 0.023 | 0.004 | 0.090 |
| *Pentatomidae 5* | Pentatomidae | 0 | 1 | 0 | 0 | 0 | 0.108 | 0.085 | 0.017 | 0.342 | 0.030 | 0.020 | 0.006 | 0.081 |
| *Pentatomidae 6* | Pentatomidae | 0 | 1 | 0 | 0 | 0 | 0.152 | 0.103 | 0.034 | 0.429 | 0.028 | 0.017 | 0.008 | 0.072 |
| *Phylinae 1* | Miridae | 0 | 1 | 0 | 0 | 0 | 0.075 | 0.057 | 0.013 | 0.223 | 0.051 | 0.040 | 0.009 | 0.161 |
| *Phylinae 2* | Miridae | 0 | 1 | 0 | 0 | 0 | 0.118 | 0.099 | 0.020 | 0.383 | 0.027 | 0.023 | 0.005 | 0.087 |
| *Phylinae 3* | Miridae | 0 | 1 | 0 | 0 | 0 | 0.178 | 0.100 | 0.042 | 0.416 | 0.034 | 0.024 | 0.008 | 0.099 |
| *Phylinae 4* | Miridae | 0 | 1 | 0 | 0 | 0 | 0.086 | 0.075 | 0.014 | 0.267 | 0.053 | 0.046 | 0.009 | 0.176 |
| *Plautia sp. 1* | Pentatomidae | 0 | 1 | 0 | 0 | 0 | 0.107 | 0.077 | 0.018 | 0.313 | 0.025 | 0.019 | 0.005 | 0.080 |
| *Plinthisus woodwardi* | Rhyparochromidae | 0 | 1 | 0 | 0 | 0 | 0.252 | 0.128 | 0.074 | 0.571 | 0.033 | 0.020 | 0.009 | 0.083 |
| *Poecilometis sp. 1* | Pentatomidae | 0 | 1 | 0 | 0 | 0 | 0.105 | 0.078 | 0.021 | 0.315 | 0.055 | 0.041 | 0.011 | 0.162 |
| *Rayieria sp. 1* | Miridae | 0 | 1 | 0 | 0 | 0 | 0.121 | 0.104 | 0.018 | 0.418 | 0.027 | 0.022 | 0.004 | 0.086 |
| *Remaudiereana inornata* | Rhyparochromidae | 0 | 1 | 0 | 0 | 0 | 0.289 | 0.143 | 0.097 | 0.643 | 0.050 | 0.026 | 0.017 | 0.113 |
| *Sidnia kinbergii* | Miridae | 0 | 1 | 0 | 0 | 0 | 0.197 | 0.116 | 0.057 | 0.512 | 0.053 | 0.032 | 0.014 | 0.133 |
| *Spilostethus hospes* | Lygaeidae | 1 | 1 | 0 | 0 | 0 | 0.110 | 0.084 | 0.017 | 0.330 | 0.029 | 0.026 | 0.005 | 0.095 |
| *Stenophylla macreta* | Pachygronthidae | 0 | 1 | 0 | 0 | 0 | 0.127 | 0.056 | 0.047 | 0.270 | 0.109 | 0.034 | 0.052 | 0.183 |
| *Stizocephalus sp. 1* | Rhyparochromidae | 0 | 1 | 0 | 0 | 0 | 0.127 | 0.096 | 0.021 | 0.388 | 0.023 | 0.017 | 0.004 | 0.071 |
| *Taylorilygus sp. 1* | Miridae | 0 | 1 | 0 | 0 | 0 | 0.277 | 0.064 | 0.168 | 0.418 | 0.163 | 0.034 | 0.102 | 0.235 |
| *Theseus sp. 1* | Pentatomidae | 0 | 1 | 0 | 0 | 0 | 0.167 | 0.129 | 0.032 | 0.561 | 0.027 | 0.020 | 0.005 | 0.080 |
| *Tingis sp. nov.* | Tingidae | 0 | 1 | 0 | 0 | 0 | 0.080 | 0.058 | 0.017 | 0.227 | 0.089 | 0.061 | 0.019 | 0.249 |

**Jumping plant lice [Hemiptera: Sternorrhyncha: Psylloidea]**

| | | | | | | | | | | | | | | |
|---|---|---|---|---|---|---|---|---|---|---|---|---|---|---|
| *Aacanthocnema dobsoni* | Triozidae | 0 | 1 | 0 | 0 | 0 | 0.175 | 0.108 | 0.041 | 0.429 | 0.032 | 0.018 | 0.009 | 0.079 |
| *Acanthocasuarina sp. nov. (muellerianae group)* | Triozidae | 0 | 1 | 0 | 0 | 0 | 0.228 | 0.138 | 0.050 | 0.579 | 0.023 | 0.013 | 0.006 | 0.057 |
| *Acizzia jucunda* | Psyllidae | 0 | 1 | 0 | 0 | 0 | 0.036 | 0.024 | 0.008 | 0.100 | 0.205 | 0.117 | 0.047 | 0.490 |
| *Acizzia sp. 1* | Psyllidae | 0 | 1 | 0 | 0 | 0 | 0.113 | 0.084 | 0.020 | 0.337 | 0.029 | 0.023 | 0.005 | 0.092 |
| *Acizzia sp. 2* | Psyllidae | 0 | 1 | 0 | 0 | 0 | 0.157 | 0.103 | 0.037 | 0.421 | 0.026 | 0.018 | 0.006 | 0.075 |
| *Acizzia sp. 3* | Psyllidae | 0 | 1 | 0 | 0 | 0 | 0.080 | 0.058 | 0.014 | 0.240 | 0.054 | 0.045 | 0.009 | 0.172 |
| *Acizzia sp. 4* | Psyllidae | 0 | 1 | 0 | 0 | 0 | 0.109 | 0.082 | 0.019 | 0.347 | 0.029 | 0.025 | 0.005 | 0.095 |
| *Acizzia sp. 5 (conspicua group)* | Psyllidae | 0 | 1 | 0 | 0 | 0 | 0.167 | 0.098 | 0.034 | 0.403 | 0.025 | 0.017 | 0.006 | 0.069 |
| *Agelaeopsylla sp. 1 (dividua group)* | Psyllidae | 0 | 1 | 0 | 0 | 0 | 0.104 | 0.079 | 0.017 | 0.324 | 0.030 | 0.025 | 0.005 | 0.095 |
| *Anoeconessa sp. 1* | Psyllidae | 0 | 1 | 0 | 0 | 0 | 0.112 | 0.090 | 0.016 | 0.368 | 0.027 | 0.022 | 0.005 | 0.088 |
| *Anoeconessa unicornuta* | Psyllidae | 0 | 1 | 0 | 0 | 0 | 0.178 | 0.110 | 0.038 | 0.458 | 0.020 | 0.013 | 0.005 | 0.054 |
| *Blastopsylla sp. 1* | Psyllidae | 0 | 1 | 0 | 0 | 0 | 0.139 | 0.108 | 0.021 | 0.428 | 0.020 | 0.014 | 0.004 | 0.058 |
| *Boreioglycaspis australiensis* | Psyllidae | 0 | 1 | 0 | 0 | 0 | 0.093 | 0.072 | 0.015 | 0.293 | 0.045 | 0.038 | 0.008 | 0.148 |
| *Cardiaspina retator* | Psyllidae | 0 | 1 | 0 | 0 | 0 | 0.119 | 0.096 | 0.019 | 0.381 | 0.028 | 0.023 | 0.005 | 0.091 |
| *Cardiaspina sp. 1 (albitextura group)* | Psyllidae | 0 | 1 | 0 | 0 | 0 | 0.161 | 0.093 | 0.038 | 0.404 | 0.041 | 0.022 | 0.013 | 0.096 |
| *Cardiaspina sp. 2 (albicollaris group)* | Psyllidae | 0 | 1 | 0 | 0 | 0 | 0.117 | 0.091 | 0.017 | 0.361 | 0.024 | 0.018 | 0.005 | 0.074 |
| *Cardiaspina sp. 3* | Psyllidae | 0 | 1 | 0 | 0 | 0 | 0.106 | 0.084 | 0.016 | 0.330 | 0.029 | 0.024 | 0.005 | 0.095 |
| *Creiis sp. 1* | Psyllidae | 0 | 1 | 0 | 0 | 0 | 0.119 | 0.101 | 0.020 | 0.391 | 0.024 | 0.017 | 0.005 | 0.069 |
| *Cryptoneossa sp. 1 (vulgaris group)* | Psyllidae | 0 | 1 | 0 | 0 | 0 | 0.140 | 0.107 | 0.027 | 0.431 | 0.036 | 0.031 | 0.006 | 0.119 |
| *Cryptoneossa triangula* | Psyllidae | 0 | 1 | 0 | 0 | 0 | 0.125 | 0.080 | 0.034 | 0.345 | 0.081 | 0.049 | 0.020 | 0.202 |
| *Ctenarytaina longicauda* | Psyllidae | 0 | 1 | 0 | 0 | 0 | 0.136 | 0.078 | 0.036 | 0.333 | 0.046 | 0.028 | 0.012 | 0.117 |
| *Ctenarytaina sp. 1* | Psyllidae | 0 | 1 | 0 | 0 | 0 | 0.253 | 0.132 | 0.083 | 0.582 | 0.044 | 0.021 | 0.015 | 0.098 |
| *Ctenarytaina spatulata* | Psyllidae | 0 | 1 | 0 | 0 | 0 | 0.116 | 0.103 | 0.017 | 0.399 | 0.029 | 0.026 | 0.005 | 0.101 |
| *Dasypsylla sp. 1* | Psyllidae | 0 | 1 | 0 | 0 | 0 | 0.118 | 0.086 | 0.018 | 0.348 | 0.025 | 0.020 | 0.005 | 0.075 |
| *Eucalyptolyma maideni* | Psyllidae | 0 | 1 | 0 | 0 | 0 | 0.105 | 0.076 | 0.020 | 0.319 | 0.056 | 0.040 | 0.011 | 0.161 |
| *Glycaspis sp. 1 (brimblecombei group)* | Psyllidae | 0 | 1 | 0 | 0 | 0 | 0.047 | 0.032 | 0.012 | 0.125 | 0.204 | 0.108 | 0.053 | 0.456 |
| *Mycopsylla sp. 1 (fici group)* | Homotomidae | 0 | 1 | 0 | 0 | 0 | 0.046 | 0.034 | 0.010 | 0.123 | 0.123 | 0.067 | 0.032 | 0.289 |
| *Mycopsylla sp. nov. (tuberculata group)* | Homotomidae | 0 | 1 | 0 | 0 | 0 | 0.060 | 0.051 | 0.011 | 0.206 | 0.085 | 0.052 | 0.018 | 0.215 |
| *Phellopsylla sp. 1* | Psyllidae | 0 | 1 | 0 | 0 | 0 | 0.086 | 0.047 | 0.025 | 0.205 | 0.093 | 0.044 | 0.030 | 0.194 |
| *Phyllolyma sp. 1 (rufa group)* | Psyllidae | 0 | 1 | 0 | 0 | 0 | 0.124 | 0.088 | 0.020 | 0.350 | 0.022 | 0.016 | 0.005 | 0.066 |
| *Platyobria biemani* | Psyllidae | 0 | 1 | 0 | 0 | 0 | 0.127 | 0.093 | 0.021 | 0.369 | 0.020 | 0.014 | 0.005 | 0.056 |

**Leafhoppers/Treehoppers [Hemiptera: Auchenorrhyncha: Cicadomorpha: Membracoidea]**

| | | | | | | | | | | | | | | |
|---|---|---|---|---|---|---|---|---|---|---|---|---|---|---|
| *Cicadellidae 1* | Cicadellidae | 0 | 1 | 0 | 0 | 0 | 0.114 | 0.084 | 0.018 | 0.331 | 0.029 | 0.025 | 0.005 | 0.096 |
| *Cicadellidae 2* | Cicadellidae | 0 | 1 | 0 | 0 | 0 | 0.074 | 0.058 | 0.013 | 0.221 | 0.059 | 0.033 | 0.015 | 0.138 |
| *Cicadellidae 3* | Cicadellidae | 0 | 1 | 0 | 0 | 0 | 0.053 | 0.035 | 0.011 | 0.147 | 0.088 | 0.052 | 0.022 | 0.223 |
| *Cicadellidae 4* | Cicadellidae | 0 | 1 | 0 | 0 | 0 | 0.120 | 0.086 | 0.021 | 0.358 | 0.024 | 0.018 | 0.004 | 0.071 |
| *Cicadellidae 5* | Cicadellidae | 0 | 1 | 0 | 0 | 0 | 0.116 | 0.090 | 0.019 | 0.345 | 0.024 | 0.018 | 0.005 | 0.071 |
| *Cicadellidae 6* | Cicadellidae | 0 | 1 | 0 | 0 | 0 | 0.102 | 0.081 | 0.017 | 0.311 | 0.030 | 0.026 | 0.005 | 0.098 |
| *Cicadellidae 7* | Cicadellidae | 0 | 1 | 0 | 0 | 0 | 0.083 | 0.080 | 0.012 | 0.334 | 0.055 | 0.047 | 0.008 | 0.187 |
| *Cicadellidae 8* | Cicadellidae | 0 | 1 | 0 | 0 | 0 | 0.126 | 0.118 | 0.018 | 0.499 | 0.028 | 0.023 | 0.004 | 0.092 |
| *Deltocephalinae 1* | Cicadellidae | 0 | 1 | 0 | 0 | 0 | 0.419 | 0.154 | 0.175 | 0.761 | 0.043 | 0.019 | 0.018 | 0.088 |
| *Deltocephalinae 2* | Cicadellidae | 0 | 1 | 0 | 0 | 0 | 0.168 | 0.096 | 0.047 | 0.404 | 0.050 | 0.023 | 0.016 | 0.105 |
| *Deltocephalinae 3* | Cicadellidae | 0 | 1 | 0 | 0 | 0 | 0.111 | 0.080 | 0.018 | 0.328 | 0.024 | 0.017 | 0.005 | 0.072 |
| *Deltocephalinae 4* | Cicadellidae | 0 | 1 | 0 | 0 | 0 | 0.105 | 0.089 | 0.016 | 0.369 | 0.029 | 0.023 | 0.005 | 0.092 |
| *Deltocephalinae 5* | Cicadellidae | 0 | 1 | 0 | 0 | 0 | 0.125 | 0.102 | 0.019 | 0.419 | 0.024 | 0.018 | 0.004 | 0.072 |
| *Dikraneurini 1* | Cicadellidae | 0 | 1 | 0 | 0 | 0 | 0.117 | 0.095 | 0.016 | 0.385 | 0.027 | 0.023 | 0.005 | 0.089 |
| *Erythroneurini 1* | Cicadellidae | 0 | 1 | 0 | 0 | 0 | 0.757 | 0.065 | 0.626 | 0.879 | 0.226 | 0.021 | 0.187 | 0.269 |
| *Eupelicini 1* | Cicadellidae | 0 | 1 | 0 | 0 | 0 | 0.080 | 0.059 | 0.013 | 0.248 | 0.045 | 0.026 | 0.011 | 0.109 |
| *Eurymelinae 1* | Cicadellidae | 0 | 1 | 0 | 0 | 0 | 0.118 | 0.078 | 0.023 | 0.330 | 0.043 | 0.028 | 0.009 | 0.117 |
| *Eurymeloides sp. 1* | Cicadellidae | 0 | 1 | 0 | 0 | 0 | 0.085 | 0.061 | 0.015 | 0.252 | 0.050 | 0.044 | 0.009 | 0.175 |
| *Exitianus sp. 1* | Cicadellidae | 0 | 1 | 0 | 0 | 0 | 0.079 | 0.065 | 0.016 | 0.271 | 0.104 | 0.077 | 0.017 | 0.304 |
| *Horouta sp. 1* | Cicadellidae | 0 | 1 | 0 | 0 | 0 | 0.270 | 0.145 | 0.068 | 0.638 | 0.025 | 0.014 | 0.007 | 0.062 |
| *Horouta sp. 2* | Cicadellidae | 0 | 1 | 0 | 0 | 0 | 0.057 | 0.045 | 0.010 | 0.179 | 0.096 | 0.073 | 0.017 | 0.296 |
| *Iassini 1* | Cicadellidae | 0 | 1 | 0 | 0 | 0 | 0.186 | 0.117 | 0.045 | 0.507 | 0.044 | 0.032 | 0.010 | 0.126 |
| *Iassini 2* | Cicadellidae | 0 | 1 | 0 | 0 | 0 | 0.077 | 0.066 | 0.013 | 0.259 | 0.056 | 0.048 | 0.009 | 0.186 |
| *Iassini 3* | Cicadellidae | 0 | 1 | 0 | 0 | 0 | 0.045 | 0.031 | 0.008 | 0.123 | 0.131 | 0.087 | 0.025 | 0.353 |
| *Iassini 4* | Cicadellidae | 0 | 1 | 0 | 0 | 0 | 0.107 | 0.075 | 0.020 | 0.305 | 0.048 | 0.034 | 0.011 | 0.142 |
| *Idiocerinae 1* | Cicadellidae | 0 | 1 | 0 | 0 | 0 | 0.110 | 0.084 | 0.018 | 0.349 | 0.030 | 0.027 | 0.005 | 0.105 |
| *Idiocerinae 2* | Cicadellidae | 0 | 1 | 0 | 0 | 0 | 0.110 | 0.095 | 0.019 | 0.371 | 0.065 | 0.056 | 0.010 | 0.214 |
| *Idiocerinae 3* | Cicadellidae | 0 | 1 | 0 | 0 | 0 | 0.112 | 0.091 | 0.017 | 0.355 | 0.029 | 0.026 | 0.005 | 0.096 |
| *Limotettix sp. 1* | Cicadellidae | 0 | 1 | 0 | 0 | 0 | 0.112 | 0.083 | 0.019 | 0.325 | 0.025 | 0.019 | 0.005 | 0.075 |
| *Macropsini 1* | Cicadellidae | 0 | 1 | 0 | 0 | 0 | 0.085 | 0.070 | 0.014 | 0.284 | 0.052 | 0.045 | 0.008 | 0.173 |
| *Membracidae 1* | Membracidae | 0 | 1 | 0 | 0 | 0 | 0.103 | 0.088 | 0.017 | 0.362 | 0.039 | 0.028 | 0.008 | 0.116 |
| *Nesoclutha sp. 1* | Cicadellidae | 0 | 1 | 0 | 0 | 0 | 0.587 | 0.104 | 0.396 | 0.808 | 0.091 | 0.021 | 0.058 | 0.137 |
| *Nirvanini 1* | Cicadellidae | 0 | 1 | 0 | 0 | 0 | 0.120 | 0.096 | 0.020 | 0.389 | 0.024 | 0.019 | 0.004 | 0.075 |
| *Orosius sp. 1* | Cicadellidae | 0 | 1 | 0 | 0 | 0 | 0.349 | 0.135 | 0.132 | 0.652 | 0.031 | 0.015 | 0.012 | 0.069 |
| *Paradorydium sp. 1* | Cicadellidae | 0 | 1 | 0 | 0 | 0 | 0.137 | 0.106 | 0.021 | 0.455 | 0.020 | 0.014 | 0.004 | 0.058 |
| *Rosopaella sp. 1* | Cicadellidae | 0 | 1 | 0 | 0 | 0 | 0.079 | 0.053 | 0.014 | 0.212 | 0.047 | 0.035 | 0.009 | 0.142 |

| | | | | | | | | | | | | | | |
|---|---|---|---|---|---|---|---|---|---|---|---|---|---|---|
| *Tartessini 1* | Cicadellidae | 0 | 1 | 0 | 0 | 0 | 0.072 | 0.051 | 0.013 | 0.205 | 0.057 | 0.038 | 0.013 | 0.155 |
| *Thymbrini 1* | Cicadellidae | 0 | 1 | 0 | 0 | 0 | 0.314 | 0.145 | 0.103 | 0.643 | 0.022 | 0.010 | 0.008 | 0.047 |
| *Thymbrini 2* | Cicadellidae | 0 | 1 | 0 | 0 | 0 | 0.130 | 0.099 | 0.022 | 0.418 | 0.020 | 0.014 | 0.004 | 0.058 |
| *Zygina sp. 1* | Cicadellidae | 0 | 1 | 0 | 0 | 0 | 0.118 | 0.083 | 0.023 | 0.343 | 0.043 | 0.030 | 0.010 | 0.122 |
| **Parasitoid wasps [Hymenoptera: Apocrita (excl. Aculeata)]** | | | | | | | | | | | | | | |
| *Agaonidae 1* | Agaonidae | 1 | 1 | 0 | 0 | 0 | 0.154 | 0.099 | 0.038 | 0.415 | 0.040 | 0.024 | 0.010 | 0.101 |
| *Braconidae 1* | Braconidae | 0 | 0 | 1 | 1 | 0 | 0.273 | 0.147 | 0.074 | 0.652 | 0.031 | 0.018 | 0.009 | 0.077 |
| *Braconidae 2* | Braconidae | 0 | 0 | 1 | 1 | 0 | 0.451 | 0.145 | 0.200 | 0.744 | 0.035 | 0.015 | 0.015 | 0.071 |
| *Braconidae 3* | Braconidae | 0 | 0 | 1 | 1 | 0 | 0.158 | 0.106 | 0.027 | 0.420 | 0.025 | 0.017 | 0.006 | 0.068 |
| *Braconidae 4* | Braconidae | 0 | 0 | 1 | 1 | 0 | 0.347 | 0.148 | 0.120 | 0.681 | 0.038 | 0.018 | 0.014 | 0.081 |
| *Braconidae 5* | Braconidae | 0 | 0 | 1 | 1 | 0 | 0.151 | 0.117 | 0.029 | 0.524 | 0.034 | 0.028 | 0.006 | 0.111 |
| *Braconidae 6* | Braconidae | 0 | 0 | 1 | 1 | 0 | 0.113 | 0.088 | 0.018 | 0.348 | 0.026 | 0.021 | 0.005 | 0.083 |
| *Braconidae 7* | Braconidae | 0 | 0 | 1 | 1 | 0 | 0.096 | 0.074 | 0.017 | 0.294 | 0.031 | 0.027 | 0.005 | 0.102 |
| *Braconidae 8* | Braconidae | 0 | 0 | 1 | 1 | 0 | 0.127 | 0.103 | 0.019 | 0.399 | 0.025 | 0.020 | 0.004 | 0.080 |
| *Braconidae 9* | Braconidae | 0 | 0 | 1 | 1 | 0 | 0.189 | 0.101 | 0.053 | 0.427 | 0.026 | 0.015 | 0.007 | 0.064 |
| *Braconidae 10* | Braconidae | 0 | 0 | 1 | 1 | 0 | 0.130 | 0.092 | 0.027 | 0.380 | 0.037 | 0.031 | 0.006 | 0.119 |
| *Braconidae 11* | Braconidae | 0 | 0 | 1 | 1 | 0 | 0.118 | 0.089 | 0.018 | 0.359 | 0.026 | 0.021 | 0.005 | 0.083 |
| *Braconidae 12* | Braconidae | 0 | 0 | 1 | 1 | 0 | 0.116 | 0.093 | 0.018 | 0.366 | 0.024 | 0.019 | 0.005 | 0.075 |
| *Braconidae 13* | Braconidae | 0 | 0 | 1 | 1 | 0 | 0.176 | 0.123 | 0.033 | 0.504 | 0.023 | 0.015 | 0.005 | 0.062 |
| *Braconidae 14* | Braconidae | 0 | 0 | 1 | 1 | 0 | 0.132 | 0.093 | 0.024 | 0.387 | 0.020 | 0.014 | 0.004 | 0.056 |
| *Braconidae 15* | Braconidae | 0 | 0 | 1 | 1 | 0 | 0.161 | 0.098 | 0.035 | 0.408 | 0.022 | 0.014 | 0.006 | 0.059 |
| *Braconidae 16* | Braconidae | 0 | 0 | 1 | 1 | 0 | 0.179 | 0.116 | 0.036 | 0.480 | 0.023 | 0.016 | 0.005 | 0.061 |
| *Braconidae 17* | Braconidae | 0 | 0 | 1 | 1 | 0 | 0.142 | 0.094 | 0.026 | 0.370 | 0.029 | 0.020 | 0.007 | 0.081 |
| *Braconidae 18* | Braconidae | 0 | 0 | 1 | 1 | 0 | 0.176 | 0.136 | 0.032 | 0.569 | 0.027 | 0.020 | 0.005 | 0.079 |
| *Braconidae 19* | Braconidae | 0 | 0 | 1 | 1 | 0 | 0.195 | 0.123 | 0.042 | 0.519 | 0.030 | 0.020 | 0.007 | 0.083 |
| *Braconidae 20* | Braconidae | 0 | 0 | 1 | 1 | 0 | 0.317 | 0.146 | 0.104 | 0.672 | 0.035 | 0.019 | 0.011 | 0.082 |
| *Braconidae 21* | Braconidae | 0 | 0 | 1 | 1 | 0 | 0.282 | 0.129 | 0.092 | 0.580 | 0.027 | 0.014 | 0.009 | 0.063 |
| *Braconidae 22* | Braconidae | 0 | 0 | 1 | 1 | 0 | 0.115 | 0.088 | 0.020 | 0.371 | 0.025 | 0.019 | 0.005 | 0.072 |
| *Braconidae 23* | Braconidae | 0 | 0 | 1 | 1 | 0 | 0.096 | 0.077 | 0.013 | 0.303 | 0.031 | 0.026 | 0.005 | 0.099 |
| *Braconidae 24* | Braconidae | 0 | 0 | 1 | 1 | 0 | 0.123 | 0.102 | 0.020 | 0.408 | 0.026 | 0.022 | 0.005 | 0.081 |
| *Braconidae 25* | Braconidae | 0 | 0 | 1 | 1 | 0 | 0.120 | 0.093 | 0.019 | 0.353 | 0.027 | 0.023 | 0.005 | 0.088 |
| *Braconidae 26* | Braconidae | 0 | 0 | 1 | 1 | 0 | 0.121 | 0.102 | 0.017 | 0.411 | 0.028 | 0.023 | 0.005 | 0.092 |
| *Braconidae 27* | Braconidae | 0 | 0 | 1 | 1 | 0 | 0.149 | 0.090 | 0.034 | 0.372 | 0.042 | 0.021 | 0.013 | 0.096 |
| *Braconidae 28* | Braconidae | 0 | 0 | 1 | 1 | 0 | 0.114 | 0.078 | 0.020 | 0.314 | 0.027 | 0.022 | 0.005 | 0.089 |
| *Braconidae 29* | Braconidae | 0 | 0 | 1 | 1 | 0 | 0.105 | 0.086 | 0.017 | 0.348 | 0.030 | 0.025 | 0.005 | 0.100 |
| *Braconidae 30* | Braconidae | 0 | 0 | 1 | 1 | 0 | 0.127 | 0.096 | 0.021 | 0.375 | 0.020 | 0.014 | 0.004 | 0.057 |
| *Braconidae 31* | Braconidae | 0 | 0 | 1 | 1 | 0 | 0.168 | 0.113 | 0.032 | 0.467 | 0.024 | 0.016 | 0.005 | 0.067 |
| *Braconidae 33* | Braconidae | 0 | 0 | 1 | 1 | 0 | 0.152 | 0.104 | 0.030 | 0.428 | 0.025 | 0.017 | 0.006 | 0.068 |
| *Braconidae 34* | Braconidae | 0 | 0 | 1 | 1 | 0 | 0.165 | 0.102 | 0.035 | 0.423 | 0.022 | 0.014 | 0.005 | 0.059 |
| *Braconidae 35* | Braconidae | 0 | 0 | 1 | 1 | 0 | 0.123 | 0.098 | 0.019 | 0.381 | 0.026 | 0.021 | 0.005 | 0.078 |
| *Braconidae 36* | Braconidae | 0 | 0 | 1 | 1 | 0 | 0.417 | 0.162 | 0.155 | 0.764 | 0.034 | 0.017 | 0.012 | 0.076 |
| *Braconidae 38* | Braconidae | 0 | 0 | 1 | 1 | 0 | 0.114 | 0.092 | 0.019 | 0.353 | 0.027 | 0.023 | 0.005 | 0.093 |
| *Braconidae 39* | Braconidae | 0 | 0 | 1 | 1 | 0 | 0.112 | 0.080 | 0.019 | 0.330 | 0.028 | 0.024 | 0.005 | 0.092 |
| *Braconidae 40* | Braconidae | 0 | 0 | 1 | 1 | 0 | 0.109 | 0.073 | 0.021 | 0.306 | 0.027 | 0.021 | 0.005 | 0.085 |
| *Braconidae 41* | Braconidae | 0 | 0 | 1 | 1 | 0 | 0.126 | 0.101 | 0.018 | 0.400 | 0.025 | 0.020 | 0.005 | 0.076 |
| *Chalcididae 1* | Chalcididae | 0 | 0 | 1 | 1 | 0 | 0.109 | 0.080 | 0.019 | 0.323 | 0.027 | 0.022 | 0.005 | 0.081 |
| *Chalcididae 2* | Chalcididae | 0 | 0 | 1 | 1 | 0 | 0.435 | 0.163 | 0.167 | 0.784 | 0.030 | 0.013 | 0.012 | 0.061 |
| *Chalcididae 3* | Chalcididae | 0 | 0 | 1 | 1 | 0 | 0.142 | 0.095 | 0.028 | 0.376 | 0.032 | 0.025 | 0.007 | 0.097 |
| *Chalcididae 4* | Chalcididae | 0 | 0 | 1 | 1 | 0 | 0.150 | 0.105 | 0.027 | 0.438 | 0.029 | 0.021 | 0.006 | 0.087 |
| *Chalcididae 5* | Chalcididae | 0 | 0 | 1 | 1 | 0 | 0.148 | 0.106 | 0.030 | 0.416 | 0.034 | 0.026 | 0.007 | 0.108 |
| *Diapriidae 1* | Diapriidae | 0 | 0 | 1 | 1 | 0 | 0.281 | 0.131 | 0.099 | 0.615 | 0.044 | 0.022 | 0.014 | 0.098 |
| *Diapriidae 2* | Diapriidae | 0 | 0 | 1 | 1 | 0 | 0.163 | 0.098 | 0.034 | 0.392 | 0.026 | 0.018 | 0.006 | 0.076 |
| *Diapriidae 3* | Diapriidae | 0 | 0 | 1 | 1 | 0 | 0.107 | 0.082 | 0.016 | 0.327 | 0.029 | 0.024 | 0.005 | 0.094 |
| *Diapriidae 4* | Diapriidae | 0 | 0 | 1 | 1 | 0 | 0.612 | 0.119 | 0.385 | 0.847 | 0.060 | 0.015 | 0.036 | 0.095 |
| *Diapriidae 5* | Diapriidae | 0 | 0 | 1 | 1 | 0 | 0.101 | 0.075 | 0.018 | 0.313 | 0.030 | 0.025 | 0.005 | 0.094 |
| *Encyrtidae 1* | Encyrtidae | 0 | 0 | 1 | 1 | 0 | 0.164 | 0.110 | 0.033 | 0.466 | 0.024 | 0.016 | 0.006 | 0.067 |
| *Encyrtidae 2* | Encyrtidae | 0 | 0 | 1 | 1 | 0 | 0.130 | 0.103 | 0.020 | 0.393 | 0.022 | 0.016 | 0.004 | 0.062 |
| *Encyrtidae 3* | Encyrtidae | 0 | 0 | 1 | 1 | 0 | 0.125 | 0.102 | 0.022 | 0.420 | 0.025 | 0.020 | 0.004 | 0.080 |
| *Encyrtidae 5* | Encyrtidae | 0 | 0 | 1 | 1 | 0 | 0.099 | 0.078 | 0.016 | 0.334 | 0.031 | 0.027 | 0.005 | 0.101 |
| *Encyrtidae 6* | Encyrtidae | 0 | 0 | 1 | 1 | 0 | 0.123 | 0.099 | 0.017 | 0.404 | 0.028 | 0.024 | 0.005 | 0.090 |
| *Encyrtidae 7* | Encyrtidae | 0 | 0 | 1 | 1 | 0 | 0.120 | 0.109 | 0.018 | 0.440 | 0.029 | 0.025 | 0.005 | 0.099 |
| *Encyrtidae 8* | Encyrtidae | 0 | 0 | 1 | 1 | 0 | 0.145 | 0.116 | 0.026 | 0.471 | 0.034 | 0.027 | 0.006 | 0.107 |
| *Encyrtidae 9* | Encyrtidae | 0 | 0 | 1 | 1 | 0 | 0.111 | 0.084 | 0.019 | 0.344 | 0.024 | 0.018 | 0.005 | 0.072 |
| *Encyrtidae 10* | Encyrtidae | 0 | 0 | 1 | 1 | 0 | 0.107 | 0.076 | 0.020 | 0.307 | 0.024 | 0.018 | 0.005 | 0.073 |
| *Encyrtidae 11* | Encyrtidae | 0 | 0 | 1 | 1 | 0 | 0.107 | 0.080 | 0.016 | 0.313 | 0.028 | 0.023 | 0.005 | 0.087 |
| *Gasteruptiidae 1* | Gasteruptiidae | 0 | 0 | 1 | 1 | 0 | 0.121 | 0.085 | 0.022 | 0.353 | 0.021 | 0.015 | 0.005 | 0.059 |
| *Ichneumonidae 1* | Ichneumonidae | 1 | 1 | 1 | 1 | 0 | 0.111 | 0.089 | 0.017 | 0.327 | 0.026 | 0.020 | 0.005 | 0.080 |
| *Ichneumonidae 2* | Ichneumonidae | 1 | 1 | 1 | 1 | 0 | 0.083 | 0.062 | 0.015 | 0.247 | 0.043 | 0.033 | 0.008 | 0.131 |
| *Ichneumonidae 3* | Ichneumonidae | 1 | 1 | 1 | 1 | 0 | 0.163 | 0.108 | 0.029 | 0.434 | 0.027 | 0.020 | 0.006 | 0.078 |
| *Ichneumonidae 4* | Ichneumonidae | 1 | 1 | 1 | 1 | 0 | 0.211 | 0.121 | 0.054 | 0.507 | 0.034 | 0.021 | 0.009 | 0.086 |
| *Ichneumonidae 5* | Ichneumonidae | 1 | 1 | 1 | 1 | 0 | 0.117 | 0.081 | 0.020 | 0.324 | 0.026 | 0.021 | 0.005 | 0.083 |
| *Ichneumonidae 6* | Ichneumonidae | 1 | 1 | 1 | 1 | 0 | 0.154 | 0.100 | 0.028 | 0.403 | 0.027 | 0.021 | 0.006 | 0.081 |
| *Ichneumonidae 7* | Ichneumonidae | 1 | 1 | 1 | 1 | 0 | 0.108 | 0.079 | 0.016 | 0.323 | 0.029 | 0.025 | 0.005 | 0.097 |
| *Ichneumonidae 8* | Ichneumonidae | 1 | 1 | 1 | 1 | 0 | 0.145 | 0.102 | 0.028 | 0.421 | 0.034 | 0.030 | 0.007 | 0.106 |
| *Ichneumonidae 9* | Ichneumonidae | 1 | 1 | 1 | 1 | 0 | 0.123 | 0.092 | 0.022 | 0.375 | 0.022 | 0.016 | 0.004 | 0.065 |
| *Ichneumonidae 10* | Ichneumonidae | 1 | 1 | 1 | 1 | 0 | 0.167 | 0.112 | 0.030 | 0.474 | 0.026 | 0.018 | 0.005 | 0.072 |
| *Ichneumonidae 11* | Ichneumonidae | 1 | 1 | 1 | 1 | 0 | 0.176 | 0.112 | 0.033 | 0.456 | 0.024 | 0.016 | 0.005 | 0.064 |
| *Ichneumonidae 12* | Ichneumonidae | 1 | 1 | 1 | 1 | 0 | 0.114 | 0.080 | 0.019 | 0.321 | 0.023 | 0.016 | 0.005 | 0.064 |
| *Ichneumonidae 13* | Ichneumonidae | 1 | 1 | 1 | 1 | 0 | 0.118 | 0.097 | 0.019 | 0.381 | 0.028 | 0.024 | 0.005 | 0.095 |
| *Netelia producta* | Ichneumonidae | 1 | 1 | 1 | 1 | 0 | 0.333 | 0.133 | 0.135 | 0.669 | 0.051 | 0.021 | 0.020 | 0.101 |
| *Pteromalidae 1* | Pteromalidae | 1 | 0 | 1 | 1 | 0 | 0.119 | 0.079 | 0.025 | 0.329 | 0.041 | 0.027 | 0.010 | 0.112 |
| *Pteromalidae 2* | Pteromalidae | 1 | 0 | 1 | 1 | 0 | 0.214 | 0.122 | 0.058 | 0.526 | 0.048 | 0.034 | 0.011 | 0.141 |
| *Pteromalidae 3* | Pteromalidae | 1 | 0 | 1 | 1 | 0 | 0.113 | 0.091 | 0.019 | 0.364 | 0.028 | 0.023 | 0.005 | 0.092 |
| *Pteromalidae 4* | Pteromalidae | 1 | 0 | 1 | 1 | 0 | 0.175 | 0.125 | 0.032 | 0.516 | 0.028 | 0.022 | 0.005 | 0.088 |
| *Pteromalidae 5* | Pteromalidae | 1 | 0 | 1 | 1 | 0 | 0.103 | 0.078 | 0.017 | 0.315 | 0.029 | 0.024 | 0.005 | 0.092 |
| *Pteromalidae 6* | Pteromalidae | 1 | 0 | 1 | 1 | 0 | 0.154 | 0.124 | 0.025 | 0.495 | 0.034 | 0.030 | 0.006 | 0.112 |
| *Pteromalidae 7* | Pteromalidae | 1 | 0 | 1 | 1 | 0 | 0.290 | 0.143 | 0.090 | 0.643 | 0.035 | 0.020 | 0.011 | 0.087 |
| *Pteromalidae 8* | Pteromalidae | 1 | 0 | 1 | 1 | 0 | 0.164 | 0.104 | 0.031 | 0.419 | 0.027 | 0.019 | 0.006 | 0.079 |
| *Pteromalidae 9* | Pteromalidae | 1 | 0 | 1 | 1 | 0 | 0.121 | 0.084 | 0.023 | 0.346 | 0.021 | 0.015 | 0.005 | 0.061 |
| *Pteromalidae 10* | Pteromalidae | 1 | 0 | 1 | 1 | 0 | 0.157 | 0.103 | 0.032 | 0.423 | 0.026 | 0.017 | 0.006 | 0.070 |
| *Pteromalidae 12* | Pteromalidae | 1 | 0 | 1 | 1 | 0 | 0.121 | 0.091 | 0.019 | 0.365 | 0.024 | 0.018 | 0.004 | 0.071 |
| *Pteromalidae 13* | Pteromalidae | 1 | 0 | 1 | 1 | 0 | 0.117 | 0.083 | 0.020 | 0.328 | 0.024 | 0.018 | 0.005 | 0.075 |
| *Pteromalidae 14* | Pteromalidae | 1 | 0 | 1 | 1 | 0 | 0.136 | 0.099 | 0.021 | 0.399 | 0.020 | 0.014 | 0.004 | 0.058 |
| *Pteromalidae 15* | Pteromalidae | 1 | 0 | 1 | 1 | 0 | 0.244 | 0.137 | 0.064 | 0.592 | 0.028 | 0.017 | 0.008 | 0.070 |
| *Pteromalidae 16* | Pteromalidae | 1 | 0 | 1 | 1 | 0 | 0.191 | 0.118 | 0.044 | 0.510 | 0.030 | 0.020 | 0.007 | 0.085 |
| *Pteromalidae 17* | Pteromalidae | 1 | 0 | 1 | 1 | 0 | 0.089 | 0.044 | 0.027 | 0.195 | 0.082 | 0.031 | 0.035 | 0.154 |
| *Pteromalidae 18* | Pteromalidae | 1 | 0 | 1 | 1 | 0 | 0.143 | 0.123 | 0.024 | 0.533 | 0.037 | 0.032 | 0.006 | 0.127 |
| *Pteromalidae 19* | Pteromalidae | 1 | 0 | 1 | 1 | 0 | 0.153 | 0.100 | 0.033 | 0.415 | 0.026 | 0.017 | 0.006 | 0.071 |
| *Pteromalidae 20* | Pteromalidae | 1 | 0 | 1 | 1 | 0 | 0.081 | 0.065 | 0.013 | 0.267 | 0.049 | 0.038 | 0.009 | 0.151 |
| *Pteromalidae 21* | Pteromalidae | 1 | 0 | 1 | 1 | 0 | 0.120 | 0.094 | 0.016 | 0.371 | 0.026 | 0.021 | 0.005 | 0.083 |
| *Pteromalidae 22* | Pteromalidae | 1 | 0 | 1 | 1 | 0 | 0.081 | 0.058 | 0.014 | 0.231 | 0.047 | 0.037 | 0.009 | 0.150 |
| *Pteromalidae 23* | Pteromalidae | 1 | 0 | 1 | 1 | 0 | 0.123 | 0.099 | 0.018 | 0.387 | 0.028 | 0.027 | 0.005 | 0.099 |
| *Pteromalidae 24* | Pteromalidae | 1 | 0 | 1 | 1 | 0 | 0.116 | 0.088 | 0.018 | 0.368 | 0.028 | 0.024 | 0.005 | 0.093 |

| | | | | | | | | | | | | | | |
|---|---|---|---|---|---|---|---|---|---|---|---|---|---|---|
| *Pteromalidae 25* | Pteromalidae | 1 | 0 | 1 | 1 | 0 | 0.122 | 0.089 | 0.020 | 0.348 | 0.026 | 0.022 | 0.005 | 0.084 |
| *Pteromalidae 26* | Pteromalidae | 1 | 0 | 1 | 1 | 0 | 0.114 | 0.077 | 0.022 | 0.316 | 0.046 | 0.032 | 0.010 | 0.130 |
| *Pteromalidae 27* | Pteromalidae | 1 | 0 | 1 | 1 | 0 | 0.117 | 0.101 | 0.019 | 0.407 | 0.025 | 0.019 | 0.005 | 0.077 |
| *Pteromalidae 28* | Pteromalidae | 1 | 0 | 1 | 1 | 0 | 0.097 | 0.071 | 0.017 | 0.273 | 0.031 | 0.025 | 0.006 | 0.098 |
| *Pteromalidae 29* | Pteromalidae | 1 | 0 | 1 | 1 | 0 | 0.169 | 0.123 | 0.033 | 0.504 | 0.025 | 0.017 | 0.005 | 0.070 |
| *Pteromalidae 30* | Pteromalidae | 1 | 0 | 1 | 1 | 0 | 0.115 | 0.089 | 0.018 | 0.348 | 0.024 | 0.020 | 0.005 | 0.077 |
| *Pteromalidae 31* | Pteromalidae | 1 | 0 | 1 | 1 | 0 | 0.121 | 0.100 | 0.019 | 0.390 | 0.026 | 0.021 | 0.005 | 0.085 |
| *Pteromalidae 32* | Pteromalidae | 1 | 0 | 1 | 1 | 0 | 0.122 | 0.100 | 0.019 | 0.398 | 0.028 | 0.024 | 0.005 | 0.092 |
| *Pteromalidae 33* | Pteromalidae | 1 | 0 | 1 | 1 | 0 | 0.115 | 0.090 | 0.019 | 0.364 | 0.028 | 0.025 | 0.005 | 0.094 |
| *Pteromalidae 34* | Pteromalidae | 1 | 0 | 1 | 1 | 0 | 0.117 | 0.099 | 0.016 | 0.392 | 0.028 | 0.025 | 0.005 | 0.096 |
| *Scelionidae 1* | Scelionidae | 1 | 0 | 1 | 1 | 0 | 0.113 | 0.092 | 0.017 | 0.352 | 0.029 | 0.026 | 0.005 | 0.100 |
| *Scelionidae 2* | Scelionidae | 1 | 0 | 1 | 1 | 0 | 0.114 | 0.092 | 0.017 | 0.357 | 0.027 | 0.022 | 0.004 | 0.088 |
| *Scelionidae 3* | Scelionidae | 1 | 0 | 1 | 1 | 0 | 0.096 | 0.077 | 0.017 | 0.295 | 0.034 | 0.024 | 0.007 | 0.095 |
| *Scelionidae 4* | Scelionidae | 1 | 0 | 1 | 1 | 0 | 0.116 | 0.082 | 0.021 | 0.337 | 0.021 | 0.015 | 0.004 | 0.057 |
| *Scelionidae 5* | Scelionidae | 1 | 0 | 1 | 1 | 0 | 0.200 | 0.118 | 0.050 | 0.526 | 0.040 | 0.027 | 0.009 | 0.114 |
| *Scelionidae 6* | Scelionidae | 1 | 0 | 1 | 1 | 0 | 0.304 | 0.147 | 0.094 | 0.650 | 0.025 | 0.013 | 0.008 | 0.058 |
| *Scelionidae 7* | Scelionidae | 1 | 0 | 1 | 1 | 0 | 0.187 | 0.120 | 0.039 | 0.501 | 0.032 | 0.021 | 0.008 | 0.088 |
| *Scelionidae 8* | Scelionidae | 1 | 0 | 1 | 1 | 0 | 0.160 | 0.114 | 0.032 | 0.475 | 0.026 | 0.019 | 0.006 | 0.077 |
| *Scelionidae 9* | Scelionidae | 1 | 0 | 1 | 1 | 0 | 0.122 | 0.096 | 0.019 | 0.390 | 0.028 | 0.024 | 0.005 | 0.092 |
| *Scelionidae 10* | Scelionidae | 1 | 0 | 1 | 1 | 0 | 0.149 | 0.131 | 0.021 | 0.557 | 0.021 | 0.016 | 0.004 | 0.066 |
| *Scelionidae 11* | Scelionidae | 1 | 0 | 1 | 1 | 0 | 0.116 | 0.090 | 0.018 | 0.363 | 0.025 | 0.021 | 0.005 | 0.081 |
| *Scelionidae 12* | Scelionidae | 1 | 0 | 1 | 1 | 0 | 0.120 | 0.098 | 0.017 | 0.392 | 0.027 | 0.022 | 0.005 | 0.090 |
| *Torymidae 1* | Torymidae | 0 | 0 | 1 | 1 | 0 | 0.214 | 0.125 | 0.050 | 0.512 | 0.027 | 0.017 | 0.007 | 0.073 |
| **Planthoppers [Hemiptera: Auchenorrhyncha: Fulgoromorpha: Fulgoroidea]** | | | | | | | | | | | | | | |
| *Anzora unicolor* | Flatidae | 0 | 1 | 0 | 0 | 0 | 0.311 | 0.088 | 0.170 | 0.518 | 0.113 | 0.027 | 0.067 | 0.170 |
| *Cixiidae 1* | Cixiidae | 0 | 1 | 0 | 0 | 0 | 0.083 | 0.067 | 0.013 | 0.265 | 0.048 | 0.038 | 0.008 | 0.147 |
| *Cixiidae 2* | Cixiidae | 0 | 1 | 0 | 0 | 0 | 0.156 | 0.109 | 0.030 | 0.446 | 0.031 | 0.025 | 0.006 | 0.101 |
| *Delphacidae 1* | Delphacidae | 0 | 1 | 0 | 0 | 0 | 0.104 | 0.082 | 0.018 | 0.338 | 0.030 | 0.027 | 0.005 | 0.103 |
| *Delphacidae 2* | Delphacidae | 0 | 1 | 0 | 0 | 0 | 0.076 | 0.062 | 0.014 | 0.259 | 0.058 | 0.039 | 0.013 | 0.160 |
| *Delphacidae 3* | Delphacidae | 0 | 1 | 0 | 0 | 0 | 0.161 | 0.106 | 0.032 | 0.429 | 0.024 | 0.017 | 0.006 | 0.068 |
| *Fulgoridae 1* | Fulgoridae | 0 | 1 | 0 | 0 | 0 | 0.127 | 0.092 | 0.021 | 0.387 | 0.020 | 0.014 | 0.004 | 0.055 |
| *Fulgoridae 2* | Fulgoridae | 0 | 1 | 0 | 0 | 0 | 0.171 | 0.119 | 0.038 | 0.498 | 0.040 | 0.031 | 0.008 | 0.123 |
| *Fulgoridae 3* | Fulgoridae | 0 | 1 | 0 | 0 | 0 | 0.130 | 0.103 | 0.018 | 0.401 | 0.026 | 0.024 | 0.005 | 0.089 |
| *Scolypopa australis* | Ricaniidae | 0 | 1 | 0 | 0 | 0 | 0.447 | 0.130 | 0.234 | 0.744 | 0.061 | 0.020 | 0.030 | 0.107 |
| *Siphanta sp. 1* | Flatidae | 0 | 1 | 0 | 0 | 0 | 0.366 | 0.134 | 0.155 | 0.674 | 0.056 | 0.022 | 0.025 | 0.107 |
| **Sawflies [Hymenoptera: Sympytha]** | | | | | | | | | | | | | | |
| *Pergidae 1* | Pergidae | 0 | 1 | 0 | 0 | 0 | 0.119 | 0.099 | 0.018 | 0.410 | 0.025 | 0.019 | 0.005 | 0.079 |
| **Stinging wasps [Hymenoptera: Apocrita: Aculeata (excl. Apoidea and Formicidae)]** | | | | | | | | | | | | | | |
| *Vespula germanica ** | Vespidae | 1 | 1 | 1 | 0 | 1 | 0.227 | 0.142 | 0.050 | 0.588 | 0.028 | 0.019 | 0.006 | 0.078 |
| *Austroscolia sp. 1* | Scolidae | 1 | 0 | 1 | 1 | 0 | 0.108 | 0.086 | 0.018 | 0.344 | 0.025 | 0.019 | 0.005 | 0.075 |
| *Bethylidae 1* | Bethylidae | 0 | 0 | 1 | 1 | 0 | 0.446 | 0.137 | 0.223 | 0.742 | 0.038 | 0.015 | 0.017 | 0.073 |
| *Bethylidae 2* | Bethylidae | 0 | 0 | 1 | 1 | 0 | 0.255 | 0.133 | 0.068 | 0.580 | 0.025 | 0.014 | 0.007 | 0.061 |
| *Bethylidae 3* | Bethylidae | 0 | 0 | 1 | 1 | 0 | 0.277 | 0.137 | 0.080 | 0.590 | 0.028 | 0.015 | 0.009 | 0.065 |
| *Bethylidae 4* | Bethylidae | 0 | 0 | 1 | 1 | 0 | 0.135 | 0.108 | 0.021 | 0.449 | 0.021 | 0.015 | 0.004 | 0.060 |
| *Bethylidae 5* | Bethylidae | 0 | 0 | 1 | 1 | 0 | 0.378 | 0.154 | 0.148 | 0.737 | 0.038 | 0.018 | 0.013 | 0.082 |
| *Bethylidae 6* | Bethylidae | 0 | 0 | 1 | 1 | 0 | 0.133 | 0.082 | 0.031 | 0.338 | 0.052 | 0.035 | 0.012 | 0.150 |
| *Bethylidae 7* | Bethylidae | 0 | 0 | 1 | 1 | 0 | 0.186 | 0.115 | 0.042 | 0.473 | 0.033 | 0.023 | 0.008 | 0.094 |
| *Mutillidae 1* | Mutillidae | 1 | 1 | 1 | 1 | 0 | 0.157 | 0.095 | 0.033 | 0.379 | 0.025 | 0.016 | 0.006 | 0.068 |
| *Mutillidae 2* | Mutillidae | 1 | 1 | 1 | 1 | 0 | 0.106 | 0.079 | 0.018 | 0.314 | 0.029 | 0.025 | 0.005 | 0.094 |
| *Mutillidae 3* | Mutillidae | 1 | 1 | 1 | 1 | 0 | 0.129 | 0.087 | 0.021 | 0.350 | 0.022 | 0.017 | 0.004 | 0.065 |
| *Mutillidae 4* | Mutillidae | 1 | 1 | 1 | 1 | 0 | 0.166 | 0.105 | 0.032 | 0.421 | 0.023 | 0.015 | 0.005 | 0.061 |
| *Mutillidae 5* | Mutillidae | 1 | 1 | 1 | 1 | 0 | 0.142 | 0.092 | 0.031 | 0.391 | 0.031 | 0.019 | 0.008 | 0.078 |
| *Mutillidae 6* | Mutillidae | 1 | 1 | 1 | 1 | 0 | 0.101 | 0.072 | 0.016 | 0.287 | 0.035 | 0.024 | 0.008 | 0.101 |
| *Mutillidae 7* | Mutillidae | 1 | 1 | 1 | 1 | 0 | 0.130 | 0.118 | 0.019 | 0.488 | 0.024 | 0.018 | 0.004 | 0.074 |
| *Pompilidae 1* | Pompilidae | 1 | 1 | 1 | 1 | 0 | 0.195 | 0.109 | 0.054 | 0.476 | 0.040 | 0.020 | 0.013 | 0.089 |
| *Pompilidae 2* | Pompilidae | 1 | 1 | 1 | 1 | 0 | 0.117 | 0.095 | 0.020 | 0.362 | 0.023 | 0.017 | 0.004 | 0.071 |
| *Scoliidae 1* | Scoliidae | 1 | 0 | 1 | 1 | 0 | 0.106 | 0.085 | 0.015 | 0.345 | 0.030 | 0.027 | 0.005 | 0.101 |
| *Tiphiidae 1* | Tiphiidae | 1 | 1 | 1 | 1 | 0 | 0.110 | 0.081 | 0.019 | 0.327 | 0.029 | 0.023 | 0.005 | 0.091 |
| *Tiphiidae 2* | Tiphiidae | 1 | 1 | 1 | 1 | 0 | 0.224 | 0.121 | 0.060 | 0.512 | 0.031 | 0.016 | 0.010 | 0.070 |

**Table S2.** Posterior estimates for the species richness of introduced and indigenous insects as estimated under the multi-species community model.

| | Species richness | | | |
|---|---|---|---|---|
| | Mean | SD | 2.50% | 97.50% |
| Introduced | 0.96 | 0.04 | 0.90 | 1.05 |
| Indigenous | 18.61 | 0.18 | 18.26 | 18.97 |
| All | 19.57 | 0.18 | 19.22 | 19.93 |

**Table S3.** The 133 plant species surveyed in the study, their plant origin, planting design element and growth form stratifications, and the posterior estimates for their associated species richness of indigenous species as estimated under the multi-species community model.

| Plant origin \| Planting design element \| Growth Form \| Species | Family | Introduced insect species richness | | | | Indigenous insect species richness | | | |
|---|---|---|---|---|---|---|---|---|---|
| | | Mean | SD | 2.50% | 97.50% | Mean | SD | 2.50% | 97.50% |
| **Nonnative** | | | | | | | | | |
| **Lawn** | | | | | | | | | |
| *Lawn complex 1* | Poaceae+ | 1.00 | 0.00 | 1.00 | 1.00 | 3.37 | 0.58 | 3.00 | 5.00 |
| *Lawn complex 2* | Poaceae+ | 0.00 | 0.00 | 0.00 | 0.00 | 2.31 | 0.46 | 2.00 | 3.00 |
| *Lawn complex 3* | Poaceae+ | 0.00 | 0.00 | 0.00 | 0.00 | 1.00 | 0.00 | 1.00 | 1.00 |
| *Lawn complex 4* | Poaceae+ | 0.00 | 0.00 | 0.00 | 0.00 | 11.55 | 1.09 | 10.00 | 14.00 |
| *Lawn complex 5* | Poaceae+ | 0.00 | 0.00 | 0.00 | 0.00 | 8.85 | 0.85 | 8.00 | 11.00 |
| *Lawn complex 6* | Poaceae+ | 0.15 | 0.35 | 0.00 | 1.00 | 9.31 | 0.91 | 8.00 | 11.00 |
| *Lawn complex 7* | Poaceae+ | 1.00 | 0.00 | 1.00 | 1.00 | 19.64 | 1.53 | 17.00 | 23.00 |
| *Lawn complex 8* | Poaceae+ | 0.23 | 0.42 | 0.00 | 1.00 | 15.61 | 1.17 | 14.00 | 18.00 |
| *Lawn complex 9* | Poaceae+ | 0.00 | 0.00 | 0.00 | 0.00 | 9.21 | 0.98 | 8.00 | 11.00 |
| *Lawn complex 10* | Poaceae+ | 2.00 | 0.00 | 2.00 | 2.00 | 5.94 | 0.87 | 5.00 | 8.00 |
| *Lawn complex 11* | Poaceae+ | 0.00 | 0.00 | 0.00 | 0.00 | 11.04 | 0.96 | 10.00 | 13.00 |
| *Lawn complex 12* | Poaceae+ | 1.00 | 0.00 | 1.00 | 1.00 | 12.38 | 1.29 | 10.00 | 15.00 |
| *Lawn complex 13* | Poaceae+ | 1.00 | 0.00 | 1.00 | 1.00 | 13.31 | 1.27 | 11.00 | 16.00 |
| *Lawn complex 14* | Poaceae+ | 0.00 | 0.00 | 0.00 | 0.00 | 7.99 | 0.88 | 7.00 | 10.00 |
| *Lawn complex 15* | Poaceae+ | 0.00 | 0.00 | 0.00 | 0.00 | 11.36 | 1.06 | 10.00 | 14.00 |
| *Lawn complex 16* | Poaceae+ | 1.00 | 0.00 | 1.00 | 1.00 | 13.93 | 1.25 | 12.00 | 17.00 |
| *Lawn complex 17* | Poaceae+ | 0.00 | 0.00 | 0.00 | 0.00 | 17.50 | 1.39 | 15.00 | 20.00 |
| *Lawn complex 18* | Poaceae+ | 0.00 | 0.00 | 0.00 | 0.00 | 10.74 | 1.12 | 9.00 | 13.00 |
| *Lawn complex 19* | Poaceae+ | 1.00 | 0.00 | 1.00 | 1.00 | 4.53 | 0.67 | 4.00 | 6.00 |
| *Lawn complex 20* | Poaceae+ | 1.00 | 0.00 | 1.00 | 1.00 | 7.08 | 0.91 | 6.00 | 9.00 |
| *Lawn complex 21* | Poaceae+ | 0.00 | 0.00 | 0.00 | 0.00 | 8.23 | 0.93 | 7.00 | 10.00 |
| *Lawn complex 22* | Poaceae+ | 0.00 | 0.00 | 0.00 | 0.00 | 5.51 | 0.66 | 5.00 | 7.00 |
| *Lawn complex 23* | Poaceae+ | 2.36 | 0.54 | 2.00 | 4.00 | 9.17 | 0.97 | 8.00 | 11.00 |
| *Lawn complex 24* | Poaceae+ | 0.00 | 0.00 | 0.00 | 0.00 | 13.77 | 1.21 | 12.00 | 16.00 |
| *Lawn complex 25* | Poaceae+ | 0.14 | 0.35 | 0.00 | 1.00 | 17.75 | 1.25 | 16.00 | 20.00 |
| *Lawn complex 26* | Poaceae+ | 0.25 | 0.43 | 0.00 | 1.00 | 8.75 | 0.80 | 8.00 | 11.00 |
| *Lawn complex 27* | Poaceae+ | 2.00 | 0.00 | 2.00 | 2.00 | 8.09 | 0.96 | 7.00 | 10.00 |
| *Lawn complex 28* | Poaceae+ | 0.00 | 0.00 | 0.00 | 0.00 | 5.70 | 0.74 | 5.00 | 7.00 |
| *Lawn complex 29* | Poaceae+ | 0.00 | 0.00 | 0.00 | 0.00 | 7.22 | 0.91 | 6.00 | 9.00 |
| *Lawn complex 30* | Poaceae+ | 1.20 | 0.40 | 1.00 | 2.00 | 14.98 | 1.27 | 13.00 | 18.00 |
| *Lawn complex 31* | Poaceae+ | 1.00 | 0.00 | 1.00 | 1.00 | 11.53 | 1.13 | 10.00 | 14.00 |
| *Lawn complex 32* | Poaceae+ | 1.36 | 0.48 | 1.00 | 2.00 | 1.00 | 0.00 | 1.00 | 1.00 |
| *Lawn complex 33* | Poaceae+ | 0.00 | 0.00 | 0.00 | 0.00 | 3.33 | 0.52 | 3.00 | 5.00 |
| *Lawn complex 34* | Poaceae+ | 0.16 | 0.37 | 0.00 | 1.00 | 14.13 | 1.20 | 12.00 | 17.00 |
| *Lawn complex 35* | Poaceae+ | 1.00 | 0.00 | 1.00 | 1.00 | 11.32 | 1.18 | 10.00 | 14.00 |
| *Lawn complex 36* | Poaceae+ | 0.00 | 0.00 | 0.00 | 0.00 | 6.96 | 0.84 | 6.00 | 9.00 |
| *Lawn complex 37* | Poaceae+ | 1.15 | 0.36 | 1.00 | 2.00 | 4.40 | 0.58 | 4.00 | 6.00 |
| *Lawn complex 38* | Poaceae+ | 1.00 | 0.00 | 1.00 | 1.00 | 3.35 | 0.55 | 3.00 | 5.00 |
| *Lawn complex 39* | Poaceae+ | 0.16 | 0.36 | 0.00 | 1.00 | 3.04 | 0.19 | 3.00 | 4.00 |
| *Lawn complex 40* | Poaceae+ | 1.00 | 0.00 | 1.00 | 1.00 | 4.51 | 0.65 | 4.00 | 6.00 |
| *Lawn complex 41* | Poaceae+ | 0.00 | 0.00 | 0.00 | 0.00 | 9.09 | 0.92 | 8.00 | 11.00 |
| **Midstorey \| Forb** | | | | | | | | | |
| *Acanthus mollis* | Acanthaceae | 1.00 | 0.00 | 1.00 | 1.00 | 19.17 | 1.46 | 17.00 | 22.00 |
| *Argyranthemum sp.* | Asteraceae | 2.00 | 0.00 | 2.00 | 2.00 | 22.98 | 1.57 | 20.00 | 26.00 |
| *Mentha pulegium* | Lamiaceae | 1.00 | 0.00 | 1.00 | 1.00 | 10.60 | 1.08 | 9.00 | 13.00 |
| *Salvia leucantha* | Lamiaceae | 3.00 | 0.00 | 3.00 | 3.00 | 24.72 | 1.67 | 22.00 | 28.00 |
| *Santolina chamaecyparissus* | Asteraceae | 0.00 | 0.00 | 0.00 | 0.00 | 7.88 | 0.84 | 7.00 | 10.00 |
| *Stachys byzantina* | Lamiaceae | 2.15 | 0.36 | 2.00 | 3.00 | 12.36 | 1.09 | 11.00 | 15.00 |
| *Tanacetum vulgare* | Asteraceae | 1.00 | 0.00 | 1.00 | 1.00 | 15.10 | 1.34 | 13.00 | 18.00 |
| **Midstorey \| Lilioid** | | | | | | | | | |
| *Agapanthus praecox* | Alliaceae | 3.32 | 0.52 | 3.00 | 5.00 | 21.18 | 1.38 | 19.00 | 24.00 |
| *Asparagus aethiopicus* | Asparagaceae | 2.21 | 0.41 | 2.00 | 3.00 | 11.69 | 1.16 | 10.00 | 14.00 |
| *Canna generalis* | Cannaceae | 2.00 | 0.00 | 2.00 | 2.00 | 13.38 | 1.11 | 12.00 | 16.00 |
| *Canna indica* | Cannaceae | 1.00 | 0.00 | 1.00 | 1.00 | 10.73 | 1.12 | 9.00 | 13.00 |
| *Chlorophytum comosum* | Phormiaceae | 0.23 | 0.42 | 0.00 | 1.00 | 6.72 | 0.76 | 6.00 | 8.00 |
| *Clivia miniata* | Amaryllidaceae | 0.13 | 0.34 | 0.00 | 1.00 | 16.90 | 1.25 | 15.00 | 20.00 |
| *Dietes sp.* | Iridaceae | 1.33 | 0.52 | 1.00 | 3.00 | 19.73 | 1.24 | 18.00 | 22.00 |
| *Hedychium sp* | Zingiberaceae | 0.00 | 0.00 | 0.00 | 0.00 | 7.95 | 0.86 | 7.00 | 10.00 |
| *Iris albicans* | Iridaceae | 0.00 | 0.00 | 0.00 | 0.00 | 3.23 | 0.46 | 3.00 | 4.00 |
| *Kniphofia sp.* | Xanthorrhoeaceae | 0.00 | 0.00 | 0.00 | 0.00 | 6.81 | 0.81 | 6.00 | 9.00 |
| *Strelitzia reginae* | Strelitziaceae | 0.00 | 0.00 | 0.00 | 0.00 | 8.77 | 0.84 | 8.00 | 11.00 |
| **Midstorey \| Graminoid** | | | | | | | | | |
| *Avena barbata* | Poaceae | 0.00 | 0.00 | 0.00 | 0.00 | 16.00 | 1.24 | 14.00 | 19.00 |
| *Carex flagellifera* | Cyperaceae | 0.45 | 0.58 | 0.00 | 2.00 | 9.84 | 0.84 | 9.00 | 12.00 |
| *Miscanthus sinensis* | Poaceae | 1.00 | 0.00 | 1.00 | 1.00 | 10.66 | 1.12 | 9.00 | 13.00 |
| **Midstorey \| Shrub** | | | | | | | | | |
| *Abelia grandiflora* | Caprifoliaceae | 1.00 | 0.00 | 1.00 | 1.00 | 6.57 | 0.72 | 6.00 | 8.00 |
| *Aloysia citrodora* | Verbenaceae | 0.00 | 0.00 | 0.00 | 0.00 | 8.06 | 0.92 | 7.00 | 10.00 |
| *Artemisia arborescens* | Asteraceae | 0.16 | 0.36 | 0.00 | 1.00 | 26.80 | 1.53 | 24.00 | 30.00 |
| *Aucuba japonica* | Garryaceae | 1.00 | 0.00 | 1.00 | 1.00 | 11.24 | 1.02 | 10.00 | 14.00 |
| *Buxus sp.* | Buxaceae | 0.00 | 0.00 | 0.00 | 0.00 | 29.61 | 1.81 | 27.00 | 34.00 |
| *Camellia sp.* | Theaceae | 0.22 | 0.42 | 0.00 | 1.00 | 11.05 | 1.06 | 10.00 | 14.00 |
| *Cercis canadensis* | Fabaceae | 0.00 | 0.00 | 0.00 | 0.00 | 4.48 | 0.63 | 4.00 | 6.00 |
| *Cistus sp.* | Cistaceae | 2.00 | 0.00 | 2.00 | 2.00 | 6.14 | 0.93 | 5.00 | 8.00 |
| *Echium candicans* | Boroginaceae | 4.00 | 0.00 | 4.00 | 4.00 | 12.01 | 1.25 | 10.00 | 15.00 |
| *Erythrina herbacea* | Fabaceae | 0.00 | 0.00 | 0.00 | 0.00 | 2.33 | 0.47 | 2.00 | 3.00 |
| *Euphorbia characias* | Euphorbiaceae | 0.31 | 0.46 | 0.00 | 1.00 | 12.89 | 1.16 | 11.00 | 15.00 |
| *Helleborus sp.* | Ranunculaceae | 1.00 | 0.00 | 1.00 | 1.00 | 10.50 | 1.11 | 9.00 | 13.00 |
| *Hydrangea sp.* | Hydrangeaceae | 0.00 | 0.00 | 0.00 | 0.00 | 12.74 | 1.16 | 11.00 | 15.00 |
| *Lavandula sp.* | Lamiaceae | 2.13 | 0.33 | 2.00 | 3.00 | 13.97 | 1.24 | 12.00 | 17.00 |
| *Nandina domestica* | Berberidaceae | 2.00 | 0.00 | 2.00 | 2.00 | 33.97 | 1.64 | 31.00 | 38.00 |
| *Nerium oleander* | Apocynaceae | 0.18 | 0.38 | 0.00 | 1.00 | 33.10 | 1.97 | 30.00 | 37.00 |
| *Pittosporum sp.* | Pittosporaceae | 1.00 | 0.00 | 1.00 | 1.00 | 11.56 | 1.15 | 10.00 | 14.00 |
| *Rosmarinus officinalis* | Lamiaceae | 4.12 | 0.32 | 4.00 | 5.00 | 23.16 | 1.62 | 20.00 | 27.00 |
| *Trachelospermum jasminoides* | Apocynaceae | 1.14 | 0.35 | 1.00 | 2.00 | 19.19 | 1.42 | 17.00 | 22.00 |
| *Viburnum sp.* | Adoxaceae | 0.15 | 0.35 | 0.00 | 1.00 | 29.33 | 1.83 | 26.00 | 33.00 |
| **Tree canopy** | | | | | | | | | |
| *Calodendrum capense* | Rutaceae | 1.16 | 0.37 | 1.00 | 2.00 | 4.69 | 0.70 | 4.00 | 6.00 |

| | | | | | | | | | |
|---|---|---|---|---|---|---|---|---|---|
| *Celtis australis* | Cannabaceae | 0.00 | 0.00 | 0.00 | 0.00 | 12.65 | 1.12 | 11.00 | 15.00 |
| *Fraxinus angustifolia* | Oleaceae | 0.00 | 0.00 | 0.00 | 0.00 | 11.01 | 0.94 | 10.00 | 13.00 |
| *Olea europaea* | Oleaceae | 0.00 | 0.00 | 0.00 | 0.00 | 7.90 | 0.84 | 7.00 | 10.00 |
| *Platanus acerifolia* | Platanaceae | 0.00 | 0.00 | 0.00 | 0.00 | 14.78 | 1.21 | 13.00 | 17.00 |
| *Pyrus calleryana* | Rosaceae | 0.00 | 0.00 | 0.00 | 0.00 | 1.00 | 0.00 | 1.00 | 1.00 |
| *Quercus bicolor* | Fagaceae | 1.56 | 0.63 | 1.00 | 3.00 | 19.79 | 1.43 | 17.00 | 23.00 |
| *Quercus robur* | Fagaceae | 1.00 | 0.00 | 1.00 | 1.00 | 12.95 | 1.25 | 11.00 | 16.00 |
| *Schinus molle* | Anacardiaceae | 2.46 | 0.62 | 2.00 | 4.00 | 60.71 | 2.39 | 56.48 | 66.00 |
| *Tilia cordata* | Tiliaceae | 0.15 | 0.36 | 0.00 | 1.00 | 7.71 | 0.77 | 7.00 | 9.00 |
| *Ulmus procera* | Ulmaceae | 1.33 | 0.74 | 1.00 | 3.00 | 39.63 | 1.78 | 37.00 | 43.00 |
| **Native** | | | | | | | | | |
| **Midstorey \| Forb** | | | | | | | | | |
| *Plectranthus argentatus* | Lamiaceae | 4.12 | 0.33 | 4.00 | 5.00 | 21.29 | 1.48 | 19.00 | 25.00 |
| **Midstorey \| Shrub** | | | | | | | | | |
| *Cassinia arcuata* | Asteraceae | 0.00 | 0.00 | 0.00 | 0.00 | 25.21 | 1.73 | 22.00 | 29.00 |
| *Melaleuca nesophila* | Myrtaceae | 1.00 | 0.00 | 1.00 | 1.00 | 7.01 | 0.91 | 6.00 | 9.00 |
| *Melaleuca viminalis* | Myrtaceae | 2.00 | 0.00 | 2.00 | 2.00 | 41.19 | 2.10 | 37.48 | 46.00 |
| *Westringia fruticosa* | Lamiaceae | 0.00 | 0.00 | 0.00 | 0.00 | 7.91 | 0.86 | 7.00 | 10.00 |
| **Tree canopy** | | | | | | | | | |
| *Angophora costata* | Myrtaceae | 0.22 | 0.41 | 0.00 | 1.00 | 16.92 | 1.27 | 15.00 | 20.00 |
| *Ficus macrophylla* | Moraceae | 1.46 | 0.63 | 1.00 | 3.00 | 31.87 | 1.53 | 29.00 | 35.00 |
| *Lophostemon confertus* | Myrtaceae | 2.00 | 0.00 | 2.00 | 2.00 | 29.79 | 1.80 | 27.00 | 33.00 |
| *Melaleuca armillaris* | Myrtaceae | 0.31 | 0.46 | 0.00 | 1.00 | 29.28 | 1.81 | 26.00 | 33.00 |
| *Melaleuca styphelioides* | Myrtaceae | 0.00 | 0.00 | 0.00 | 0.00 | 20.62 | 1.45 | 18.00 | 24.00 |
| **Indigenous** | | | | | | | | | |
| **Midstorey \| Lilioid** | | | | | | | | | |
| *Dianella sp.* | Phormiaceae | 1.00 | 0.00 | 1.00 | 1.00 | 11.26 | 1.04 | 10.00 | 13.00 |
| *Lomandra longifolia* | Xanthorrhoeaceae | 3.00 | 0.00 | 3.00 | 3.00 | 46.74 | 2.14 | 43.00 | 51.00 |
| **Midstorey \| Graminoid** | | | | | | | | | |
| *Baumea sp.* | Cyperaceae | 0.00 | 0.00 | 0.00 | 0.00 | 9.12 | 0.95 | 8.00 | 11.00 |
| *Dichanthium sericeum* | Poaceae | 0.00 | 0.00 | 0.00 | 0.00 | 11.27 | 1.01 | 10.00 | 13.00 |
| *Poa labillardierei* | Poaceae | 1.54 | 0.69 | 1.00 | 3.00 | 109.18 | 2.69 | 104.00 | 115.00 |
| *Rytidosperma sp.* | Poaceae | 3.44 | 0.76 | 3.00 | 5.00 | 95.64 | 2.63 | 91.00 | 101.00 |
| *Themeda triandra* | Poaceae | 3.17 | 0.37 | 3.00 | 4.00 | 66.01 | 2.71 | 61.00 | 72.00 |
| **Midstorey \| Shrub** | | | | | | | | | |
| *Acacia acinacea* | Fabaceae | 2.12 | 0.32 | 2.00 | 3.00 | 57.56 | 2.40 | 53.00 | 63.00 |
| *Acacia cognata* | Fabaceae | 1.44 | 0.57 | 1.00 | 3.00 | 16.82 | 1.24 | 15.00 | 20.00 |
| *Acacia verniciflua* | Fabaceae | 0.00 | 0.00 | 0.00 | 0.00 | 34.51 | 1.92 | 31.00 | 39.00 |
| *Bursaria spinosa* | Pittosporaceae | 2.39 | 0.58 | 2.00 | 4.00 | 57.47 | 2.13 | 54.00 | 62.00 |
| *Correa glabra* | Rutaceae | 1.22 | 0.42 | 1.00 | 2.00 | 18.11 | 1.34 | 16.00 | 21.00 |
| *Correa reflexa* | Rutaceae | 0.00 | 0.00 | 0.00 | 0.00 | 24.46 | 1.63 | 22.00 | 28.00 |
| *Dillwynia sp.* | Fabaceae | 0.00 | 0.00 | 0.00 | 0.00 | 12.60 | 1.16 | 11.00 | 15.00 |
| *Duma florulenta* | Polygonaceae | 2.00 | 0.00 | 2.00 | 2.00 | 14.25 | 1.36 | 12.00 | 17.00 |
| *Enchylaena tomentosa* | Amaranthaceae | 1.36 | 0.54 | 1.00 | 3.00 | 38.02 | 1.88 | 35.00 | 42.00 |
| *Goodenia ovata* | Goodeniaceae | 2.00 | 0.00 | 2.00 | 2.00 | 47.96 | 2.11 | 44.00 | 52.00 |
| *Hakea sp.* | Proteaceae | 1.00 | 0.00 | 1.00 | 1.00 | 11.38 | 1.07 | 10.00 | 14.00 |
| *Kunzea leptospermoides* | Myrtaceae | 2.00 | 0.00 | 2.00 | 2.00 | 20.95 | 1.54 | 18.00 | 24.00 |
| *Ozothamnus ferrugineus* | Asteraceae | 1.00 | 0.00 | 1.00 | 1.00 | 14.72 | 1.33 | 12.00 | 17.00 |
| *Rhagodia parabolica* | Amaranthaceae | 0.81 | 1.10 | 0.00 | 3.00 | 87.42 | 2.43 | 83.00 | 93.00 |
| **Tree canopy** | | | | | | | | | |
| *Acacia implexa* | Fabaceae | 0.00 | 0.00 | 0.00 | 0.00 | 15.29 | 1.29 | 13.00 | 18.00 |
| *Acacia mearnsii* | Fabaceae | 3.15 | 0.35 | 3.00 | 4.00 | 58.94 | 2.50 | 54.00 | 64.00 |
| *Acacia melanoxylon* | Fabaceae | 2.00 | 0.00 | 2.00 | 2.00 | 15.42 | 1.45 | 13.00 | 19.00 |
| *Allocasuarina verticillata* | Casuarinaceae | 1.00 | 0.00 | 1.00 | 1.00 | 29.79 | 1.87 | 27.00 | 34.00 |
| *Corymbia maculata* | Myrtaceae | 1.51 | 0.65 | 1.00 | 3.00 | 63.02 | 2.53 | 59.00 | 68.00 |
| *Eucalyptus camaldulensis* | Myrtaceae | 0.00 | 0.00 | 0.00 | 0.00 | 29.79 | 2.13 | 26.00 | 34.52 |
| *Eucalyptus sideroxylon* | Myrtaceae | 3.13 | 0.91 | 2.00 | 5.00 | 46.02 | 1.95 | 43.00 | 50.00 |
| *Eucalyptus tricarpa* | Myrtaceae | 3.52 | 0.65 | 3.00 | 5.00 | 15.00 | 1.27 | 13.00 | 18.00 |
| *Melaleuca lanceolata* | Myrtaceae | 1.24 | 0.43 | 1.00 | 2.00 | 8.82 | 0.85 | 8.00 | 11.00 |

Notes: + The dominant plant species in the lawn complexes was always a member of the Poaceae family, typically kikuyu (*Pennisetum clandestinum*) or couch (*Cynodon dactylon*). A range of small, ruderal herbaceous plants were often also present, including species in the follwoing families: Asteraceae, Brassicaceae, Caryophyllaceae, Fabaceae, Geraniaceae, Malvaceae, Oxalidaceae, Plantaginaceae and Polygonaceae.

**Table S4.** Community-level posterior estimates as estimated under the multi-species community model.

| | Mean | SD | 2.50% | 97.50% |
|---|---|---|---|---|
| Probability of occurrence \| Introduced | 0.196 | 0.059 | 0.104 | 0.330 |
| Probability of occurrence \| Indigenous | 0.135 | 0.010 | 0.115 | 0.155 |
| Probability of detection | 0.029 | 0.002 | 0.026 | 0.034 |

**Table S5.** Planting design element, midstorey growth form and plant origin groups and the posterior estimates for their associated species richness of indigenous insect species as estimated under the multi-species community model.

| | Indigenous insect species richness | | | |
|---|---|---|---|---|
| | **Mean** | **SD** | **2.50%** | **97.50%** |
| **Planting design element** | | | | |
| Lawn | 8.99 | 0.16 | 8.68 | 9.29 |
| Midstorey | 22.44 | 0.25 | 21.97 | 22.94 |
| Tree canopy | 24.14 | 0.35 | 23.48 | 24.84 |
| **Midstorey growth form** | | | | |
| Forb | 16.76 | 0.49 | 15.88 | 17.75 |
| Lilioid | 14.24 | 0.34 | 13.62 | 14.92 |
| Graminoid | 40.97 | 0.70 | 39.63 | 42.38 |
| Shrub | 22.53 | 0.30 | 21.97 | 23.16 |
| **Plant origin** | | | | |
| Nonnative | 12.43 | 0.15 | 12.15 | 12.73 |
| Native | 23.11 | 0.51 | 22.20 | 24.20 |
| Indigenous | 36.25 | 0.42 | 35.43 | 37.10 |

**Table S6.** Planting design element by plant origin and midstorey growth form by plant origin groups and the posterior estimates for their associated species richness of indigenous insect species for the whole community and for each insect functional group as estimated under the multi-species community model.

| | Whole community | | | | Pollinators | | | | Herbivores | | | | Predators | | | | Parasitoids | | | | Detritivores | | | |
|---|---|---|---|---|---|---|---|---|---|---|---|---|---|---|---|---|---|---|---|---|---|---|---|---|
| | Mean | SD | 2.50% | 97.50% | Mean | SD | 2.50% | 97.50% | Mean | SD | 2.50% | 97.50% | Mean | SD | 2.50% | 97.50% | Mean | SD | 2.50% | 97.50% | Mean | SD | 2.50% | 97.50% |
| **Planting design element by plant origin** | | | | | | | | | | | | | | | | | | | | | | | | |
| Nonative \| Lawn | 8.99 | 0.16 | 8.68 | 9.29 | 3.48 | 0.14 | 3.23 | 3.75 | 5.41 | 0.10 | 5.22 | 5.61 | 3.32 | 0.12 | 3.08 | 3.57 | 2.05 | 0.10 | 1.88 | 2.25 | 3.35 | 0.12 | 3.13 | 3.61 |
| Nonative \| Midstorey | 14.52 | 0.22 | 14.10 | 14.95 | 5.35 | 0.22 | 4.95 | 5.79 | 7.32 | 0.12 | 7.10 | 7.56 | 5.10 | 0.17 | 4.74 | 5.44 | 2.87 | 0.15 | 2.57 | 3.17 | 5.14 | 0.17 | 4.82 | 5.49 |
| Nonnative \| Tree canopy | 17.53 | 0.40 | 16.73 | 18.36 | 5.66 | 0.30 | 5.10 | 6.30 | 7.01 | 0.22 | 6.64 | 7.45 | 10.53 | 0.44 | 9.70 | 11.40 | 6.61 | 0.46 | 5.75 | 7.50 | 10.56 | 0.45 | 9.70 | 11.50 |
| Native \| Midstorey | 23.90 | 0.82 | 22.50 | 25.50 | 4.62 | 0.47 | 3.67 | 5.67 | 11.16 | 0.50 | 10.33 | 12.33 | 4.02 | 0.42 | 3.33 | 5.00 | 1.00 | 0.00 | 1.00 | 1.00 | 5.88 | 0.50 | 5.00 | 7.00 |
| Native \| Tree canopy | 22.58 | 0.62 | 21.50 | 23.83 | 8.53 | 0.52 | 7.50 | 9.67 | 11.33 | 0.32 | 10.71 | 12.00 | 10.44 | 0.52 | 9.43 | 11.43 | 6.52 | 0.51 | 5.50 | 7.50 | 10.39 | 0.59 | 9.33 | 11.67 |
| Indigenous \| Midstorey | 38.35 | 0.49 | 37.43 | 39.38 | 13.20 | 0.54 | 12.19 | 14.33 | 18.36 | 0.24 | 17.95 | 18.86 | 15.71 | 0.49 | 14.67 | 16.62 | 10.27 | 0.58 | 9.29 | 11.41 | 15.84 | 0.53 | 14.90 | 17.00 |
| Indigenous \| Tree canopy | 31.34 | 0.64 | 30.11 | 32.67 | 7.31 | 0.42 | 6.50 | 8.13 | 16.34 | 0.37 | 15.67 | 17.11 | 14.10 | 0.59 | 13.00 | 15.22 | 7.06 | 0.47 | 6.22 | 8.00 | 14.29 | 0.60 | 13.22 | 15.56 |
| **Midstorey growth form by plant origin** | | | | | | | | | | | | | | | | | | | | | | | | |
| Nonative \| Forb | 16.12 | 0.52 | 15.14 | 17.14 | 5.73 | 0.39 | 5.00 | 6.57 | 8.62 | 0.30 | 8.14 | 9.29 | 5.20 | 0.32 | 4.57 | 5.86 | 2.07 | 0.22 | 1.75 | 2.50 | 5.20 | 0.34 | 4.57 | 5.86 |
| Native \| Forb | 21.29 | 1.48 | 19.00 | 25.00 | 5.11 | 0.90 | 4.00 | 7.00 | 14.45 | 1.14 | 13.00 | 17.00 | 5.03 | 0.88 | 4.00 | 7.00 | 0.00 | 0.00 | 0.00 | 0.00 | 5.11 | 1.10 | 4.00 | 7.00 |
| Nonnative \| Lilioid | 11.55 | 0.32 | 11.00 | 12.18 | 4.48 | 0.27 | 4.00 | 5.00 | 6.07 | 0.20 | 5.73 | 6.55 | 3.30 | 0.20 | 2.90 | 3.70 | 1.84 | 0.17 | 1.60 | 2.20 | 3.37 | 0.21 | 3.00 | 3.80 |
| Indigenous \| Lilioid | 29.00 | 1.19 | 27.00 | 31.50 | 12.35 | 1.00 | 10.50 | 14.50 | 14.90 | 0.66 | 14.00 | 16.50 | 9.96 | 0.90 | 8.50 | 12.00 | 4.45 | 0.56 | 3.50 | 5.50 | 10.25 | 0.99 | 8.50 | 12.50 |
| Nonative \| Graminoid | 12.17 | 0.63 | 11.00 | 13.33 | 6.52 | 0.55 | 5.67 | 7.67 | 7.33 | 0.44 | 6.67 | 8.33 | 4.41 | 0.43 | 3.67 | 5.33 | 3.06 | 0.43 | 2.50 | 4.00 | 4.49 | 0.46 | 3.67 | 5.33 |
| Indigenous \| Graminoid | 58.24 | 1.03 | 56.40 | 60.40 | 20.86 | 1.05 | 19.00 | 23.00 | 28.02 | 0.47 | 27.20 | 29.00 | 27.03 | 1.07 | 25.00 | 29.20 | 15.60 | 1.04 | 13.80 | 17.80 | 27.19 | 1.13 | 25.20 | 29.60 |
| Nonative \| Shrub | 15.93 | 0.31 | 15.35 | 16.55 | 5.54 | 0.28 | 5.06 | 6.11 | 7.55 | 0.17 | 7.25 | 7.90 | 6.11 | 0.25 | 5.63 | 6.58 | 3.53 | 0.23 | 3.08 | 4.00 | 6.15 | 0.24 | 5.68 | 6.63 |
| Native \| Shrub | 20.33 | 0.77 | 19.00 | 22.00 | 7.54 | 0.63 | 6.33 | 8.67 | 9.81 | 0.36 | 9.25 | 10.50 | 4.57 | 0.40 | 3.75 | 5.50 | 2.87 | 0.41 | 2.33 | 3.67 | 4.60 | 0.40 | 4.00 | 5.50 |
| Indigenous \| Shrub | 32.59 | 0.52 | 31.64 | 33.64 | 10.58 | 0.49 | 9.71 | 11.57 | 15.41 | 0.27 | 14.93 | 15.93 | 12.49 | 0.47 | 11.57 | 13.43 | 8.78 | 0.55 | 7.80 | 9.90 | 12.58 | 0.49 | 11.71 | 13.64 |
